# Metabolic enzymes moonlight as selective autophagy receptors to protect plants against viral-induced cellular damage

**DOI:** 10.1101/2024.05.06.590709

**Authors:** Marion Clavel, Anita Bianchi, Roksolana Kobylinska, Roan Groh, Juncai Ma, Ranjith K. Papareddy, Nenad Grujic, Lorenzo Picchianti, Ethan Stewart, Michael Schutzbier, Karel Stejskal, Juan Carlos de la Concepcion, Victor Sanchez de Medina Hernandez, Yoav Voichek, Pieter Clauw, Joanna Gunis, Gerhard Durnberger, Jens Christian Muelders, Annett Grimm, Arthur Sedivy, Mathieu Erhardt, Victoria Vyboishchikov, Peng Gao, Esther Lechner, Emilie Vantard, Jakub Jez, Elisabeth Roitinger, Pascal Genschik, Byung-Ho Kang, Yasin Dagdas

## Abstract

RNA viruses co-opt the host endomembrane system and organelles to build replication complexes for infection. How the host responds to these membrane perturbations is poorly understood. Here, we explore the autophagic response of *Arabidopsis thaliana* to three viruses that hijack different cellular compartments. Autophagy is significantly induced within systemically infected tissues, its disruption rendering plants highly sensitive to infection. Contrary to being an antiviral defense mechanism as previously suggested, quantitative analyses of the viral loads established autophagy as a tolerance pathway. Further analysis of one of these viruses, the Turnip Crinkle Virus (TCV) that hijack mitochondria, showed that despite perturbing mitochondrial integrity, TCV does not trigger a typical mitophagy response. Instead, TCV and Turnip yellow mosaic virus (TYMV) infection activates a distinct selective autophagy mechanism, where oligomeric metabolic enzymes moonlight as selective autophagy receptors and degrade key executors of defense and cell death such as EDS1. Altogether, our study reveals an autophagy-regulated metabolic rheostat that gauges cellular integrity during viral infection and degrades cell death executors to avoid catastrophic amplification of immune signaling.

## Introduction

Positive sense single stranded RNA (+ssRNA) viruses pose a serious threat to global food security and human health. They are responsible for some of the deadliest pandemics in history, including the recent COVID-19 outbreak, and continue to cause billions of dollars worth of crop losses every year ^1^. A universal feature of the +ssRNA virus infection cycle is its dependence on host endomembranes for viral genome replication. These viruses extensively remodel host endomembranes to build viral replication complexes (VRCs), wherein viral and host factors are recruited to create an optimal microenvironment for replication and possibly evade host immune pathways ^2,3^. Understanding how viruses hijack cellular compartments for infection, and how the host counteracts viral membrane remodeling is paramount for the development of novel strategies to mitigate the effects of viral infections. Three major mechanisms of modified-self or non-self recognition counteract viral infections: RNA interference, intracellular nucleotide-binding and leucine-rich repeat (NLR) immune receptors, and selective autophagy ^4–6^. Selective autophagy is mediated by selective autophagy receptors (SARs) − modular proteins that bind both the autophagy cargo and the core autophagy protein ATG8 to sequester the harmful substrates within the autophagosome ^7^. SAR-ATG8 interaction is mediated by the short linear ATG8 Interacting Motif (AIM) that is functionally conserved throughout eukaryotes, denoted as Θ_0_-X_1_-X_2_-Γ_3_, where Θ represents an aromatic residue (W/F/Y), Γ an aliphatic residue (L/V/I), and X any amino acid ^8^. Bound cargos are engulfed into the autophagosome, the biogenesis of which is coordinated by conserved ATG (autophagy related gene) proteins ^9^, and delivered to the lytic compartment for recycling. Despite numerous reports highlighting selective autophagy as a critical defense system against viruses ^10–15^, particularly in plants, the molecular mechanisms underlying viral-induced autophagy remain elusive.

## Results

### Autophagy safeguards host fitness during viral infection without exerting an antiviral role

To assess the role of autophagy in viral infection, we challenged Arabidopsis *atg2* and *atg5* mutants with three different single-stranded +ssRNA viruses. The chosen viruses all efficiently replicate and move within Arabidopsis, leading to persistent systemic infections with distinct symptoms. Each virus hijacks distinct organelles for replication: Turnip crinkle virus (TCV) colonizes mitochondria ^16^, Turnip yellow mosaic virus (TYMV) chloroplasts ^17^, and Turnip mosaic virus (TuMV, with reporter 6k2-scarlet protein) endoplasmic reticulum (ER) ^18^. Time-course analyses revealed more severe infection symptoms in autophagy-deficient plants across all viruses, either early (TCV and TuMV) or at a later time point (TYMV) (Figure 1A). In all mutants, necrotic lesions were apparent and not observed in infected wild-type Columbia-0 (Col-0) plants. This prompted us to quantitatively assess symptom progression in a wide range of autophagy mutants, using image-based high-throughput phenotyping. We included mutants of *nbr1*, a well-characterized aggrephagy receptor ^19^ and of *fmt*, a possible mitophagy receptor^20^. To account for a potential effect of antiviral RNAi on the observed symptoms, we also included *dcl2/dcl4/atg2* and *dcl2/dcl4/atg5* triple mutant plants. All *atg* mutants were validated by their sensitivity to nitrogen starvation (Figure S1A), lack of ATG8 lipidation and NBR1 recycling (Figure S1B). We obtained dense time series for 1134 plants (14 genotypes, 4 treatments, 9 replicates for mock and 18 replicates for each virus). Our analysis revealed that all canonical autophagy mutants had more severe infection symptoms than wild-type plants as scored by % disease symptoms relative to Col-0 (Figure 1B, Figure S1C, Supplemental videos 1 to 4). Both *dcl2/dcl4/atg2* and *dcl2/dcl4/atg5* mutants exhibited either *atg*-like symptoms (TuMV) or additive symptoms (TCV and TYMV), suggesting autophagy and RNAi act independently during viral infection (Figure 1B). Receptor mutants resembled wild-type plants in all experimental conditions (Figure 1B, Figure S1D, E). Based on these observations, we conclude that autophagy plays a critical role during +ssRNA virus infection in Arabidopsis.

**FIGURE 1:**
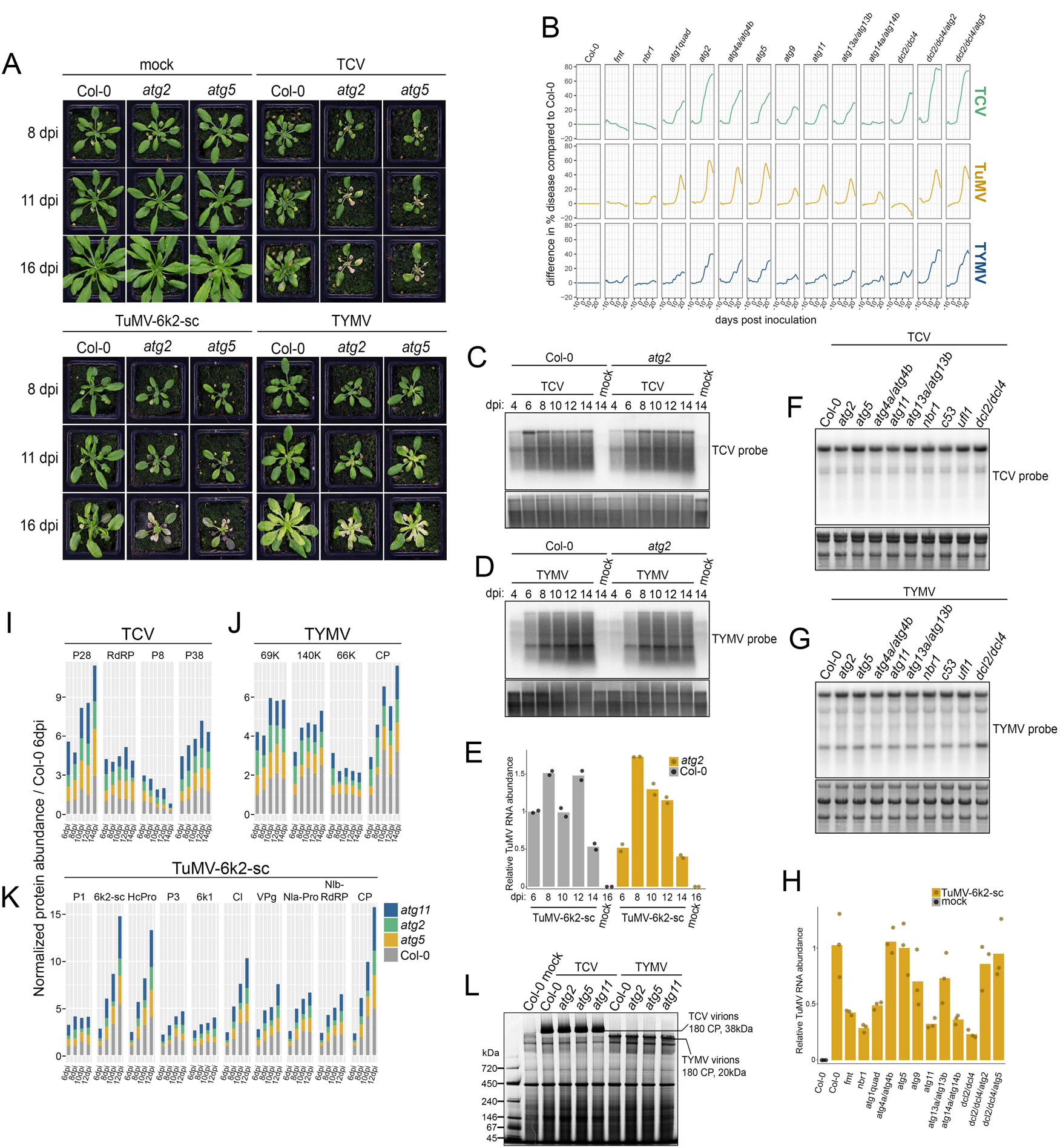
Autophagy safeguards host fitness during viral infection but does not degrade viral components. **(A)** Phenotypic characterization of *Arabidopsis* Col-0 (WT), *atg2* and *atg5* mutant plants after inoculation with buffer (mock), with TCV, TYMV and TuMV-6k2-sc, 8, 11 and 16 days-post-inoculation (dpi). **(B)** Phenotypic characterization of *Arabidopsis* Col-0 (WT), *fmt, nbr1, atg1* quadruple mutant*, atg2, atg4* double mutant*, atg5, atg9, atg11, atg13* double mutant*, atg14* double mutant*, dcl2/dcl4* double mutant*, dcl2/dcl4/atg2* and *dcl2/dcl4/atg5* triple mutant combination using automated camera phenotyping. % of disease is defined as plant area harboring symptomatic colors. The obtained value for Col-0 is subtracted to all mutants to obtain the difference in % disease compared to Col-0 from -10 days (pre inoculation) to + 28 days (post inoculation). **(C)** Northern blot of viral (+) RNA accumulation in Col-0 and *atg2* TCV-infected systemic leaves. 3-week-old rosettes were inoculated with buffer (mock) or with TCV and young systemic leaves were collected at the indicated time points (4 to 14 dpi). n=3 to 4 plants per sample-genotype combination. 10µg of total RNA was loaded per well and membrane was hybridized with a radiolabeled probe specific for the 3’ sequence of TCV. Membrane was stained with methylene blue to verify loading. **(D)** Northern blot of viral (+) RNA accumulation in Col-0 and *atg2* TYMV-infected systemic leaves. 3-week-old rosettes were inoculated with buffer (mock) or with TYMV and young systemic leaves were collected at the indicated time points (4 to 14 dpi). n=3 to 4 plants per sample-genotype combination. 10µg of total RNA was loaded per well and membrane was hybridized with a radiolabeled probe specific for the 3’ sequence of TYMV. Membrane was stained with methylene blue to verify loading. **(E)** TuMV-6k2-sc RNA accumulation in systemic leaves of Col-0 and *atg2* measured by RT-qPCR. 3-week-old rosettes were inoculated with buffer (mock) or with TuMV-6k2-sc and young systemic leaves were collected at the indicated time points (6 to 14 dpi, 16 dpi for mock samples). n=3 to 4 plants per sample-genotype combination. Individual technical replicates (n=2) are represented by dots and bar graph represent the average ΔΔCt value, expressed as relative to the Col-0 TuMV-6k2-sc 6 dpi sample. **(F)** Northern blot of viral (+) RNA accumulation in TCV-infected systemic leaves of the indicated genotypes. 3-week-old rosettes were inoculated with buffer (mock) or with TCV and young systemic leaves were collected at 10 dpi. n=3 to 4 plants per sample-genotype combination. 6µg of total RNA was loaded per well and membrane was hybridized with a radiolabeled probe specific for the 3’ sequence of TCV. Membrane was stained with methylene blue to verify loading. **(G)** Northern blot of viral (+) RNA accumulation in TYMV-infected systemic leaves of the indicated genotypes. 3-week-old rosettes were inoculated with buffer (mock) or with TYMV and young systemic leaves were collected at 10 dpi. n=3 to 4 plants per sample-genotype combination. 6µg of total RNA was loaded per well and membrane was hybridized with a radiolabeled probe specific for the 3’ sequence of TYMV. Membrane was stained with methylene blue to verify loading. **(H)** TuMV-6k2-sc RNA accumulation in systemic leaves of the indicated genotypes measured by RT-qPCR. 3-week-old rosettes were inoculated with buffer (mock) or with TuMV-6k2-sc and young systemic leaves were collected at 10 dpi. n=3 to 4 plants per sample-genotype combination. Individual technical replicates (n=3) are represented by dots and bar graph represent the average ΔΔCt value, expressed as relative to the Col-0 TuMV-6k2-sc sample. **(I)** Quantification of TCV translation products P28 (4 peptides), RdRP (4 peptides), P8 (3 peptides) and P38 (9 peptides) by parallel reaction monitoring (PRM). 3-week-old rosettes were inoculated with buffer (mock) or with TCV and young systemic leaves were collected at the indicated time points (6 to 14 dpi). n=4 to 8 plants per sample-genotype combination. Average peptide area was calculated for each protein and normalized to average peptide area for all control plant proteins (7 proteins, a total of 27 peptides). For each protein, values are expressed relative to the Col-0 6 dpi samples and displayed as stacked histograms for Col-0, *atg2*, *atg5* and *atg11*. **(J)** Quantification of TYMV translation products 69K (10 peptides), 140K (18 peptides), 66K (5 peptides) and CP (6 peptides) by parallel reaction monitoring (PRM). 3-week-old rosettes were inoculated with buffer (mock) or with TYMV and young systemic leaves were collected at the indicated time points (6 to 14 dpi). n=4 to 8 plants per sample-genotype combination. Average peptide area was calculated for each protein and normalized to average peptide area for all control plant proteins (7 proteins, a total of 23 peptides). For each protein, values are expressed relative to the Col-0 6 dpi samples and displayed as stacked histograms for Col-0, *atg2*, *atg5* and *atg11*. **(K)** Quantification of TuMV-6k2-sc translation products P1 (5 peptides), 6k2-scarlet (5 peptides), HcPro (6 peptides), P3 (5 peptides), 6k1 (2 peptides), CI (6 peptides), VPg (6 peptides), Nla-Pro (4 peptides), Nlb-RdRP (4 peptides) and CP (6 peptides) by parallel reaction monitoring (PRM). 3-week-old rosettes were inoculated with buffer (mock) or with TuMV-6k2-sc and young systemic leaves were collected at the indicated time points (6 to 12 dpi). n=4 to 8 plants per sample-genotype combination. Average peptide area was calculated for each protein and normalized to average peptide area for all control plant proteins (7 proteins, a total of 16 peptides). For each protein, values are expressed relative to the Col-0 6 dpi samples and displayed as stacked histograms for Col-0, *atg2*, *atg5* and *atg11*. **(L)** BN-PAGE analysis of TCV and TYMV particles from native total plant extract in mock and TCV inoculated systemic leaves of WT, *atg2*, *atg5* and *atg11* plants. 3-week-old rosettes were inoculated with buffer (mock) or with TCV and young systemic leaves were collected at 12 dpi. n=10 plants. Total soluble proteins were extracted at a fixed fresh weight to buffer volume ratio (1:2) and equal volume (10 µl) was loaded per well.

We then asked if autophagy operates as an antiviral or tolerance pathway. To test this, we first examined the impact of autophagy deficiency on virus production. Viral genome accumulation of all three viruses in systemic tissue showed no difference between Col-0 and *atg2* at all time points studied (Figure 1C, D and E). We observed a similar outcome for various autophagy mutants at 10 dpi, with only minor variations (Figure 1F, G, H and Figure S1F, G). We then measured viral protein abundance in autophagy mutants, using parallel reaction monitoring (PRM) proteomics. We could successfully quantify 4 out of 5 TCV proteins, all proteins and polyprotein cleavage products of TYMV and all TuMV polyprotein cleavage products (including the 6k2-scarlet fusion protein), except PIPO (Table S1). After normalization to plant peptides, no viral protein was more abundant in any of the autophagy mutants tested at any given time point when compared to Col-0 (Figure 1I, J and K and Tables S2 to S4). Finally, we assessed the ability of TCV and TYMV to form fully assembled virions in systemic tissues using BN (Blue Native)-PAGE (Figure 1L). Similar to viral genome and protein quantifications, there was no appreciable difference in *atg* mutants compared to wild-type samples. We therefore conclude that autophagy does not exert a broad antiviral effect, *i.e* by direct degradation of viral components, but rather acts as a tolerance mechanism and the striking symptoms that we observed are due to the failure to degrade damaged host components that arise during infection.

### Autophagy is induced by viral replication-associated organelle damage

Next, we sought to determine if viral infection induces autophagy. First, we measured protein levels of ATG8 reporters that are commonly used to assess autophagy. Monitoring the abundance of GFP-ATG8E and GFP-ATG8H, representative of their respective clades ^21^, along with the stable GFP fragment released after vacuolar delivery ^22^, we consistently observed an increase in cleavage product and/or full-length reporter abundance after viral spread to systemic tissues (Figure 2A). Consistently, GFP-ATG11 autophagic flux was also increased in infected samples (Figure 2B). For live cell imaging of infection, we established an assay where we TCV-inoculated the first pair of true leaves in young seedlings and observed systemic infection in roots (Figure 2C). We used the B2-GFP reporter − a protein with high affinity to double stranded RNA ^23,24^ to visualize viral replication sites and TMRE − a membrane potential-sensitive dye to visualize mitochondria. TCV infection induced clumping of mitochondria and re-localization of B2-GFP to these mitochondrial clusters (Figure 2D). Although the shape and size of the observed clumps varied greatly (Figure S2A, supplemental video 5), we interpret those as mitochondria-derived viral replication complexes (VRCs), housing TCV replication in roots ^16,25^. We also observed a reproducible increase in GFP-ATG8E-labelled puncta in cells containing VRCs (Figure 2E and 2F), indicative of autophagy induction. Since VRC-supporting mitochondria were positive for TMRE (Figure 2D and E), this suggested hijacked mitochondria retained their membrane potential. To test this, we treated mock and TCV-infected mito-GFP plants with p-trifluoro-methoxyphenylhydrazone (FCCP), a membrane-depolarizing drug, to independently monitor mitochondria localization and membrane potential. Live cell imaging analysis revealed that the vast majority of VRC-supporting mitochondria retained TMRE staining (Figure S2B), with colocalization of both signals being lost only in the presence of FCCP (Figure S2C). Altogether, these results suggest that TCV infection leads to mitochondria clumping and autophagy induction, but does not affect mitochondrial membrane potential.

**FIGURE 2:**
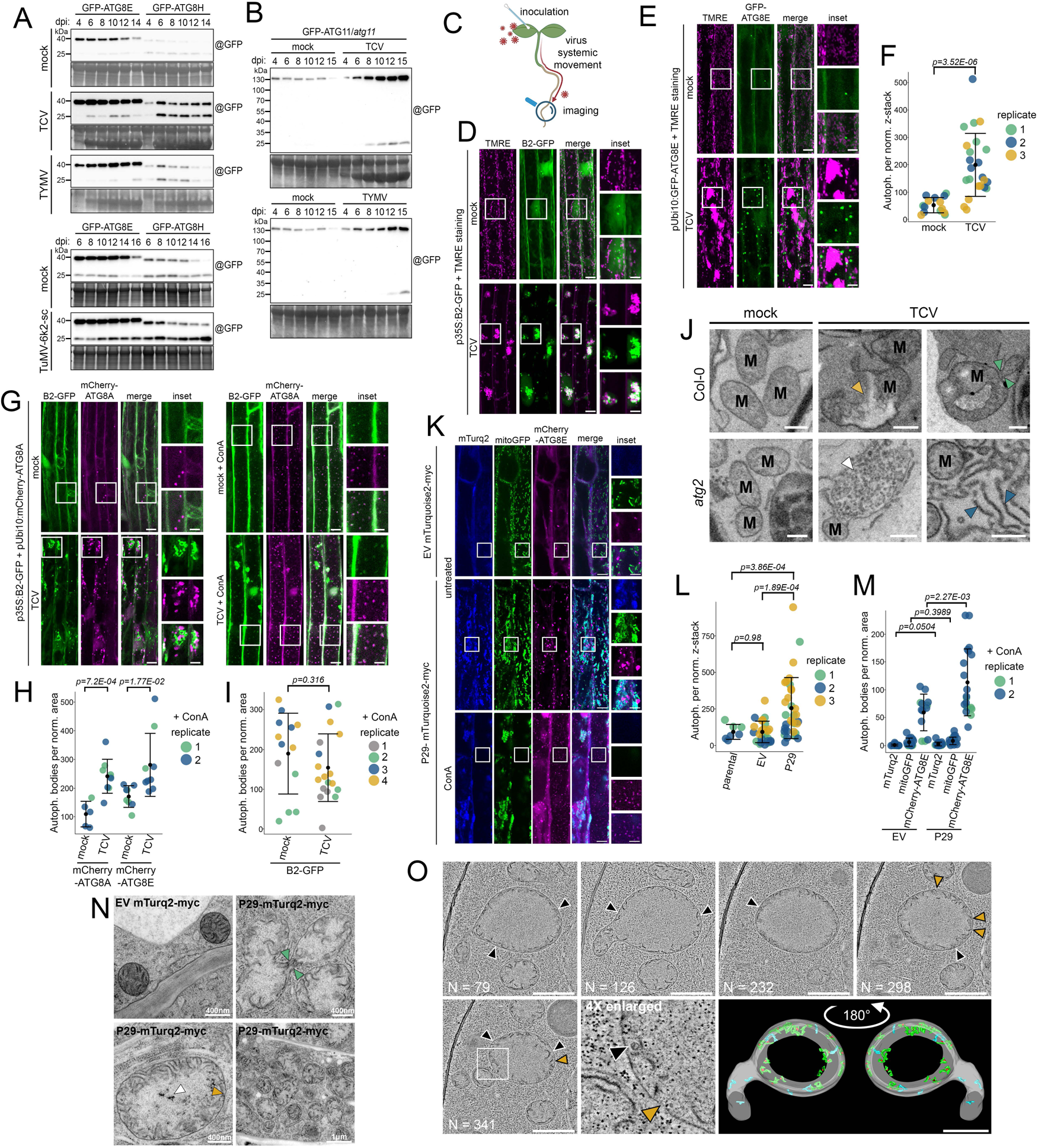
Viral replication organelle remodeling induces autophagy. **(A)** Western blots showing GFP-ATG8E (left) and GFP-ATG8H (right) and GFP cleavage product abundance in mock inoculated, TCV, TYMV and TuMV-6k2-sc-infected systemic leaves. 3-week-old rosettes were inoculated with buffer (mock), with TCV, TYMV or TuMV-6K2-sc and young systemic leaves were collected at the indicated time points (4 to 14 dpi upper panels, 6 to 16 dpi lower panels). n=3 to 4 plants per sample-genotype combination. GFP-ATG8 protein was immunoprecipitated using GFP beads and equal amount of purified protein extract was loaded per well. Membranes were hybridized with a GFP specific antibody. Input samples were loaded separately and stained in gel with Coomassie or on membrane with amidoblack to verify loading. **(B)** GFP cleavage assay of GFP-ATG11 in *atg11 Arabidopsis* stable lines. 3-week-old rosettes were inoculated with buffer (mock), with TCV or with TYMV and young systemic leaves were collected at the indicated time points (4 to 15 dpi). n=3 to 4 plants per sample-genotype combination. 30µg of total protein extract was loaded per well. Membranes were hybridized with a GFP specific antibody and post-stained with amidoblack to verify loading. **(C)** Graphical representation of seedling inoculation with TCV: 9-day-old seedlings are rub-inoculated with TCV sap using two small q-tips on one of the first true leaves and imaging is then performed in systemic roots at the indicated day post inoculation (dpi). An identical setup was also used for further seedling treatment and whole seedling were collected for analysis. **(D)** Confocal microscopy images of *Arabidopsis* root expressing B2-GFP (green) under the 35S promoter and stained with TMRE (magenta) in mock-inoculated or TCV-infected conditions at 5 dpi. Seedlings were immerged in ½ MS with 500nM of TMRE for a few minutes then mounted in ½ MS for image acquisition. Each image is a maximum intensity projection of a full z-stack. Scale bar 10µm and 5µm in inset. See Figure S2D and supplemental movie S5 for additional examples. **(E)** Confocal microscopy images of *Arabidopsis* root expressing GFP-ATG8E (green) under a ubiquitin 10 promoter and stained with TMRE (magenta). 9-day-old seedlings were inoculated with TCV and systemic root images were acquired at 5dpi or 7dpi. Seedlings were immerged in ½ MS with 500nM of TMRE for a few minutes then mounted in ½ MS for image acquisition. Each image is a maximum intensity projection of a full z-stack. Scale bar 10µm and 5µm in inset. **(F)** Quantification of dataset shown in (E). Quantification of GFP-ATG8E puncta in root cells per normalized area and number of slices. Each dot represents the obtained value for a different root and all three biological replicates are shown in a different color. Bars indicate the mean ± SD for each condition. Two-tailed, unpaired with unequal variance Student’s t-test was performed and p.value is displayed above the graph. **(G)** Confocal microscopy images of *Arabidopsis* root expressing B2-GFP (green) under the 35S promoter and mCherry-ATG8A (magenta) under a ubiquitin 10 promoter in mock-inoculated or TCV-infected conditions at 5 and 6 dpi. Left panel: Seedlings were directly mounted in ½ MS for image acquisition. Each image is a maximum intensity projection of a full z-stack. Right panel: Seedlings were immerged in ½ MS with 1µM ConcanamycinA (ConA) for 3.5h to 5h then directly mounted in ½ MS for image acquisition. Each image is a single snap in the focal plane of the vacuole. Scale bar 10µm and 5µm in inset. **(H)** Quantification of mCherry-ATG8A and mCherry-ATG8E (see Figure S2D) puncta inside the vacuole per normalized area in the mock + ConA and the TCV + ConA roots expressing indicated marker and B2-GFP. Each dot represents the obtained value for a different cell/root combination and biological replicates are shown in a different color. Bars indicate the mean ± SD for each condition. Two-tailed, unpaired with unequal variance Student’s t-test was performed and p.value is displayed above the graph. **(I)** Quantification of B2-GFP puncta inside the vacuole per normalized area in the mock + ConA and the TCV + ConA roots. Each dot represents the obtained value for a different cell/root combination and biological replicates are shown in a different color (grey and green in mCherry-ATG8A background and blue and yellow in mCherry-ATG8E background). Bars indicate the mean ± SD for each condition. Two-tailed, unpaired with unequal variance Student’s t-test was performed and p.value is displayed above the graph. **(J)** Representative electron micrographs of mock and TCV infected Col-0 and *atg2* systemic leaf cells at 7 dpi. Infected cells show abnormal mitochondria morphology with notable loss of membrane integrity (yellow arrow) and mitochondria constriction (green arrows). Electron dense material accumulates in *atg2* infected cells, punctate (white arrow) and tubular (blue arrow). M: mitochondria. Scale bar 1µm. **(K)** Confocal microscopy images of *Arabidopsis* root expressing P29-mTurquoise2-myc or empty vector (EV) control (blue) under the ubiquitin 10 promoter, mito-GFP (green) and mCherry-ATG8E (magenta) under a ubiquitin 10 promoter. Seeds were germinated on ½ MS vertical agar plates under long day conditions and 4-day-old seedlings were mounted in ½ MS liquid media and directly imaged (untreated samples). Each image is a maximum intensity projection of a full z-stack. For Concanamycin A treatment, seedlings were immerged in ½ MS with 1µM ConcanamycinA (ConA) and imaged between 1.5 to 4 hours after onset of treatment. Each image is a single snap in the focal plane of the vacuole. Scale bar 10µm and 5µm in inset. **(L)** Quantification of dataset shown in (K). Quantification of GFP-ATG8E puncta in root cells per normalized area and number of slices. Each dot represents the obtained value for a different root and all three biological replicates are shown in a different color. Bars indicate the mean ± SD for each condition. Two-tailed, unpaired with unequal variance Student’s t-test was performed and p.value is displayed above the graph. **(M)** Quantification of dataset shown in (K). Quantification of P29-mTurquoise2-myc, mitoGFP and mCherry-ATG8E puncta inside the vacuole per normalized area in ConA treated roots expressing the indicated markers. Each dot represents the obtained value for a different cell/root combination and biological replicates are shown in a different color. Bars indicate the mean ± SD for each condition. Two-tailed, unpaired with unequal variance Student’s t-test was performed and p.value is displayed above the graph. **(N)** Representative root cells electron micrographs of 4-day-old seedlings expressing P29-mTurquoise2-myc or empty vector (EV) control. Upon expression of the viral protein P29, mitochondria show enlarged and abnormal morphology with marked constrictions (green arrows), loss of membrane integrity (yellow arrow) and electron dense material accumulating in the mitochondrial matrix (white arrows), as well as clumping. Scale is as indicated in the image. **(O)** Transmission electron tomogram series (N = section position) of a mitochondrion in root cells expressing P29. Yellow arrows mark the instance of visible loss of membrane integrity and black arrow the visible small vesicles that associate with cristae or the outer mitochondria membrane. The right-hand panel shows the three-dimensional model of the mitochondrion with a 180° rotation. P29-induced small vesicles are in red and their associated cristae in green. Vesicle-free cristae are rendered in blue and outer mitochondrial membrane in white. Scale bar 1µm. See supplemental video S6 for a movie rendition.

To assess if autophagy degrades VRCs, we monitored the localization of B2-GFP in mCherry-ATG8A (Figure 2G) and mCherry-ATG8E backgrounds (Figure S2D). We observed an increased number of autophagosomes, some of which were in close proximity to B2-positive VRCs (Figure 2G and S2D, left panels). Concanamycin A (ConA) treatment, which is commonly used to block vacuolar degradation ^26^, showed increased autophagic bodies upon viral infection (Figure 2G and S2D right panels, Figure 2H). However, we did not observe a significant increase in vacuolar B2-GFP signal after ConA treatment indicating that VRCs are not significantly targeted for autophagic degradation (Figure 2I).

What drives autophagy induction upon TCV infection? Since we observed an aberrant chondriome alongside increased autophagic activity, we hypothesized that mitochondrial remodeling and clustering could cause autophagy induction. To test this, we interrogated TCV-induced mitochondrial deformation at the ultrastructural level. Serial block face scanning electron microscopy (SBEM) images of TCV-infected Col-0 and *atg2* young systemic leaves revealed altered mitochondria morphologies, with enlarged mitochondria, loss of membrane integrity, membrane proliferation, constrictions, and herniations (Figure 2J and Figure S3). Although no major differences were observed between Col-0 and *atg2* mitochondria morphology, the cytosol of *atg2* leaves appeared filled with electron-dense punctate or tubular structures (Figure 2J and Figure S2E). In one instance, dense punctate material appeared to spill from a ruptured mitochondrion (Figure 2J and Figure S3 bottom series), suggesting some of this unknown material could originate from mitochondria.

To elucidate the connection between mitochondrial hijacking and autophagy activation, we tested if TCV-induced autophagy could be mimicked by the expression of a single mitochondria-clustering viral protein. We chose the P29 protein from TCV relative Melon necrotic spot virus (MNSV isolate Mα5), which typifies a class of *Tombusviridae* proteins that orchestrate VRC assembly by recruiting viral and host factors to mitochondria ^27,28^. Roots expressing P29-mTurquoise2-myc exhibited mitochondrial clumping (Figure 2K), akin to TCV-infected roots (Figure 2D), with P29 colocalizing with the mitochondrial marker. We could measure a robust increase in autophagosomes in P29-expressing plants compared to control mTurquoise2-myc (empty vector, EV) (Figure 2L). This demonstrates that mitochondrial hijacking is sufficient to induce autophagy.

Next, we tested if P29 or mitochondria are autophagy substrates. ConA treatment of P29/ATG8E lines showed that while mCherry-ATG8E autophagic bodies were more abundant in the presence of P29, no P29-mTurquoise2-myc or mito-GFP puncta could be observed in the vacuole (Figure 2K and M). This indicates that neither mitochondria nor the viral protein are autophagy substrates. TEM of P29-expressing roots revealed ultrastructural similarities to TCV infection, including constrictions, membrane integrity loss, proliferation, and mitochondrial matrix aggregates, alongside large clumps of mitochondria (Figure 2N and Figure S2F). Dual-axis tomography and 3D reconstruction further revealed small spherules closely associated with the outer mitochondrial membrane and cristae, which likely constitute apo-replication complexes ready to house viral replication (Figure 2O). Altogether, these findings suggest that viral-induced membrane remodeling triggers autophagy to alleviate cellular stress caused by organelle perturbations.

### Autophagy prevents accumulation of mitochondrial matrix proteins in the cytosol during TCV infection

Given TCV’s tropism and the link between virus-induced mitochondrial damage and autophagy activation, we decided to further explore the connection between TCV-induced autophagy and mitophagy. We first noted that no vacuolar mito-GFP (Figure 2K and M) or loss of membrane potential (Figure S2B and C), both hallmarks of autophagy^20,29,30^, were observed in P29-expressing or TCV-infected plants. Using antibodies against mitochondria-localized proteins, we observed contrasting accumulation patterns: Isocitrate Dehydrogenase (IDH) and Serine Hydroxymethyltransferase (SHMT), two matrix proteins, decreased during TCV infection, while Voltage-Dependent Anion Channel (VDAC) strongly accumulated (Figure 3A, B and Figure S4A). This effect was specific to TCV infection as chloroplast-targeting TYMV did not alter IDH, SHMT or VDAC levels (Figure 3C). IDH and SHMT decay was suppressed in *atg2*, *atg5*, and *atg11* mutants, confirming their autophagic degradation (Figure 3D, Figure S4B and C). We also tested cytochrome c, a component of the electron transport chain, which behaved like VDAC and did not show any evidence for autophagic degradation. The abundance of GDC-H, a matrix protein, was unaffected by infection. However, AOX proteins seemed to be induced by TCV infection with stronger accumulation upon autophagy loss (Figure3D, Figure S4B and C). Since AOX is implicated in mitochondrial retrograde signaling and stress response ^31,32^, whether the observed phenotype stems from lack of autophagic degradation or increased stress in its absence remains to be investigated. While SHMT and IDH degradation was reproducibly blocked in *atg2* plants, VDAC accumulation was insensitive to the genetic background (Figure 3E and Figure S4D). Finally, additional TEM analysis of P29-mTurquoise2myc roots revealed numerous autophagosomes, but only few mitophagosomes (6% of 50 observed) (Figure 3F). Altogether, these results suggest TCV infection induces selective mitochondrial protein degradation, rather than wholesale mitophagy ^20^.

**FIGURE 3:**
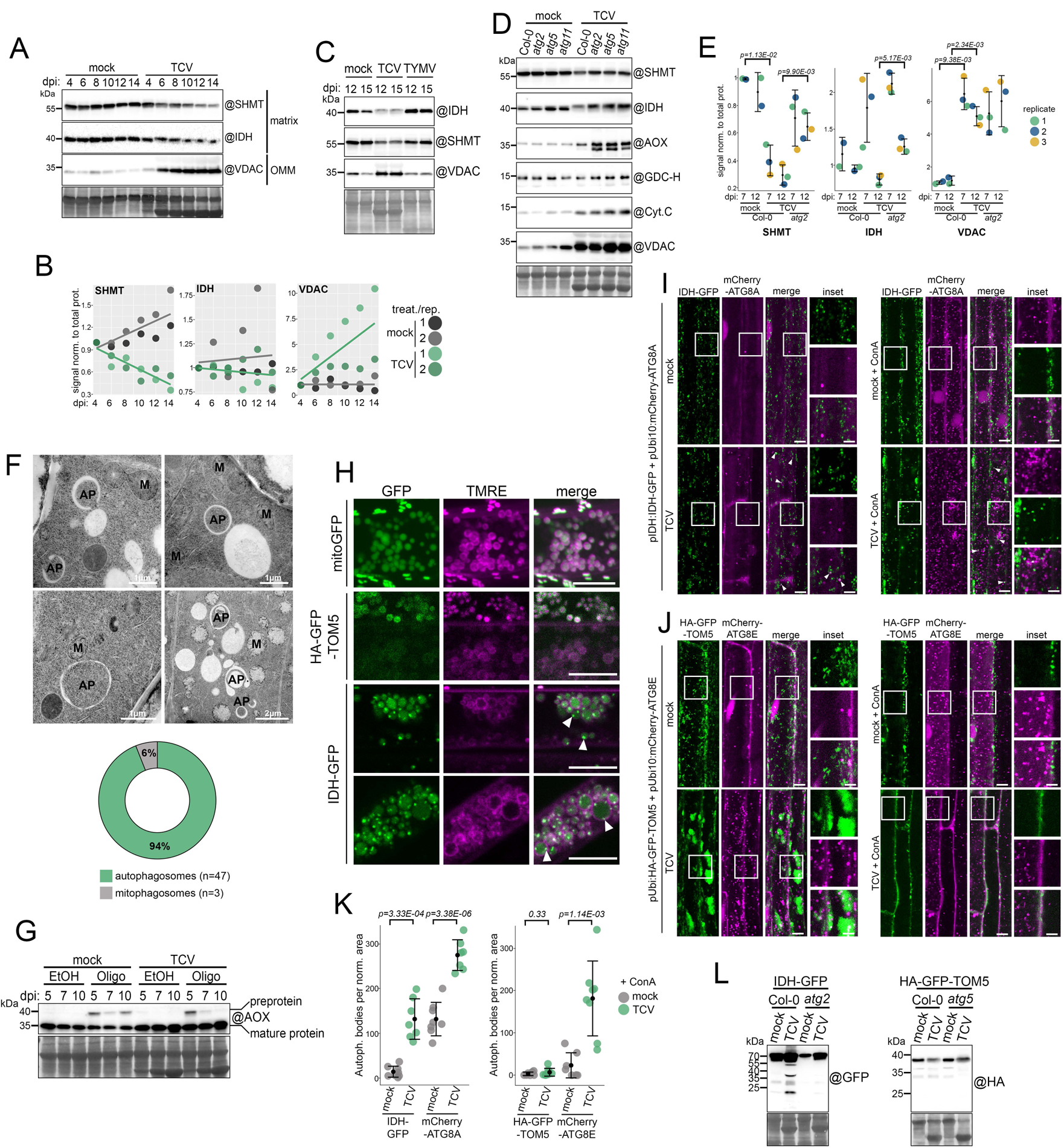
Autophagy degrades single mitochondrial proteins that leak into the cytosol. **(A)** Immunoblot of mitochondrial proteins in mock-inoculated and TCV infected systemic leaves. 3-week-old rosettes were inoculated with buffer (mock) or with TCV and young systemic leaves were collected at the indicated time points. n=3 to 4 plants per sample-timepoint combination. Total soluble proteins were extracted at a fixed fresh weight to buffer volume ratio and equal volume was loaded per well. Membranes were hybridized with the indicated antibodies (@). Membranes were then stained with amidoblack to verify loading. A biological replicate experiment is available in Figure S4A. **(B)** Quantification of protein abundance from Figure 3A (replicate 1) and Figure S4A (replicate 2). Signal intensity is normalized to total protein loading and expressed relative to the 4 dpi time point and plotted for each time point in the kinetic. Solid lines are linear regression across the kinetic for the indicated treatment. **(C)** Immunoblot of mitochondrial proteins in mock inoculated, TCV and TYMV infected systemic leaves. 3-week-old rosettes were inoculated and young systemic leaves were collected at 12 and 15 dpi. n=4 plants per treatment-timepoint combination. Total soluble proteins were extracted at a fixed fresh weight to buffer volume ratio and equal volume was loaded per well. Membranes were hybridized with the indicated antibodies (@). Membranes were then stained with amidoblack to verify loading. **(D)** Immunoblot of mitochondrial proteins in mock and TCV inoculated systemic leaves of WT, *atg2*, *atg5* and *atg11* plants. 3-week-old rosettes were inoculated with buffer (mock) or with TCV and young systemic leaves were collected at 12 dpi. n=10 plants. Total soluble proteins were extracted at a fixed fresh weight to buffer volume ratio and equal volume was loaded per well. Membranes were hybridized with the indicated antibodies (@). Membranes were then stained with amidoblack to verify loading. See Figure S4C for biological replicates. **(E)** Quantification of SHMT, IDH and VDAC protein abundance in mock and TCV inoculated systemic leaves of WT and *atg2* plants. 3-week-old rosettes were inoculated with buffer (mock) or with TCV and young systemic leaves were collected at the indicated time points. Signal intensity is normalized to total protein loading and expressed relative to Col-0 mock samples. See Figure S4D for immunoblot data used for quantification. **(F)** Top panel: Representative root cells electron micrographs of 4-day-old seedlings expressing P29-mTurquoise2-myc. Change in mitochondria morphology is accompanied by the formation of autophagosomes engulfing cytosolic material, rather than mitophagosomes. AP: autophagosome, M: mitochondria. Scale is as indicated in the image. Bottom panel: Quantification of relative abundance of auto/mitophagosomes expressed in percentage (94% autophagosomes, 6% mitophagosomes). A total of 30 images were used for manual counting, out of 6 individual roots. **(G)** Western blots showing mature and pre-protein of endogenous AOX1/2 protein from *Arabidopsis* whole seedling. 9-day-old seedlings were mock-inoculated or TCV-infected and grown for an additional 5, 7 and 10 days before being transferred to liquid ½ MS + 1% sucrose with either ethanol (EtOH, vehicle) or 20µM Oligomycin A (Oligo) for 6 hours. 15 µg of total protein was loaded per well and membrane was hybridized with an antibody against AOX1/2 protein. Membrane was stained with amidoblack to verify loading. Two biological replicate experiments are available in Figure S4E and G. **(H)** Spinning disk microscopy image (mito-GFP) and confocal microscopy image (IDH-GFP and HA-GFP-TOM5) of *Arabidopsis* roots expressing the indicated reporter protein stained with TMRE (magenta) and displaying swollen mitochondria. 9-day-old seedlings were inoculated with TCV and systemic root images were acquired at 7dpi. Seedlings were immerged in ½ MS with 500nM of TMRE for a few minutes then mounted in ½ MS for image acquisition. Each image is a maximum intensity projection of a z-stack. Scale bar 10µm. **(I)** Confocal microscopy images of *Arabidopsis* root expressing IDH-GFP (green) under its own promoter and mCherry-ATG8A (magenta) under a ubiquitin 10 promoter in mock-inoculated or TCV-infected conditions at 7 dpi. Left panel: Seedlings were directly mounted in ½ MS for image acquisition. Each image is a maximum intensity projection of a full z-stack. Right panel: Seedlings were immerged in ½ MS with 1µM ConcanamycinA (ConA) for 3.5h to 5h then directly mounted in ½ MS for image acquisition. Each image is a single snap in the focal plane of the vacuole. Scale bar 10µm and 5µm in inset. **(J)** Confocal microscopy images of *Arabidopsis* root expressing HA-GFP-TOM5 (green) under a ubiquitin promoter and mCherry-ATG8E (magenta) under a ubiquitin 10 promoter in mock-inoculated or TCV-infected conditions at 7 dpi. Left panel: Seedlings were directly mounted in ½ MS for image acquisition. Each image is a maximum intensity projection of a full z-stack. Right panel: Seedlings were immerged in ½ MS with 1µM ConcanamycinA (ConA) for 3h to 4.5h then directly mounted in ½ MS for image acquisition. Each image is a single snap in the focal plane of the vacuole. Scale bar 10µm and 5µm in inset. **(K)** Quantification of IDH-GFP, mCherry-ATG8A (left panel), HA-GFP-TOM5 and mCherry-ATG8E (right panel) puncta inside the vacuole per normalized area in the mock -/+ ConA and the TCV -/+ ConA roots expressing the indicated markers. Each dot represents the obtained value for a different cell/root combination. Bars indicate the mean ± SD for each condition. Two-tailed, unpaired with unequal variance Student’s t-test was performed and p.value is displayed above the graph. **(L)** Western blots showing HA-GFP-TOM5 and IDH-GFP cleavage level in mock and TCV inoculated systemic leaves of Col-0, *atg5* or *atg2*. 3-week-old rosettes were inoculated with buffer (mock) or with TCV and young systemic leaves were collected at the indicated time points. n=10 plants per condition. Total proteins were extracted and 25µg of total protein was loaded per well and membranes were hybridized with the indicated antibodies (@). Membranes were then stained with amidoblack to verify loading. A biological replicate experiment is available in Figure S4H.

As autophagy occurs exclusively in the cytosol, we hypothesized that IDH and SHMT should leak to the cytosol prior to degradation. We considered two scenarios leading to this situation: (i) accumulation of mature mito-proteins in the cytosol due to organelle rupture or (ii) inhibition of the mitochondrial import pathway resulting in cytosolic accumulation of mitochondrial pre-proteins. To test these scenarios, we used oligomycin, a mitochondrial H^+^-ATP-synthase inhibitor causing hyperpolarization of the inter-membrane space and depletion of mitochondrial ATP required for import ^33,34^. Oligomycin treatment led to the accumulation of AOX precursor at the expected size ^35^ (cleavage at position 62-63 predicted by TargetP2.0), while no pre-protein was observed in TCV-infected seedlings (Figure 3G and Figure S4E). Accordingly, oligomycin treatment induced partial dissipation of membrane potential indicated by lack of TMRE staining, which was not observed in TCV-infected roots (Figure S4F). Combining TCV infection with oligomycin treatment led to severe mitochondria swelling, depolarization, and accumulation of pre-protein, whose abundance was not autophagy-dependent (Figure S4G). This suggests that TCV infection does not lead to a widespread inhibition of mitochondrial protein import, and that autophagic degradation of IDH and SHMT is caused by their leakage to the cytosol. This is consistent with our electron microscopy data where we have observed loss of membrane integrity during infection (Figure 2J, N, O).

We then compared the localization of two categories of mitochondrial proteins (*i.e* autophagy cargo or not). We chose IDH-GFP ^36^ as a representative cargo, while mito-GFP ^37^ and HA-GFP-TOM5 (Translocase at the Outer Membrane5) ^38^ served as model non-cargo mitochondrial proteins. During infection, we observed a minor population of mitochondria that had undergone swelling and contained bright IDH-GFP puncta (Figure 3H), which were not observed for mito-GFP, suggesting aggregation of the model cargo. Colocalization experiments revealed that in infected roots, autophagosomes (labeled by mCherry-ATG8A) engulfed bright IDH-GFP puncta (Figure 3I, left panel) that could be seen moving together (Supplemental video 7). Accordingly, ConA treatment specifically led to the accumulation of IDH-GFP puncta in the vacuole of TCV-infected root cells (Figure 3I and K). We detected clumped GFP signal in close proximity to autophagosomes when we imaged HA-GFP-TOM5 under identical conditions (Figure 3J), consistent with our previous observations (Figure 2G), but no autophagosome engulfing discrete GFP signal. Additionally, ConA treatment did not lead to the accumulation of HA-GFP-TOM5 in the vacuole (Figure 3J and K). We then employed GFP release assay to monitor autophagy flux in systemic shoot tissue for both proteins. TCV infection led to accumulation of multiple GFP cleavage products from IDH-GFP, which were absent in both *atg2* and *atg5* plants (Figure 3L and Figure S4H), indicating autophagy-dependent vacuolar degradation of IDH-GFP. In contrast, while a ∼30 kDa cleavage product could be observed for HA-GFP-TOM5, it was neither sensitive to infection nor to the loss of autophagy (Figure 3L and Figure S4H), indicating that TOM5 is not a substrate for TCV-induced autophagy. Overall, these results indicate that TCV infection does not lead to wholesale mitophagy, but rather to the deployment of autophagy for targeted degradation of leaked mitochondrial proteins in the cytosol following virus-induced mitochondrial rupture.

### Autophagy engages two metabolic enzymes as selective autophagy receptors and counteracts hyperactive immunity during infection

To uncover selective autophagy receptors involved in viral infection, we conducted an ATG8 affinity purification-mass spectrometry (AP-MS) screen coupled with peptide competition, a method we previously utilized to identify AIM-dependent ATG8 interactors during ER stress ^39^. We employed a three-tiered approach to categorize the interactome: (*i*) ATG8-specific interactors compared to control AP, (*ii*) AIM-dependent ATG8 interactors depleted upon treatment with AIM *wt* but not with a mutated (*mut*) AIM peptide, and (*iii*) virus-induced interactors, proteins enriched in the presence of viruses compared to mock. Furthermore, we reasoned that if autophagy constitutes a general response to infection, we should retrieve the same candidate SARs from both TCV and TYMV-infected samples. Given the absence of specificity for ATG8 isoform induction in the presence of viruses (Figure 2A, G, and H), we opted to conduct the same AP with both mCherry-ATG8A and mCherry-ATG8E. Following pairwise comparison for each AP, overlapping proteins were considered candidate SARs only if they appeared in at least three out of the four sets of AP-MS experiments (Figure S5A). For mCherry-ATG8E, infection led to enrichment of several host proteins (Figure 4A and D) with 26 and 39 proteins meeting all three criteria for TCV and TYMV infection, respectively (Figure 4B and E, Tables S5 and S6). The most prominent interactors of mCherry-ATG8E were ATG3 and ATG7 and not outcompeted by AIM *wt* treatment (Figure S5B and C). Similar results were obtained for mCherry-ATG8A (Figure S5D-I, Tables S6 and S7). Overall, 16 candidate SARs met our criteria (Figure 4C, F and Figure S5F, I). We decided to focus on Nitrilase1 (NIT1, AT3G44310) and Inosine Monophosphate Dehydrogenase1 (IMPDH1, AT1G79470), two metabolic enzymes not previously linked to autophagy, of which several paralogues were detected in the ATG8-enriched interactome.

**FIGURE 4:**
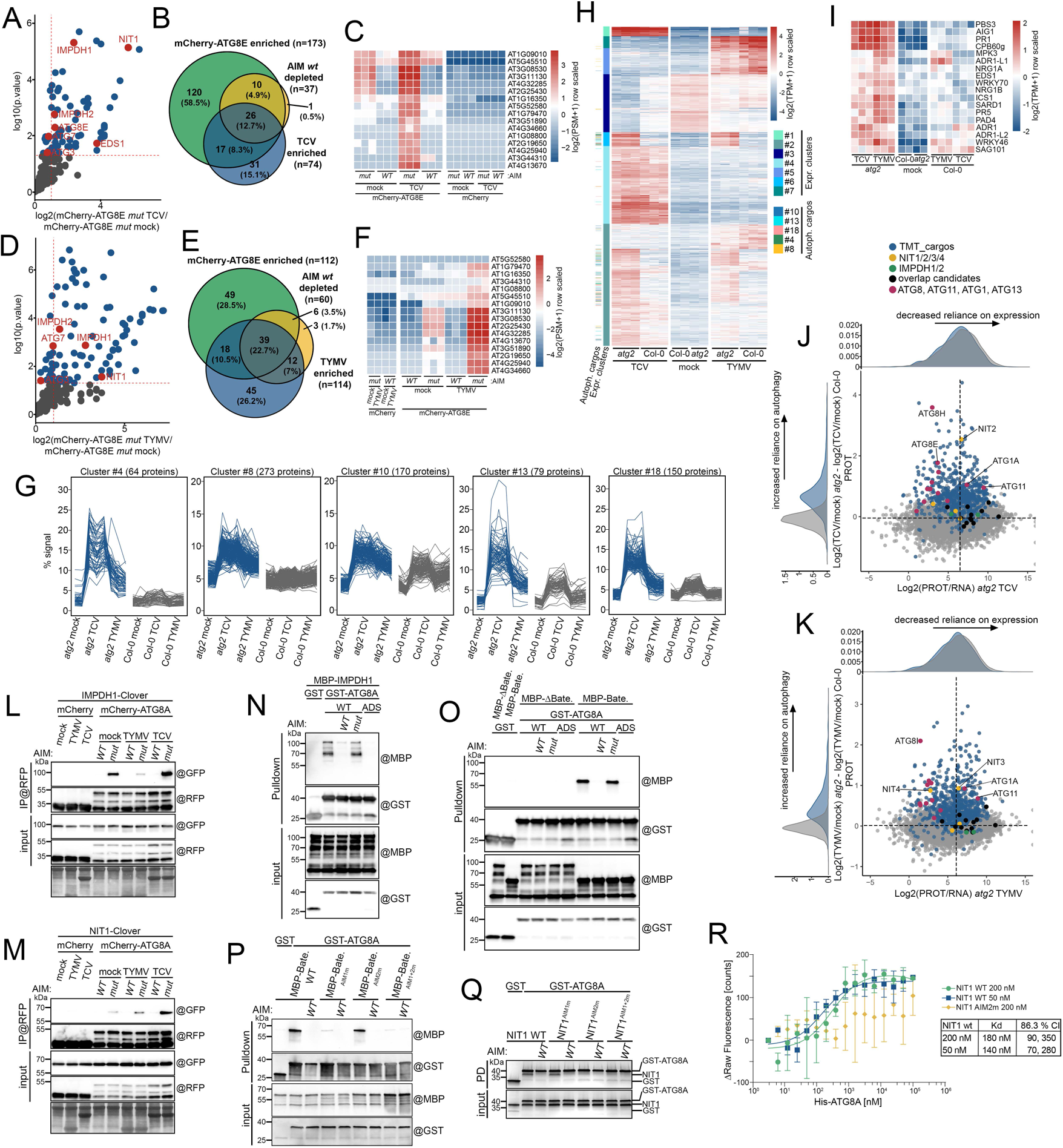
NIT1 and IMPDH1, two metabolic enzymes, are virus-specific selective autophagy receptors. **(A)** Enrichment of proteins co-purified with mCherry-ATG8E in TCV-infected systemic leaves *vs.* mock leaves of the same age, represented by a volcano plot. The horizontal dashed line indicates the threshold above which proteins are significantly enriched (*p* value < 0.05, quasi-likelihood negative binomial generalized log-linear model) and the vertical dashed line the threshold for which proteins log2 fold change is above 1. **(B)** Venn diagram of three overlapping pairwise comparisons for AP-MS conducted in mCherry-ATG8E plants infected with TCV: mCherry-ATG8E (only mutant AIM-treated replicates) *vs.* mCherry control; green circle. mCherry-ATG8E AIM mutant-treated samples (mock and TCV treated combined) *vs.* mCherry-ATG8E AIM *wt*-treated samples; yellow circle. mCherry-ATG8E TCV-treated samples (mutant AIM-treated only) *vs.* mCherry-ATG8E mock-treated samples (mutant AIM-treated only); blue circle. **(C)** Protein abundance pattern represented by a heatmap (Log2(PSM+1) – meanPSM per protein) for the sixteen proteins identified as uniquely enriched in AIM-dependent ATG8 interactome upon infection (at least three of four datasets), in the mCherry-ATG8E TCV dataset. **(D)** Enrichment of proteins co-purifed with mCherry-ATG8E in TYMV-infected systemic leaves *vs.* mock leaves of the same age, represented by a volcano plot. The horizontal dashed line indicates the threshold above which proteins are significantly enriched (*p* value < 0.05, quasi-likelihood negative binomial generalized log-linear model) and the vertical dashed line the threshold for which proteins log2 fold change is above 1. **(E)** Venn diagram of three overlapping pairwise comparisons for AP-MS conducted in mCherry-ATG8E plants infected with TYMV: mCherry-ATG8E (only mutant AIM-treated replicates) *vs.* mCherry control; green circle. mCherry-ATG8E AIM mutant-treated samples (mock and TYMV treated combined) *vs.* mCherry-ATG8E AIM *wt*-treated samples; yellow circle. mCherry-ATG8E TYMV-treated samples (mutant AIM-treated only) *vs.* mCherry-ATG8E mock-treated samples (mutant AIM-treated only); blue circle. **(F)** Protein abundance pattern represented by a heatmap (Log2(PSM+1) – meanPSM per protein) for the sixteen proteins identified as uniquely enriched in AIM-dependent ATG8 interactome upon infection (at least three of four datasets), in the mCherry-ATG8E TYMV dataset. **(G)** k-means clustering of protein abundance reveals autophagy-dependent degradome in infected leaves. Protein abundance was measured in mock, TCV and TYMV infected systemic leaves of Col-0 and *atg2* plants (duplicates for mock and triplicates for infected samples) using 16-plex TMT labelling. Normalized abundance of proteins regulated > 2-fold was then used for k-means clustering and sorted in 20 separate clusters. Five clusters that showed increased abundance in *atg2* only during infection were used to obtain the list of candidate autophagy cargos proteins, specific to viral infection. All 20 clusters are represented in Figure S6. **(H)** Transcriptomic analysis of infected *atg2* systemic leaves reveals virus-specific and *atg2*-specific transcriptional reprograming. Abundance of transcripts (Log2(TPM+1) – meanTPM per transcript) that are significantly deregulated by infection with either TCV and TYMV in Col-0 and in *atg2* are displayed in a heatmap. Amongst 7 expression clusters, cluster #2 and cluster #6 contain transcripts that are upregulated in infected *atg2* plants, regardless of the virus. Mapping of virus-specific cargos defined in Figure 4G shows that they are enriched in the *atg2*-affected expression clusters. **(I)** Expression of several defense-related genes is elevated in infected *atg2* systemic tissue. Gene expression is represented as in Figure 4H. **(J)** Graphical representation of the ratio of protein abundance to RNA abundance in *atg2* TCV samples (log2(protein abundance/transcript abundance) as captured by quantitative proteomics and transcriptomic) on the x axis versus the ratio of protein abundance in atg2 TCV-infected samples with the removed contribution of the WT Col-0 background (log2(TCV/mock) in *atg2* – log2(TCV/mock) in Col-0) as captured by quantitative proteomics on the y axis. Reliance on transcription for overall abundance of a protein decreases as x axis values increase, while increased reliance on the autophagic machinery at the protein level increases with the y axis. Quadrants are determined using centroids for all the plotted datapoints. Data distribution for both axis is plotted next to the relevant axis. Proteins mapped as virus-specific autophagy cargos are colored in blue, known autophagy machinery components that undergo degradation are plotted in magenta and the detectable candidate SARs identified by AP-MS are plotted in black. NIT paralogues are plotted in yellow and IMPDH-paralogues in green (the later are not present in the cargo list). **(K)** Graphical representation of the ratio of protein abundance to RNA abundance in *atg2* TYMV samples (log2(protein abundance/transcript abundance) as captured by quantitative proteomics and transcriptomic) on the x axis versus the ratio of protein abundance in atg2 TYMV-infected samples with the removed contribution of the WT Col-0 background (log2(TYMV/mock) in *atg2* – log2(TYMV/mock) in Col-0) as captured by quantitative proteomics on the y axis. Reliance on transcription for overall abundance of a protein decreases as x axis values increase, while increased reliance on the autophagic machinery at the protein level increases with the y axis. Quadrants are determined using centroids for all the plotted datapoints. Data distribution for both axis is plotted next to the relevant axis. Proteins mapped as virus-specific autophagy cargos are colored in blue, known autophagy machinery components that undergo degradation are plotted in magenta and the detectable candidate SARs identified by AP-MS are plotted in black. NIT paralogues are plotted in yellow and IMPDH-paralogues in green (the latter are absent from the cargo list). **(L)** *In vivo* affinity purification of mCherry-ATG8A coupled to AIM peptide competition confirms AIM-dependent interaction with IMPDH1-Clover. Systemic leaves expressing mCherry control or mCherry-ATG8A under ubiquitin 10 promoter and IMPDH1-Clover under HTR5 promoter in mock-inoculated, TYMV or TCV-infected conditions at 11 dpi (n=40 plants). 150 µM of AIM *wt* or mutant was added to the total cell lysate and affinity purification was performed using @RFP beads. Immunoblot against the indicated antibodies (@) was used to detect bait and prey in the input and RFP purified fraction. Membrane corresponding to the input fraction was then stained with amidoblack to verify loading. See Figure S8D for biological replicate. **(M)** *In vivo* affinity purification of mCherry-ATG8A coupled to AIM peptide competition confirms AIM-dependent interaction with NIT1-Clover. Systemic leaves expressing mCherry control or mCherry-ATG8A under ubiquitin 10 promoter and NIT1-Clover under HTR5 promoter in mock-inoculated, TYMV or TCV-infected conditions at 11 dpi (n=40 plants). 150 µM of AIM *wt* or mutant peptide was added to the total cell lysate and affinity purification was performed using @RFP beads. Immunoblot against the indicated antibodies (@) was used to detect bait and prey in the input and RFP purified fraction. Membrane corresponding to the input fraction was then stained with amidoblack to verify loading. See Figure S8E for biological replicate. **(N)** IMPDH1 interacts with ATG8A in an AIM-dependent manner. Bacterial lysates containing recombinant MBP-IMPDH1, GST, GST-ATG8A WT or GST-ATG8A deficient in the AIM docking site (ADS) were mixed, 200 µM AIM *wt* or mutant peptide was added to the indicated reaction and proteins were pulled down using GST beads. Immunoblot against the indicated antibodies (@) was used to detect bait and prey in the input and pull-down fraction. See Figure S8F for a biological replicate. **(O)** The bateman regulatory domain of IMPDH1 is necessary and sufficient to mediate interaction with ATG8A. Bacterial lysates containing recombinant MBP-Bateman, MBP-ΔBateman, GST, GST-ATG8A WT or GST-ATG8A ADS were mixed, 150 µM AIM *wt* or mutant peptide was added to the indicated reaction and proteins were pulled down using GST beads. Immunoblot against the indicated antibodies (@) was used to detect bait and prey in the input and pull-down fraction. See Figure S8G for a biological replicate. **(P)** The AIM1 contained in the Bateman regulatory domain of IMPDH1 mediates interaction with ATG8A. Bacterial lysates containing MBP-Bateman WT, derivative AIM mutants, GST and GST-ATG8A were mixed with, 200 µM AIM peptide was added to the indicated reaction and proteins were pulled down using GST beads. Immunoblot against the indicated antibodies (@) was used to detect bait and prey in the input and pull-down fraction. **(Q)** The C-terminal AIM (AIM2) of NIT1 is necessary and sufficient to mediate interaction with ATG8A. Purified recombinant NIT1 WT, NIT1^AIM1m^, NIT1^AIM2m^, NIT1^AIM1+2m^ were mixed with purified recombinant GST or GST-ATG8A, 200 µM AIM WT or mutant peptide was added to the indicated reaction and proteins were pulled down using GST beads. In-gel Coomassie staining was used to detect bait and prey proteins in the input and pull-down fraction. See Figure S8I for additional experimental evidence. **(R)** ΔRaw Fluorescence curves of ATG8A with NIT1 WT (binding) and NIT1^AIM2m^ (no binding) measured with Monolith NT.115. NIT1 WT shows a Kd in the nM range, with an 86.3 % confidence interval (CI) between 70 nM and 350 nM. The error bars describe the standard deviation of each data point calculated from three independent experiments. Binding controls show no interaction between ATG8A and dye. The ligand-induced fluorescence change specificity was tested by the SDS denaturation test (SD-Test) (see S8J).

We then conducted tandem mass tag (TMT) labeling-based quantitative proteomics analysis on systemic leaves of Col-0 and *atg2* plants that were either mock-inoculated or infected with TCV/TYMV. A total of 9586 proteins were detected across all conditions. Employing k-means clustering on proteins with a fold change > 2 resulted in 20 distinct clusters (2833 proteins, Figure S6). This clustering approach revealed proteins that were uniquely up-and down-regulated by each virus, including TCV ORFs (cluster #6) and TYMV ORFs (cluster #11) (Figure S6, Table S8), and identified 735 proteins that were specifically up-regulated in infected *atg2* plants (Figure 4G). Notably, we did not observe clusters of *atg2*-sensitive proteins that were virus-specific, indicating that plant autophagy did not differentiate between TCV and TYMV infections, suggesting a shared degradome despite their differing tropisms. We classified these proteins as autophagy cargos.

Given the pronounced and persistent symptoms observed in infection, we decided to evaluate transcriptional changes during infection. As expected, differential gene expression analysis revealed that many transcripts underwent virus-specific deregulation (Figure 4H, Tables S9 to S12). Surprisingly, cluster #2 and #6 contained transcripts that increased only in *atg2* infected samples and were enriched for genes encoding autophagy cargos identified earlier (Figure 4H). We observed elevated expression of several well-known defense genes (Figure 4I), which was mostly absent in wild-type, resembling an autoimmune response ^40–42^. To better characterize the response of infected *atg2* plants, we analysed expression of defense genes identified in other studies. We could observe an increase of transcripts identified as local responders to *Pseudomonas syringae* (*Pto*) carrying the effector *AvrRpm1* in our dataset, which overlapped with identified cargo proteins (Figure S7A), while we could not observe any association with genes defined as distal-responsive genes in the same study ^43^. A similar pattern was observed for Brassicaceae-conserved flg22-responsive genes (Figure S7B) ^44^, and for the downstream regulon of the EDS1/PAD4 heterodimer (Figure S7C) ^45^, a key sensing and signaling component of plant defense. Out of the 735 identified autophagy cargos, 266 (36.2%) were also differentially expressed genes (DEGs), with the vast majority (95,4%) being affected in *atg2* (Figure S7D). The cargo-DEG category was enriched in gene ontology (GO) molecular functions associated with defense, immunity and response to stress, while the remaining cargos showed enrichment for oxidative stress and metabolism-associated GO terms (Figure S7E). To analyse cargo behavior in the context of expression, we represented protein abundance as a ratio to RNA abundance in *atg2* infected samples (x axis), against the infected-to-mock ratio in *atg2*, subtracted from the influence of Col-0 (y axis). This approach highlights genes in the top-right quadrant that exhibit increased reliance on autophagy and decreased reliance on transcriptional regulation to explain their apparent abundance (Figure 4J and K). Cargo proteins (in blue) showed a sharp separation on the y-axis, while their distribution on the x-axis was largely overlapping with the rest of the detected genes and was equally distributed around the centroid. Many key autophagy genes (maroon) were found in the top-left quadrant suggesting a combined transcriptional and post-translational regulation while candidates identified by AP-MS (black) showed only mild autophagy-dependent stabilization, with the exception of NIT genes (yellow) that were strongly affected in *atg2*. IMPDH1/2 (green), which failed to cluster in our previous strategy, were found close to the centroid. A similar picture emerged when Col-0 proteome and expression data was used for the x-axis (Figure S8A and B). Finally, we showed that Glycolate Oxidase1 (GLO1) a non-DEG cargo, was consistently degraded by autophagy in infected samples (Figure S8C). These results suggest autophagy tailors the defense-related proteome to prevent cellular collapse and self-amplification of an immune transcriptional feedback loop ^41^.

We then set out to characterize the interaction of IMPDH1 and NIT1 with ATG8. Co-immunoprecipitation experiments showed an AIM-dependent interaction between IMPDH1-Clover and mCherry-ATG8A (Figure 4L) and mCherry-ATG8E (Figure S8D). We also detected an AIM-dependent interaction between NIT1-Clover and both ATG8 reporters, which became stronger upon viral infection (Figure 4M and Figure S8E). Consistently, using an antibody against NIT1/2/3 proteins (Figure S11A), we could recover Nitrilase in infected samples for both ATG8 isoforms tagged with mCherry (Figure S8E). As autophagy receptors interact with ATG8 directly via an AIM ^46,47^, we investigated the IMPDH1-ATG8A interaction in *E. coli* lysates using GST-pulldowns. MBP-IMPDH1 interacted with GST-ATG8A but not with an AIM-docking site mutant (ADS) ^47,48^, and the interaction was out-competed by addition of the AIM *wt* peptide (Figure 4N and Figure S8F). IMPDH consists of a catalytic domain and a Bateman regulatory domain inserted within a loop of the catalytic domain ^49,50^. Bioinformatics analysis predicted 6 AIMs for AtIMPDH1, equally divided between both domains (Figure S9). GST-pulldowns with MBP-IMPDH1 ΔBateman and MBP-IMPDH1 Bateman domain identified this domain as interacting with GST-ATG8A (Figure 4O and Figure S8G). Upon examination of the sequence conservation of IMPDH proteins, we selected two candidate AIMs: AIM1, which is positionally conserved in flowering plants, algae and animals but absent in Solanaceae, and AIM2, which was conserved in land plants (Figure S9A). Phe128Ala mutation at position Θ of AIM1 in MBP-Bateman led to a strongly reduced interaction with GST-ATG8A (Figure 4P), a finding mirrored when full-length MBP-IMPDH1 was mutated (Figure S8H). These results indicate AIM1 mediates ATG8 binding. Next, we purified AtNIT1 recombinantly expressed in *E. coli* and conducted pulldowns with GST-ATG8A. Co-elution of NIT1 with GST-ATG8A was evident from 1µM of purified NIT1, and the interaction was disrupted by addition of AIM *wt* peptide (Figure S8I). Examination of the NIT protein sequence conservation highlighted two likely AIMs positioned at the N-and C-terminal disordered region of the protein (Figure S10) that are unresolved in the structure of AtNIT4 ^51^. Phe16Ala/Val19Ala (AIM1) and Phe307Ala/Val310Ala (AIM2) mutations in positions Θ and Γ showed that the C-terminal AIM2, conserved in all plant species tested (Figure S10), was necessary and sufficient to mediate the interaction to GST-ATG8A (Figure 4Q). MicroScale Thermophoresis (MST) analysis of NIT1-ATG8A interaction also confirmed that NIT1 binds ATG8A with low nanomolar affinity (K_D_ = ∼150nM), and AIM2 mutant lost its ability to bind ATG8A (Figure 4R and S8J). In conclusion, we reveal two metabolic enzymes as novel selective autophagy receptors that are induced during infection with both TCV and TYMV.

### Nitrilase filament-to-monomer switch controls EDS1 abundance and cell death

To characterize the *in vivo* dynamics of NIT1 and IMPDH1, we employed autophagic flux assays. Employing a NIT1-clade ^52^ specific antibody (Figure S11A), in carbon and nitrogen starvation assays, we showed that Nitrilase abundance is unchanged in seedlings, even with addition of ConA or genetic ablation of autophagy (Figure 5A, B, C and Figure S11B), similar to the mitochondrial matrix protein IDH. However, NBR1 was sensitive to all perturbations. We next introduced our reporter constructs into the *atg5* background and performed a Clover release assay during infection. We observed a consistent increase of free Clover for both TCV and TYMV (Figure 5D and Figure S11C), which was absent in *atg5* (Figure 5D), indicative of its autophagic origin. In line with the recovery of NIT proteins as autophagy cargos by our TMT approach (Figure 4J and K), we noted a significant increase in total NIT1-clade proteins in infected *atg* mutants (Figure 5E, F, G, H and Figure S11D, E). Transient co-expression of NIT1-mCherry and IMPDH1-mCherry with a TCV infectious clone in stable B2-GFP *N. benthamiana* ^25^ revealed accumulation of vacuolar mCherry puncta in infected cells (Figure S11F). Consistently, ConA treatment led to the vacuolar accumulation of both reporter proteins in Arabidopsis, suggesting both enzymes undergo autophagic degradation during viral infection (Figure 5I and Figure S11G).

**FIGURE 5:**
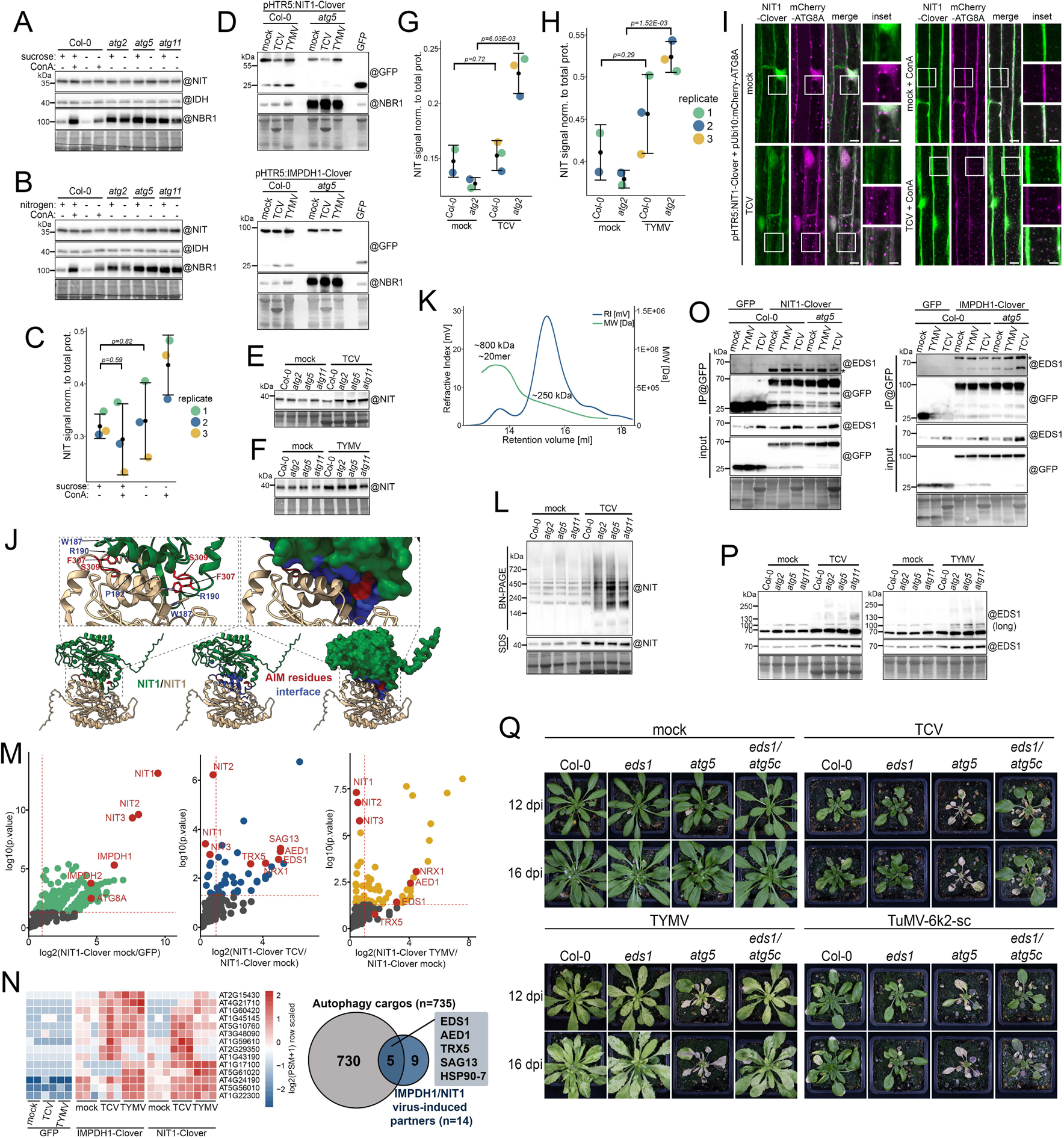
Conditional activation of SAR activity by oligomerization controls EDS1 abundance and suppresses cell death. **(A)** Autophagic flux in 7-day-old Col-0, *atg2*, *atg5* and *atg11* seedlings treated with or without carbon starvation (top panel), with or without 1µM ConcanamycinA (ConA) for 24 hours. 10 µg of total protein was loaded per well and membranes were hybridized with the indicated antibodies (@). Membranes were then stained with amidoblack to verify loading. **(B)** Autophagic flux in 7-day-old Col-0, *atg2*, *atg5* and *atg11* seedlings treated with or without nitrogen starvation (bottom panel), with or without 1µM ConcanamycinA (ConA) for 24 hours. 10 µg of total protein was loaded per well and membranes were hybridized with the indicated antibodies (@). Membranes were then stained with amidoblack to verify loading. **(C)** Quantification of endogenous NIT protein signal in 8-day-old seedlings treated with or without carbon starvation with or without 1µM ConcanamycinA (ConA) for 24 hours. Signal intensity is normalized to total protein signal. Signal intensity was measured from biological triplicates presented in Figure S11B. **(D)** Autophagic flux in systemic leaves of plants expressing NITRILASE1-Clover (NIT1-clo, top panel) and IMPDH1-Clover (IMPDH1-clo, bottom panel) under the HTR5 promoter in a Col-0 or *atg5* background. Plants were either treated with buffer (mock), TCV or TYMV and samples were collected at 11 dpi. n=12-21 plants per genotype∼treatment combination. 15µg of total protein was loaded per well and membranes were hybridized with a GFP antibody (@). Membranes were then stained with amidoblack to verify loading. See Figure S11C for a biological replicate. **(E)** Immunoblot of endogenous NIT protein content in mock and TCV inoculated systemic leaves of Col-0, *atg2*, *atg5* and *atg11* plants. 3-week-old rosettes were inoculated with buffer (mock) or with TCV and young systemic leaves were collected at 12 dpi. n=10 plants. Total soluble proteins were extracted at a fixed fresh weight to buffer volume ratio and equal volume was loaded per well. Membranes were hybridized with the NIT antibody (@). Membranes were then stained with amidoblack to verify loading. See Figure S11D for a biological replicate. **(F)** Immunoblot of endogenous NIT protein content in mock and TYMV inoculated systemic leaves of Col-0, *atg2*, *atg5* and *atg11* plants. 3-week-old rosettes were inoculated with buffer (mock) or with TYMV and young systemic leaves were collected at 12 dpi. n=10 plants. Total soluble proteins were extracted at a fixed fresh weight to buffer volume ratio and equal volume was loaded per well. Membranes were hybridized with the NIT antibody (@). Membranes were then stained with amidoblack to verify loading. See Figure S11D for a biological replicate. **(G)** Quantification of endogenous NIT protein signal in mock and TCV inoculated systemic leaves of Col-0 and *atg2* plants. 3-week-old rosettes were inoculated with buffer (mock) or with TCV and young systemic leaves were collected at12 dpi. Signal intensity is normalized to total protein signal and measured from biological replicates presented in Figure S11E. **(H)** Quantification of endogenous NIT protein signal in mock and TYMV inoculated systemic leaves of WT and *atg2* plants. 3-week-old rosettes were inoculated with buffer (mock) or with TYMV and young systemic leaves were collected at 12 dpi. Signal intensity is normalized to total protein signal and measured from biological replicates presented in Figure S11E. **(I)** Confocal microscopy images of *Arabidopsis* root expressing NIT1-Clover (green) under the HTR5 promoter and mCherry-ATG8A (magenta) under a ubiquitin 10 promoter in mock-inoculated or TCV-infected conditions at 7 dpi. Left panels: Seedlings were directly mounted in ½ MS for image acquisition. Each image is a maximum intensity projection of a full z-stack. Right panels: Seedlings were immerged in ½ MS with 1µM ConcanamycinA (ConA) for 3h to 4h then directly mounted in ½ MS for image acquisition. Each image is a single snap in the focal plane of the vacuole. Scale bar 10µm and 5µm in inset. **(J)** AIM residues contribute to the dimerization interface. Bottom. Schematic representation of the AlphaFold2 model for NIT1 dimer. NIT1 monomers are shown as ribbon diagrams (left, middle spanels) or as a surface representation (right panel). NIT1 monomers are coloured in green and tan, respectively, AIM residues are highlighted in red and residues in the dimerization interface are highlighted in blue. Top. Insets highlighting contacts of AIM residues in the dimer (left panel) and the position of AIM residues in the dimerization interface as a surface representation (right panel). **(K)** Size exclusion chromatography (SEC)-multiangle light scattering (MALS) analysis of NIT1. NIT1 elutes in two peaks, whereas the first peak yields a molecular weight (MW) of ∼800 kDa, corresponding to a NIT1 20mer. The second peak shows a shoulder and elutes several species with an average weight of 250 kDa. **(L)** *In vivo* NIT complexes observed by blue native PAGE (BN-PAGE) in mock inoculated and TCV-infected systemic leaves of Col-0, *atg2*, *atg5* and *atg11*. 3-week-old rosettes were inoculated with buffer (mock) or with TCV and young systemic leaves were collected at 12 dpi. n=10 plants. Total native complexes were extracted at a fixed fresh weight to buffer volume ratio and equal volume was loaded per well. Membranes were hybridized with the NIT antibody (@). Total quantity of NIT in the soluble extract is verified by SDS-PAGE and the membrane was then stained with amidoblack to verify loading. **(M)** Enrichment of proteins co-purified with NIT1-Clover represented by a volcano plot. Left panel: pairwise comparison of NIT1-Clover mock libraries to GFP control (all treatments) to reveal constitutive NIT1-Clover interaction partners. Middle panel: pairwise comparison of NIT1-Clover treated with TCV to NIT1-Clover mock treated to reveal TCV-inducible interacting partners of NIT1-Clover. Right panel: pairwise comparison of NIT1-Clover treated with TYMV to NIT1-Clover mock treated to reveal TYMV-inducible interacting partners of NIT1-Clover. The horizontal dashed line indicates the threshold above which proteins are significantly enriched (*p* value < 0.05, quasi-likelihood negative binomial generalized log-linear model) and the vertical dashed line the threshold for which proteins log2 fold change is above 1. **(N)** Left Panel: Protein abundance pattern represented by a heatmap (Log2(PSM+1) – meanPSM per protein) for the fourteen proteins identified as both specific and inducible interaction partners of NIT1-Clover and IMPDH1-Clover (at least two of four datasets). Right Panel: overlap between autophagy cargos identified by quantitative MS (grey circle) and identified candidate virus-induced partners of NIT1 and/or IMPDH1 (blue circle). **(O)** *In vivo* interaction of NIT1-Clover and IMPDH1-Clover with endogenous EDS1 protein. Systemic leaves expressing GFP control, NIT1-Clover and IMPDH1-Clover under HTR5 promoter in mock-inoculated, TYMV or TCV-infected conditions at 11 dpi in Col-0 background (n= 12-13 plants) or *atg5* (n=17-21 plants). Affinity purification was performed using @GFP beads. Immunoblot against the indicated antibodies (@) was used to detect bait and prey in the input and GFP purified fraction. Membrane corresponding to the input fraction was then stained with amidoblack to verify loading. * denotes Clover bait proteins detected with EDS1 antibody. **(P)** Immunoblot of endogenous EDS1 protein content in mock, TCV (left panel) and TYMV (right panel) inoculated systemic leaves of WT, *atg2*, *atg5* and *atg11* plants. 3-week-old rosettes were inoculated with buffer (mock), with TCV or TYMV and young systemic leaves were collected at 12 dpi. n=10 plants. Total soluble proteins were extracted at a fixed fresh weight to buffer volume ratio and equal volume was loaded per well. Membranes were hybridized with the EDS1 antibody (@). Membranes were then stained with amidoblack to verify loading. See Figure S14E for a biological replicate. **(Q)** Phenotypic characterization of *Arabidopsis* Col-0 (WT), *eds1*, *atg5*, *eds1/atg5* crispr (*atg5c*) mutant plants after inoculation with buffer (mock), with TCV, TYMV and TuMV-6k2-sc, 12 and 16 days-post-inoculation (dpi). See Figure S14G for additional genotypes.

How are both SARs conditionally degraded during infection? Remarkably, both NIT and IMPDH enzymes form catalytically active filaments ^50,51^, prompting us to speculate that the assembly state of these proteins might regulate their accessibility to the autophagy machinery. To test this hypothesis, we first generated an AlphaFold multimer model of the AtNIT1 dimeric structure in agreement with AtNIT4 cryoEM structure ^51^ (Figure S12A, B) and examined the residues involved in the dimerization interface. Our model revealed that Phe307, in position Θ of the validated FDSV AIM contacts Trp187 and Arg190 on the opposing protomer, while Ser309 further contacts Pro192 (Figure 5J). As the dimer is symmetric, the AIMs of both protomers are always occupied and thus inaccessible for interaction with ATG8 in any assembly configuration. Therefore, the only form of NIT1 capable of direct interaction with ATG8 is the monomeric form (Figure S12C). We then measured the stoichiometry of purified NIT1 using SEC-MALS (Figure 5K) and mass photometry (Figure S12D). Both approaches confirmed that NIT1 forms higher order assemblies of varying size. SEC-MALS indicated the presence of a 20mer and various smaller assemblies averaging at 250 kDa, while mass photometry indicated the presence of every species up to a 10mer, in dimeric increments. However, we could not detect a monomeric species, suggesting that while the filament size is dynamic, monomers are rare in normal conditions. We inferred that if only the monomeric NIT1 is targeted by autophagy and viral infection triggers monomerization of NIT1, we could detect the monomeric form in infected autophagy mutants. Consistently, Blue Native PAGE (BN-PAGE) analysis revealed the presence of low molecular weight species (below 146 kDa) only in TCV-infected autophagy mutants (Figure 5L). We conclude that most of NIT is sequestered in the filamentous form of the enzyme under normal growth conditions, preventing its association with ATG8. Upon infection, NIT filaments can disassemble and release monomeric NIT, which binds ATG8 with high affinity, leading to its recruitment to the autophagy machinery. Altogether, our results suggest NIT1 and IMPDH1 serve as SARs, determined by their oligomeric state.

We then set out to identify the cargo degraded by NIT1 and IMPDH. We conducted Clover AP in presence of TCV and TYMV for both NIT1-Clover and IMPDH1-Clover lines. Proteins significantly enriched by both baits in presence of the virus were considered for the following steps. For maximum specificity, we only considered candidates recovered using this method in at least two out of four AP sets (Figure S13A). AP of NIT1-Clover uncovered an interaction with NIT2/3, suggesting the existence of *in vivo* hetero-oligomers, and with IMPDH1/2 (Figure 5M, Table S13). AP of IMPDH1-Clover also recovered IMPDH2, suggesting mixed *in vivo* filaments (Figure S12B, Table S14), but not NIT proteins. ATG8A was detected in both baits (Figure 5M and Figure S13B). Both APs yielded ∼200 significant interactors. Of these, for IMPDH, 21 and 23 were enriched in TYMV and TCV infected samples, respectively. For NIT1, 31 were unique to TYMV, whereas 26 proteins specifically associated in TCV infected samples (Figure S13C). Applying our stringent filtering led to the identification of 14 core virus-induced interactors, amongst which 5 were identified as autophagy cargos in our proteomics experiments (Figure 5N and Figure S13D).

Of those 5 candidates, we focused on EDS1, as the infected *atg2* transcriptome shared similarities with EDS1-controlled genes (Figure S7C) ^45^, and EDS1 was detected in infected mCherry-ATG8E AP (Figure 4A). EDS1 senses pathogens by binding NAD+-derived small molecules produced by active TIR domain-containing NLR immune receptors ^53,54^, inducing transcriptional reprogramming ^55,56^ and oligomerization of pore-forming helper NLRs at the plasma membrane leading to cell death ^57–59^. CoIP experiments using the NIT1 and IMPDH1 reporter lines in WT and *atg5* background confirmed the interaction with the endogenous EDS1 (Figure 5O). Carbon starvation assays revealed that EDS1 is not a bulk autophagy cargo, but is sensitive to ConA treatment, confirming its vacuolar degradation (Figure S14A, B). In contrast, EDS1 level rose sharply after systemic invasion by TCV suggesting EDS1 activation (Figure S14C). Abundance of EDS1 appeared unchanged in TCV-infected *atg2* and *atg5*, but SDS-resistant high molecular weight products were enriched in the mutants (Figure S14D). We confirmed this observation in several autophagy mutants infected with TCV and TYMV (Figure 5P and Figure S14E), suggesting EDS1 is degraded in an autophagy-dependent manner during viral infection. Altogether, these results suggest NIT1 and IMDPH degrade EDS1 during viral infection to prevent autoimmunity.

To functionally test this, we mutated *ATG5* using CRISPR (*atg5c*) in *eds1* and *eds1*/EDS1-YFP backgrounds (Figure S14F) and compared their symptoms with the *atg5* mutant. Removal of EDS1 led to the complete or near-complete rescue of the *atg5* phenotype, as evidenced by the lack of necrotic lesions in the double mutant plants (Figure 5Q), while EDS1-YFP complementation resembled the autophagy mutants (Figure S14G). Collectively, these results unveil a novel selective autophagy mechanism that degrades major executors of cell death such as EDS1 to promote survival during viral infection.

## Discussion

In this study, we uncover a novel selective autophagy pathway that links organelle integrity with the proteolytic regulation of immunity and cell death. Contrary to what has been reported, for all three viruses that we study here, autophagy does not function as an antiviral response (Figure 1). Instead, autophagy promotes tolerance to infection and survival by clearing host proteins involved in cell death, defense and oxidative stress (Figures 4 and 5). Consistently, the autophagy degradome and the identified SARs are shared between TCV and TYMV infection, indicating a concerted response to these two viruses colonizing different organellar membranes. While the exact nature of the trigger remains to be uncovered, autophagy activation is intimately tied to organelle membrane integrity as expression of a single viral protein that remodels mitochondria is sufficient to recapitulate TCV infection (Figure 2). These findings are broadly relevant in virology since +ssRNA viruses co-opt endomembranes for their replication in host from all kingdoms of life.

We also identified two types of metabolic enzymes as SARs, Nitrilase and IMPDH. NIT hydrolyzes nitrile groups into acids via thiol-mediated catalysis ^60^ and IMPDH catalyzes the oxidative reaction of IMP to xanthosine 5′-monophosphate (XMP), the rate-limiting step in the synthesis of guanine nucleotides ^61^. These enzymes are found in prokaryotes and eukaryotes and, although they seemingly have nothing in common, both belong to a growing list of filament-forming enzymes whose activity is affected by their oligomeric state ^62,63^. We propose that both NIT1 and IMPDH1 moonlight as SARs to monitor organelle health in an oligomerization-dependent manner. Consistently, our findings suggest NIT1 monomers are released from the oligomeric filaments during infection to expose an AIM that mediates recruitment to the autophagosome (Figure 5). As the identified AIMs for both proteins are well conserved, this oligomerization-dependent organelle health monitoring system is likely to be of relevance to many other species.

Drawing inspiration from the plant immune system, we envision NIT and IMPDH as guards to organelle health, in which membrane rupture that arises during viral infection is monitored by leaked organelle content to the cytosol, which serves as DAMPs (Damage Associated Molecular Patterns) (Figure 6). In this model, the organelle integrity is the guardee and its modified-self is remotely monitored by oligomeric metabolic enzymes. NIT guarding function is conditioned by its oligomeric status and is in a self-repressed state in normal conditions, inaccessible to the autophagy machinery. This mirrors the self-repressed state of NLR proteins which are kept inactive by intramolecular interactions in the absence of their ligand ^64–66^. The release of a yet-to-be discovered DAMP to the cytosol changes the oligomeric status of the enzymes and converts them into SARs. However, in contrast with NLR-mediated immunity, SAR activation enables autophagic degradation of EDS1, ultimately limiting cell death and promoting the survival of infected cells. Interestingly, Nitrilase domain is co-opted by bacterial anti-phage defense systems and confers phage resistance ^67,68^, suggesting Nitrilase-based modified-self recognition could be an ancient tolerance mechanism.

**FIGURE 6:**
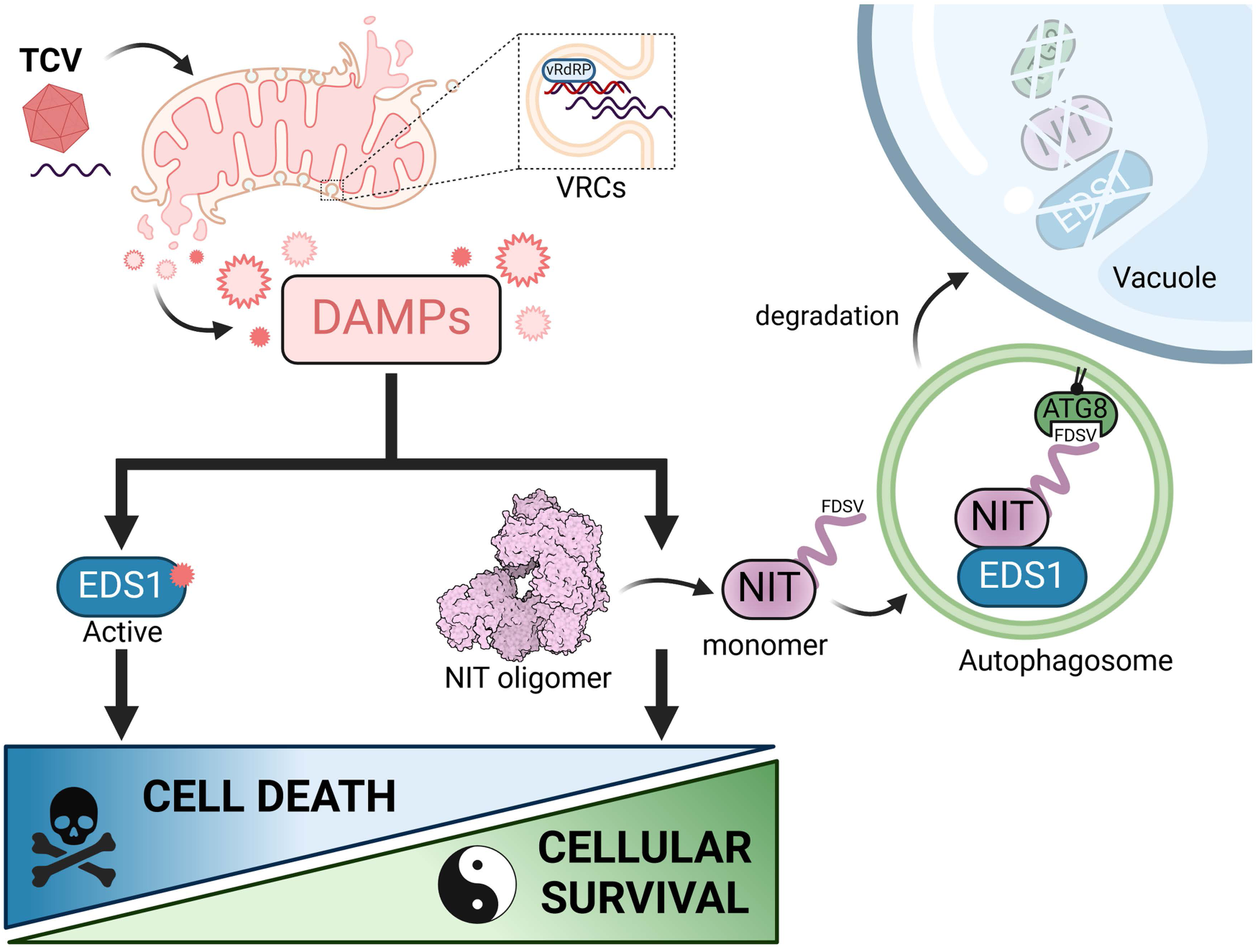
NIT-selective autophagy pathway safeguards the cell against excessive defense and cell death. TCV replication at the mitochondrial membranes causes membrane rupture and leakage of mitochondrial DAMPs into the cytosol of the infected cell. The plant intracellular immune system is activated by an unknown avirulence factor or directly by an organelle released DAMP. In parallel, DAMPs trigger the release of monomeric NIT1 from the oligomeric enzyme, and its association with ATG8 and EDS1. NIT1 and EDS1 are both recruited to autophagosomes and degraded in the vacuole. This novel proteostatic pathway promotes cellular survival, antagonizing defense and cell death by counteracting EDS1 activity.

Why would autophagy degrade EDS1 during viral infection? Plant viruses are obligate intracellular pathogens that largely move cell-to-cell symplastically. Should RNAi or hypersensitive response fail to contain the infection, the virus can outpace the immune system, establish VRCs in neighboring cells, and trigger cell death beyond the initial entry point. This could lead to a catastrophic amplification of immune signaling in distal tissues and lead to systemic necrosis ^69^, as seen with an engineered plant immune receptor recognizing TuMV ^70^ and akin to what we observe in autophagy mutants. By controlling the level of EDS1, selective autophagy prevents cell death when the cost outweighs the benefits and promotes organismal survival. In summary, NIT/IMPDH filaments represent a novel form of cellular health rheostats that monitor organelle integrity and balance immune responses to promote host survival during viral infection.

## Materials and methods

### Plant material and molecular cloning procedures

All *Arabidopsis thaliana* lines used in this study originate from the Columbia (Col-0) ecotype, with the exception of the *eds1-2* mutant, and are listed in Table S15. The primers used for genotyping are listed in Table S16. *atg2-2* ^71^, *atg5-1* ^72^, *atg1abct* ^73^, *atg4a-2/atg4b-2* ^74^, *atg9-3* ^75^, *atg11-1, atg11-1*/pUbi:GFP-ATG11 ^76^, *atg13a-2/atg13b-2* ^77^, *atg14a-1/atg14b-4* ^78^, *dcl2-1/dcl4-2* ^79^, *c53*, *ufl1*, pUbi:mCherry-ATG8A, pUbi:GFP-ATG8E, pUbi:GFP-ATG8H ^39^, *fmt*/mito-GFP ^80^, *nbr1-2* ^81^, *nit1-3* ^82^, *nit2* and *nit3-1* ^83^, Δnit1/2/3 ^84^, Col-0 *eds1-2* ^85^, Col-0 *eds1-2*/pEDS1:EDS1-YFP ^56^, pUbi:mCherry-ATG8E ^86^, p35S:B2-GFP ^87^, mito-GFP ^37^, mito-GFP/pUbi:mCherry-ATG8E ^20^, pUbi:HA-GFP-TOM5 ^38^, pIDH:IDH-GFP ^36^, pIDH:IDH-GFP/*atg2-1* ^88^ have been previously described. *Nicotiana benthamiana* expressing p35S:B2-GFP have been described previously ^25^.

Triples *dcl2-1/dcl4-2/atg2-2* and *dcl2-1/dcl4-2/atg5-1* were obtained by genetic crossing of the double *dcl* mutant to the respective *atg* mutant. F2 segregating populations were first screened using sulfadiazine and glufosinate resistance for *dcl2-2* and *atg5-1* T-DNA, respectively. Homozygous plants were obtained by genotyping PCR using the provided primers. *atg2-2* mutants were screened by Sanger sequencing of genomic DNA amplification using the provided primers. Descendants of isolated triple mutants were verified by genotyping PCR and autophagy deficiency was verified using carbon and nitrogen starvation assays. p35S:B2-GFP/pUbi:mCherry-ATG8A and p35S:B2-GFP/pUbi:mCherry-ATG8E were obtained by genetic crossing of the respective single mutants. Descendants containing pUbi:mCherry-ATG8A and pUbi:mCherry-ATG8E were selected on glufosinate and hygromycin-containing medium, respectively, and B2-GFP expression was verified using a stereomicroscope equipped with GFP filter. pIDH:IDH-GFP/pUbi:mCherry-ATG8A and pIDH:IDH-GFP/*atg5-1* were obtained by genetic crossing of the respective single plant line. F2 segregating populations for pIDH:IDH-GFP and pUbi:mCherry-ATG8E were selected on hygromycin and glufosinate-containing medium, respectively. F3 homozygous *atg5-1* were selected on glufosinate-containing medium and using genotyping PCR. Autophagy deficiency was verified by stabilization of NBR1 using western blot. pHTR5:NIT1-clover/*atg5-1* was obtained by genetic crossing of pHTR5:NIT1-clover/pUbi:mCherry-ATG8A to *atg5-1* plants. F2 segregating populations were screened for fluorescent yellow seeds using a stereomicroscope and for growth on glufosinate-containing medium followed by genotyping PCR. Autophagy deficiency was verified by stabilization of NBR1 using western blot. Note that the pUbi:mCherry-ATG8A transgene has been lost during segregation.

Stable Arabidopsis lines of pUbi:HA-GFP-TOM5/pUbi:mCherry-ATG8E lines were obtained by transformation of pUbi:mCherry-ATG8E plant line with pUTKan-3xHA-sGFP-TOM5 vector ^38^. Transformants were selected on Kanamycin-containing medium. pHTR5:NIT1-clover/pUbi:mCherry-ATG8A, pHTR5:NIT1-clover/pUbi:mCherry-ATG8E and pHTR-NIT1-clover/pUbi:mCherry-ATG8A Arabidopsis stable lines were obtained by transformation of pUbi:mCherry-ATG8A, pUbi:mCherry-ATG8E and pUbi:mCherry plants respectively, with pGGSun-pHTR5:NIT1-clover vector. Transformants were selected by picking fluorescent red seeds using a stereomicroscope. pHTR5:IMPDH1-clover/pUbi:mCherry-ATG8A, pHTR5:IMPDH1-clover/pUbi:mCherry-ATG8E, pHTR5:IMPDH1-clover/pUbi:mCherry and pHTR5:IMPDH1-clover/pUbi:mCherry-ATG8E/*atg5-1* Arabidopsis stable lines were obtained by transformation of pUbi:mCherry-ATG8A, pUbi:mCherry-ATG8E, pUbi:mCherry and pUbi:mCherry-ATG8E/*atg5-1* plants respectively, with pGGSun-pHTR5:IMPDH1-clover vector. Transformants were selected by picking fluorescent red seeds using a stereomicroscope. Mito-GFP/pUbi:mCherry-ATG8E/pUbi:mTurquoise2-myc and mito-GFP/pUbi:mCherry-ATG8E/pUbi:P29-mTurquoise2-myc Arabidopsis stable lines were obtained by transformation of mito-GFP/pUbi:mCherry-ATG8E plants with pGGSun-pUbi:mTurquoise2-myc and pUbi:P29-mTurquoise2-myc vectors respectively. Transformants were selected by picking fluorescent yellow seeds using a stereomicroscope.

Coding sequences from genes of interest were amplified from Col-0 cDNA with primers listed in Table S16 or obtained as synthetic DNA from Twist Biosciences (Table S16). All plasmids were assembled with the GreenGate (GG) cloning system ^89^ using the recommended protocol. For plant expression, all pieces were assembled in the pGGSun backbone ^90^ conferring kanamycin resistance. mTurquoise2 CDS was amplified from the pSW596 plasmid and converted to GG D modules. 3xMyc was added as a second step by PCR product stitching using BsaI restriction enzyme. The GG C module was then derived from the D module. The NIT1 (AT3G44310) and IMPDH1 (AT1G79470) CDSs were obtained as synthetic GG C modules and directly used for GG reactions. P29 sequence is derived from Melon necrotic spot virus isolate Mα5 (AY122286) ^91^. Introduction of point mutations was done from the entry modules with site directed mutagenesis using the primers listed in Table S16. IMPDH1 truncations were amplified from the WT module using the provided primers. IMPDH1 Δbateman was obtained by PCR product stitching and introduction of a short flexible linker region using BsaI restriction enzyme. His-MBP and mCherry B modules were modified to match the frame of the synthetic sequences using the primers listed in Table S16. All constructs used in this study are listed in Table S17. Plant expression plasmids were introduced into *Agrobacterium tumefaciens* strain GV3101 (C58 [RifR] Ti pMP90 [pTiC58DT-DNA] [gentR] Nopaline [pSoup-tetR]) and used for stable plant transformation using the floral-dip method ^92^ All plant lines are listed in Table S15.

### CRISPR/Cas9 mutant design and screening

Col-0 *eds1-2*/*atg5c* and Col-0 *eds1-2*/pEDS1:EDS1-YFP/*atg5c* were generated using a CRISPR/Cas9 approach. Two single guide RNAs (sgRNAs), each driven by an AtU6 promoter, were simultaneously expressed to target and remove a fragment of the gene of interest (AT5G17290). sgRNAs targeting the 5’ UTR region and the last exons were designed using CRISPR-P v2.0 (http://crispr.hzau.edu.cn/cgi-bin/CRISPR2/CRISPR) online tool ^93^ using the modular cloning system MoClo as previously described^94^. Two PCR products, each containing one guide RNA sequence, are amplified using plasmid pAGM9037 as a template with the primer pairs critarev/gRNA11_atg5 and critarev/gRNA21_atg5, respectively. Each PCR product was cloned with AtU6 promoter (pAGM38869) in MoClo Level 1 destination vector using BsaI-T4 ligase digestion/ligation reaction. Level 2 construct carrying two sgRNAs was generated using the selected level 1 constructs, a chosen end-linker (pICH41780), a yellow seed coat selection marker and a level 1 plasmid containing the Arabidopsis Rps5a promoter, Cas9 (Zcasi), and the nos terminator (pAGM51323). The abovementioned plasmids were cloned into L2 plant expression vector pAGM37663 via BpiI-T4 digestion-ligation reaction. The resulting vector was used to transform Col-0 *eds1-2*/*atg5* and Col-0 *eds1-2*/pEDS1:EDS1-YFP plants as indicated. Positive transformants seeds were selected and genotyping PCR was used to select *atg5* homozygous deletion. Sanger sequencing was used to identify the cut site. All primers used for cloning and genotyping are listed in Table S16.

### Virus material and inoculations

p35S:TuMV-6k2-scarlet infectious clone has been described ^90^. pBin p35S:TCV has been described ^79^. TYMV was obtained from Bioreba and directly used for Arabidopsis inoculation (Table S18). Sap was sourced from Col-0 infected systemic leaves for the different viruses and was verified using northern blot. Inoculum was prepared from a small amount of sap ground in 4 to 8 ml of 50 mM phosphate buffer pH 7.4, 0.2% sodium sulfite using a mortar and pestle. The inoculum was incubated at 4°C on a carousel and centrifuged 2 minutes at 2000 g. The resulting supernatant was kept on ice and three leaves of 21-day-old plants grown in 12 h light/12 h dark (120 μmol.m^-2^) were rub-inoculated using carborundum and q-tips. Excess inoculum was rinsed off and plants were shifted to 16 h light/8 h dark (110 μmol.m^-2^) growth conditions for the reminder of the experiment. Systemic leaves were collected at the indicated day post inoculation (dpi).

For inoculation of *in vitro* grown seedlings, the inoculum was sterile-filtered through a 0.2 µm filter with a syringe and the first true leaves of 9-day-old seedlings grown on vertical plates were rub-inoculated using two miniature q-tips under a laminar flow hood. Plates were resealed using leucopore and kept in identical growth conditions for the reminder of the experiment. Full seedlings bearing visible symptoms were collected for additional treatment or systemic roots were further processed for microscopy

### Plant growth and plant treatments

For standard plant growth, seeds were sown on water-saturated peat-based substrate (Klasmann Substrat 2, Klasmann-Deilmann GmbH, Gerste, DE) and supplemented with fertilizer (Wuxal). A solution containing *Bacillus thuringiensis* was applied as a preventative control against fungus gnat (Gnatrol, Valent BioSciences, Libertyville, Il, USA). Illumination was supplied by a custom multi-channel LED light co-developed with RHENAC GreenTec AG (Germany) which was optimized for uniformity. Plants were grown in 16 h light/8 h dark photoperiod with 165 µmol.m^-2^.s^-1^ light intensity (PPFD). using the following LED channels: 400nm, white_4500K, 660nm and 730nm, resulting in the following spectral light composition: 22% PPF-Blue (400-500nm) + 31% PPF-Green (500-600nm) + 47% PPF-Red (600-700nm). Far-red light (PPF-NIR; 700-780nm) was supplemented in an amount of 17% of PPFD (R:FR ratio = 1:0.35).

For *in vitro* seedling growth, Arabidopsis seeds were surface sterilized in 70% ethanol 0.05% SDS for 15 minutes, rinsed twice in ethanol absolute and dried on sterile paper. Seeds were plated in ½ MS salts (Duchefa)/1% agar/1% sucrose plates and stratified for 24-to-48 hours in the dark at 4°C. Plates were then grown under LEDs with 50 µmol.m^-2^.s^-1^ and a 16 h light/8 h dark photoperiod for the indicated amount of time. For *in vitro* infection experiments, ∼30 seeds per plate were sown on a single line and grown for the indicated time (typically 9 days) in vertical plates to let roots elongate on the medium surface in a growth cabinet equipped with neon lights for a 12 h light/12 h dark photoperiod.

For starvation phenotypical assays, ∼30 seeds were ethanol-sterilized and grown in 12-well plates containing liquid ½ MS salts (CaissonLabs), 1% sucrose medium. After stratification, plates were kept in in 16 h light/8 h dark photoperiod with 50 µmol.m^-2^.s^-1^ light intensity under constant gentle shaking. After 9 days, medium was drained and replaced with either liquid ½ MS salts, 1% sucrose medium (control +N) or Nitrogen deficient 1/2 MS medium (Murashige and Skoog salt without nitrogen [CaissonLabs] + Gamborg B5 vitamin mixture [Duchefa] supplemented with 0.5 g/liter MES and 1% sucrose, pH 5.7). Seedlings were rinsed twice in their respective medium and put back in the same culture conditions for 6 days before pictures were taken.

For carbon and nitrogen starvation and protein content analysis, seedlings were grown as indicated above for 7 days. Medium was drained and replaced with either liquid ½ MS salts (CaissonLabs) medium or Nitrogen deficient 1/2 MS medium (Murashige and Skoog salt without nitrogen [CaissonLabs] + Gamborg B5 vitamin mixture [Duchefa] supplemented with 0.5 g/liter MES and 1% sucrose, pH 5.7) in the case of Nitrogen starvation, or ½ MS salts (Duchefa)/1%sucrose or ½ MS without sucrose in the case of carbon starvation. For carbon starvation, +C plants were kept in regular growth conditions while -C plants were wrapped in foil to prevent photosynthesis. Plants were kept 24 h before samples processing. For autophagic flux experiments, 1 μM concanamycin A (conA; CAS 80890-47-7; Santa Cruz) was added to the relevant treatment condition and an equal volume of DMSO was added in the control condition. For measurement of steady state protein abundance, plants were treated for 24 h. For live imaging-based assays, treatment was restricted to a maximum of 6 hours. For pre-product analysis, mock-inoculated and TCV-infected seedlings were transferred to liquid ½ MS/1% sucrose medium supplemented with either 20 μM Oligomycin (Sigma-Aldrich O4876) or an equal amount of ethanol for 5.5 to 7 hours, under light condition. For depolarization experiments, mock-inoculated and TCV-infected seedlings were transferred to liquid ½ MS, 1% sucrose medium supplemented with 150 µM of FCCP (Sigma-Aldrich C2920) or an equivalent volume of DMSO for 10 to 60 minutes.

### Plant pictures

All plant pictures were acquired using a Canon EOS 80D DSLR equipped with either a 60mm fixed lenses or 18-135mm lenses, using manual setting in .CR2 format. Camera was mounted on a fixed stand and a dark cloth was used as background. Images were then opened using camera raw (adobe) and white balance was homogenized throughout all images. Pictures were exported in .tiff format and used to assemble panels.

### Automated phenotyping of infected plants

Pots measuring 65 x 65 x 60 mm (width, depth, height) were filled with 82 g of sieved peat-based substrate (Gramoflor Topf+Ton XL, Gramoflor GmbH & Co. KG, Vechta, DE.) Pots were placed within fixed grids and left to soak water for 24 hours. Seeds of the 14 indicated genotypes (Figure 1, second experiment) or the 10 indicated genotypes (Figure S1, first experiment) were surface sterilized and ∼10 seeds were sown per pot. The pots were covered with clean blue mats ^95^ to facilitate background subtraction for image analysis. Genotypes and treatment positions were randomized throughout the growth chamber to mitigate position specific effects, with the exception of the mock inoculated plants that were kept separate from infected plants that flowered later in the second experiment. Randomized pots were placed in a growth chamber and seeds were stratified directly on soil for 4 days at 4°C in the dark. Seeds were left to germinate and grow for at least 7 days and singularization of each pot was performed over a period of 5 days. Light intensity (110 μmol.m^-2^) and relative humidity were kept constant with 12 h light/12 h dark photoperiod during the initial phase of the experiment, using the following LED channels: 400nm, 440nm, white_3000K, 660nm and 730nm, resulting in the same spectral light composition as described above. Inoculation was performed on 3-week-old plants as indicated, and inoculated plants were placed back exactly in their assigned position and orientation. Care was taken to remove excess carborundum by gently spraying the rosettes with tap water to avoid unwanted white spots on the plants and mats. This was repeated the following day. For the first experiment, photoperiod was shifted to 16 h light/8 h dark to mirror usual growth condition during infection, but photoperiod was kept at 12 h light/12 h dark for the second experiment to delay flowering. During the pre-inoculation period, plants were photographed twice a day using an RGB camera (IDS uEye UI-548xRE-C; 5MP) mounted to a robotic arm. After inoculation, pictures were taken five times a day. Progression of symptoms was recorded during 30 days post inoculation (first experiment) and 28 days post inoculation (second experiment). Plants that flowered before the end of the kinetic were removed from the imaged pool. Images were exported from the phenotyping system in .png format with one plant present in each image. For the first experiment, images were read using the imread(). function from OpenCV. Due to the color variation in the infected plants, a 2-step threshold was used to segment the plants from the background. Firstly, pixel values between 0-154 in the b* channel of the La*b* color space were used to select green parts of the plant. A second threshold was applied in the YUV color channel with the values Y= 0-255, U = 90-255 and V = 150-255 to select non green plant parts. The 2 resulting binary masks were combined with the bitwise_or function in numpy. Plant masks were smoothed by up-sampling the mask, applying a median blur, down-sampling to the original dimensions and applying a binary threshold. Remaining noise was removed from the masks by excluding objects with a size of less than 300 pixels and performing an erosion operation with a kernel size of 3,3. To segment plants into healthy and symptomatic tissue, a threshold in the YUV color space was applied with the values Y = 0-150, U = 0-105, V = 0-160 to select the healthy green parts of the plants. The symptomatic areas were subsequently calculated by subtracting the mask of healthy tissue from the total plant mask. Total plant area and symptomatic area were calculated from the number of non-zero pixels in the two masks. For each plant, histogram values for each channel of the HSV color space were extracted for plant pixels using the calcHist() function of OpenCV. For the second experiment, camera sensitivity in the individual red green and blue channels were adjusted using a color checker (X-rite colorchecker Classic, X-Rite Europe GmbH, Regensdorf, CH) to compensate for the inhomogeneous color of the LED illumination. Images were read using the imread() function of OpenCV. Plants were masked from the background by blurring the image with a kernel size of 5,5 using the blur() function of OpenCV and then setting a threshold in the U channel of the YUV color space with values between 0-125 to select the blue background. Resulting masks were inverted to select the plants and noise removed by excluding objects less than 100 pixels in size. Virus symptoms were segmented from healthy plant tissue by applying a threshold in the HSV color space with values H = 1-35, S = 1 - 255, V = 1 - 255. Histogram values for plant pixels were extracted from plant pixels in the same way as the first experiment.

### Preparation of samples for confocal microscopy

For confocal microscopy of primary roots, Arabidopsis seedlings were placed in ½ MS medium or ½ MS medium containing 500 nM TMRE (resuspended in DMSO). After a few minutes in TMRE, stained seedlings were then rinsed in fresh ½ MS medium. After complete immersion and air bubble removal, whole seedlings were mounted on microscopy slides and a coverslip was placed on top of the roots and the epidermal cells of the elongation zone were used for image acquisition. For agro-infiltrated *N. benthamiana* leaves confocal microscopy, several leaf-disks were mounted as described previously ^96^.

### Confocal microscopy

Most confocal imaging was performed on an upright point laser scanning confocal microscope (Zeiss LSM780, Axio Imager, Carl Zeiss) equipped with a Plan-Apochromat 20x/0.8 dry objective, laser diode (405nm), Argon laser (488nm) and DPSS (561nm) laser and GaAsP (Gallium Arsenide) detector array (416-690 in 8.3 nm steps), driven by Zen black (Carl Zeiss). GFP and clover fluorescence were excited at 488 nm and detected between 499 nm and 550 nm with variations depending on the fluorophore and construct used. mCherry and TMRE were excited at 561 nm and detected between 580 nm and 630 nm with some variations depending on the fluorophore and construct used. mTurquoise2 was excited at 405 nm and detected between 410 nm and 490 nm. For each combination of fluorophores, one track was set at 1 Airy Unit (usually the marker corresponding to autophagosomes) and optimal sampling was achieved for each acquisition, following the Nyquist criterion. Signal-to-noise ratio was set with lowest gain possible and different laser power, averaging twice every line and 8.48E-07 seconds pixel dwell time. Fluorescent channels were kept on separate tracks and acquisition was done in sequential mode. For z-stack imaging, interval between the layers was set to 2 μm. For each experiment, all replicate images were acquired using identical confocal microscopic parameters and zoom factor. Confocal images were processed with Fiji and exported as .tiff for panel assembly. For acquisition of viral replication complexes presented in Figure S2A an upright point laser scanning confocal microscope (Zeiss LSM800, Axio Imager2, Carl Zeiss) equipped with an EC Plan-Neofluar 40x/1.30 Oil objective or a Plan-Apochromat 63x/1.40 Oil objective, GaAsP detectors (Gallium Arsenide) and driven with ZEN software (version 3.2, blue edition, Carl Zeiss) was employed. GFP fluorescence was excited at 488 nm and detected between 496 nm and 509 nm. TMRE was excited at 561 nm and detected between 587 nm and 610 nm. For acquisition of mito-GFP alongside TMRE staining, due to the movement of mitochondria, an inverted spinning disk (Yokogawa CSU-W1, Yokogawa) confocal microscope (OLYMPUS IX83, Olympus) equipped with a 60x UPLSAPO 1.20 water immersion objective and a dual-camera (Hamamatsu Orca Flash 4.0, Hamamatsu) controlled by CellSense Dimension software was used. Laser power was set at 70% and 100ms exposure time was used. GFP was excited at 488 nm and TMRE at 561 nm. The camera was used in standard mode (480 Hz), and no binning (pixel size of the camera 6.5 µm, image pixel size 108 nm). Single roots were imaged in z-stack mode and microscopy settings were identical between treatments. Channels were realigned using Huygens Professional software (SVI) before further processing and colocalization measurement, to correct for chromatic shift.

### Microscopy image processing and quantification

Quantification of autophagosomes was performed in Fiji using the sice spot detector macro ^97^. Maximum intensity projections of z-stacks (5 to 8 slices) were used as input with the following settings: size of objects was constrained between 0.4 and 99 µm, circularity between 0,7 and 1, difference of gaussian (DoG) filtering and triangle method for auto-thresholding. Best sigma value was determined empirically for each experiment and fixed for analysis. All images were analysed at once and measured area (region of interest, ROI) was manually defined for each and recorded. The raw number of autophagosomes (spots) was normalized to the normalized area and normalized slice number and expressed as autophagosomes per normalized z-stack. Measured values for all replicates were exported to R for plotting.

Quantification of autophagic bodies was performed in Fiji from single snap micrographs or single slices extracted from a z-stack. For each measurement, a single cell in the focal plane of the vacuole was selected per root and ROI was manually set to only select the vacuolar area. A threshold for detection was then applied using the MaxEntropy method and the number of puncta was counted using the analyse particle function of Fiji. Only particles above a size of 0.4 µm were considered. Number of particles was extracted for each channel and normalized as described above.

For colocalization measurement, Mander’s coefficient was used. Huygens-aligned spinning disk-acquired z-stacks were imported into Fiji and assembled as composites. ROIs with low movement and noise were manually picked for a single slice in the root z-stack (up to two individual slices were measured per root). Cropped ROI were used as input for JACoP plugin ^98^. M1 is the fraction of mito-GFP overlapping with TMRE and M2 the fraction of TMRE overlapping with mito-GFP. Thresholding was automated using Costes’. M1 and M2 values were collected for all four treatment combinations and exported to R for plotting.

### Transmission electron microscopy and dual-axis tomography in roots

The TEM assay was performed following a previously established method ^20,99^. Briefly, 4-d-old seedlings were germinated on 1/2 MS plate and dissected under a microscope before freezing. For high-pressure freezing, the root tips were collected and immediately frozen with a high-pressure freezer (EM ICE, Leica). For freeze substitution, the root tips were substituted with 2% osmium tetroxide in anhydrous acetone and maintained at -80°C for 24 hours using an AFS2 temperature-controlling system (Leica). Subsequently, the samples were subjected to three washes with precooled acetone and slowly warmed up to room temperature over a 60-h period before being embedded in EPON resin. After resin polymerization, samples were mounted and trimmed. For the ultrastructure studies, 100 nm thin sections were prepared using an ultramicrotome (EM UC7, Leica) and examined with a Hitachi H-7650 TEM (Hitachi-High Technologies) operated at 80 kV.

For dual-axis tomography analysis, semi-thick sections (250 nm) were collected on formvar-coated copper slot grids (Electron Microscopy Sciences) and stained with 2% uranyl acetate in 70% methanol followed by Reynold’s lead citrate. Electron tomography was performed with a 200 kV Tecnai F20 electron microscope (FEI Company) as described previously ^100^. The tilt image series were captured from -60° to 60° (1.5° intervals) and collected again after rotating the grid by 90°. Tomograms were generated with IMOD software (https://bio3d.colorado.edu/imod/) by calculating the stack pairs. The contours of mitochondria were drawn with the 3dmod program in IMOD ^101^.

### Transmission Electron microscopy in systemic leaves

Young systemic leaves (7dpi) were fixed for 16 hours in 4% (v/v) glutaraldehyde, postfixed in 0.1% (v/v) osmium tetroxide in 150 mM sodium phosphate buffer, pH 7.2 and stained in 2% (w/v) uranyl acetate. Samples were dehydrated through an ethanol series, infiltrated with EPON 812 medium grade resin (Electron Microscopy Sciences) and polymerized for 48 h at 60°C. Ultrathin sections (70 nm) were collected on grids coated with formvar (Electron Microscopy Sciences). Transmission electron microscopy was performed with a Hitachi H-7500 electron microscope at 80 kV. Images were captured with a CDD Advantage HR Hamamatsu camera and AMT software (Advanced Management Technology).

### Protein extraction and western blotting

Total denatured protein extraction and quantification was performed as described previously ^96^, with denaturation performed for 10 minutes at 70°C. The indicated total protein amount was loaded on SDS-PAGE gels (4-20% Mini-protean^®^ TGX™ precast gels, [Bio-Rad], NuPAGE 4-12% Bis-Tris gels [Thermofisher], or self-cast minigels) and blotted on PVDF Immobilon-P membrane (Millipore) using a wet transfer apparatus (Bio-Rad) in cold 25 mM Tris Base, 192 mM Glycine, 20% ethanol for 2 hours at 170 mA. Membranes were blocked in TBST (10 mM Tris-HCl pH 7.5, 150 mM NaCl, 0.05% Tween 20) +5% skimmed milk or in TBST +2.5% BSA and hybridized overnight with the indicated antibody in blocking buffer. All primary and secondary antibodies are provided in Table S19. Images were captured on an iBright Imaging System (Invitrogen) with SuperSignal West Pico PLUS Chemiluminescent Substrate (Thermo Fisher Scientific). Equivalent loading was verified on the membrane by staining the membrane in Amidoblack staining solution (0.1% naphthol blue black, 10% acetic acid).

For analysis of soluble mitochondria-derived autophagy cargos, total soluble proteins were extracted from frozen tissue powder (several plants per genotype∼treatment∼timepoint combination) in a 3:1 v/w ratio. Collected tissue was coarsely ground with a spatula in liquid nitrogen and a fixed amount of material was transferred into a 1.5 ml Eppendorf SafeLock tube with glass beads and further ground using a Silamat S6 (Ivoclar Vivadent) for 7 seconds. GTEN buffer (10% glycerol, 50 mM Tris-HCl [pH 7.5], 150 mM NaCl, 1 mM EDTA, 0.2% IGEPAL^®^ CA-630, 0.5 mM DTT, 1X protease inhibitor cocktail [Roche]). Crude extracts were incubated on ice for 10 minutes and centrifuged for 15 minutes twice at 14.000 g, 4°C. Cleared supernatant was transferred to a fresh tube and 1 volume of 4X Laemmli (200 mM Tris–HCl [pH 6.8], 8% [w/v] SDS, 40% [v/v] glycerol, 0.05%, [w/v] bromophenol blue, 3% [v/v] β-mercaptoethanol) was added. Proteins were denatured for 10 minutes at 70°C. SDS-PAGE and western blots were done as indicated above but equal volume were loaded instead of fixed protein amounts. For signal quantification, the plot lane function of ImageJ was used to obtain the raw intensity for a signal of interest as well as from the whole amidoblack stained lane. Each signal was subsequently normalized to total protein signal and the obtained values were imported into R for plotting. For quantification of GFP cleavage, signal intensity was quantified for both the intact fusion protein and the vacuolar free GFP/clover product and expressed as a ratio and were used directly for plotting.

### Blue-Native PAGE

To analyse native NIT complexes, frozen infected tissue was recovered and ground as described in the western blotting section. Tissue was resuspended in 2:1 v/w ratio in 25 mM Bis-Tris [pH 7], 20% Glycerol, 1 mM DTT, 1X EDTA-free cocktail inhibitor (Roche) and kept on ice for the whole procedure. Crude extracts were centrifuged for 15 minutes at 14.000 g, 4°C. Cleared supernatant was transferred to a fresh tube and 4X NativePAGE™ sample buffer (Invitrogen) was added to a final 1X concentration. Native proteins were kept on ice while half of the crude soluble extract was resuspended in 1 volume of 4X Laemmli buffer and denatured for 10 minutes at 70°C to control for total protein extracted by this method. Denatured samples were resolved by SDS-PAGE as described and native protein extracts were resolved in precast NativePAGE™ Novex 3–12% Bis-Tris Gels (Invitrogen) alongside 10 μl of Serva Native Marker. Gel pockets were cleaned and filled with cold Anode buffer (50 mM Bis-Tris, 50 mM Tricine [pH 6.8]) before carefully layering equal volume of samples in the pockets with gel loading tips. Lower part of the tank was filled with cold anode buffer and upper part with cold cathode buffer (50 mM Bis-Tris, 50 mM Tricine [pH 6.8], 0.002% Coomassie G-250) without disturbing gel pockets. The gel was left to run for ∼30 minutes at 100 V, then the voltage was increased to 150 V for about 1h and finally 250 V when current dropped below 6 mA, until migration front was at the bottom of the gel. Tank was surrounded with ice during the whole separation process. The gel was immediately denatured in 1% SDS, 50 mM Tris-HCl pH 7.5 for 20 minutes with gentle rocking, rinsed briefly in cold transfer buffer (25 mM Bicine, 25 mM Bis-Tris, 1 mM EDTA [pH 7.2], 20% methanol) and subjected to wet transfer for 1 h at 25 V with a cool block. Membrane was then fixed and destained in 25% methanol, 10% acetic acid for 15 minutes with gentle rocking, then rinsed twice in TBST, blocked in TBST + 5% skimmed milk and hybridized as for a western blot.

To analyse native virions of TCV and TYMV, the same protocol was used with the following differences: cathode buffer contained 0.02% Coomassie G-250, and after separation was complete gel was immersed in fixing solution (40% methanol, 10% acetic acid) and microwaved for 45 seconds, then incubated another 15 minutes with gentle rocking. The gel was then immersed in destaining buffer (8% acetic acid) and microwaved 45 seconds. The gel was incubated for as long as needed to obtain optimal signal-to-background ratio. The gel was rinsed and stored in water until imaged using the iBright imager.

### RNA extraction, northern blot and RTqPCR

RNA extraction was performed using TRI Reagent (Zymo Research) as described ^96^. Total RNA was resuspended in ultrapure water and quantified using a nanodrop. Formaldehyde-based RNA gels were performed as described ^102^ and glyoxal-based RNA-gels were performed as described ^90^. Separated RNA species were transferred onto Hybond-NX (Amersham) membrane overnight by capillarity and UV cross-linked the following morning. After staining with methylene blue to verify equal loading, membranes were destained and probed with [γ-^32^P]ATP-end-labeled DNA oligonucleotides (Table S16), with T4 polynucleotide kinase [ThermoFischer]) overnight at 50°C in 1 mM EDTA, 7% SDS, and 500 mM sodium phosphate pH 7.2. Membranes were washed 10 minutes twice in 2% SDS, 2× SSC and once in 1X SSC, 1% SDS at 55°C. Membranes were exposed to phosphor screen and signals detected with a Typhoon FLA 9000 imager.

For quantitative RT-PCR, 2-3 µg of total RNA were treated with DNase I (Thermo fisher), and half was reverse transcribed using the SuperScript™ IV First-Strand Synthesis System (Invitrogen) with 2.5 µM oligo d(T)_20_ primer. Quantitative PCR reactions were performed in 10 µl with Kapa SYBR Fast qPCR universal mix (Kapa Biosystems) with 100 nM of each PCR primer per reaction in a Roche LightCycler 96 with the following program: initial denaturation: 95°C 5 min. Cycling: 95°C 10 s, 60°C 15 s, 72°C 15 s single acquisition mode. Each cycle was repeated 45 times. Melting curve: 95°C 5 s, 55° C 1 min, 95°C in continuous acquisition mode with a ramp rate of 0.11°C*/*s, 5 acquisition per °C. All primers were designed with LightCycler Probe Design Software 2.0 with the following settings: minimum amplicon size 60, primer Tm 64°C, reaction conditions LC DNA master SYBR and are available in Table S16. For each cDNA technical duplicates or triplicates were obtained from the same plate and expression data was normalized using SAND (AT2G28390) as internal control. For both pair of primers, efficiency was calculated from a Col-0 TuMV-infected sample with cDNA serial dilutions. Relative quantification was obtained using the ΔΔCt method with the measured efficiencies, using an excel macro ^96^. ΔΔCt values were exported to R for plotting.

### Affinity purification coupled to mass spectrometry

For purification of mCherry, mCherry-ATG8A and mCherry-ATG8E, 7-11dpi mock-inoculated and TCV/TYMV infected young systemic leaves were used as starting material (n=40 plants per combination). Tissue mass was measured and ground in GTEN buffer (10% glycerol, 50 mM Tris-HCl [pH 7.5], 150mM NaCl, 1 mM EDTA, 0.2% IGEPAL^®^ CA-630, 0.5 mM DTT, 1X protease inhibitor cocktail [Roche]) in a 3:1 v/w ratio. For each sample, tissue was first ground to a fine powder with a pre-chilled mortar and pestle in liquid nitrogen. Chilled buffer was then directly added to the mortar and mixed into a fine paste. Crude extract was left to thaw in the mortar on ice for about 15 minutes while other samples were processed in the same manner. All samples were collected in 15ml tubes and incubated on a carousel for 15 minutes at 4°C. Samples were centrifuged in a swinging-bucket rotor for 15 minutes at maximum speed and cleared crude extract was pipetted into fresh tubes and kept on ice during total protein quantification. Quantification was obtained from 10 µl of crude extract using the amidoblack method ^96^ and crude extracts were normalized to an equal total protein concentration (between 1.7 µg/µl and 3.3 µg/µl). For each immune reaction an equal volume of crude extract was added to a fresh Maxymum Recovery tube (Axygen) corresponding to 3.28 mg for mCherry-ATG8A TCV experiment, 4.1 mg for mCherry-ATG8E experiment and 3.24 mg for both TYMV experiments. Each reaction was spiked with 150 µM AIM *wt* or AIM *mut* peptide ^39^ (stock peptide prepared at 5 mM in 50 mM phosphate buffer). Chromotek RFP-Trap magnetic agarose (Proteintech) bead slurry was equilibrated three times in 1 ml GTEN buffer on a carousel at 4°C. Slurry volume was set at 30 µl per reaction and the total volume of slurry for a given experiment was equilibrated in a single tube. Crude extract + AIM tubes were immobilized on magnetic stand and fixed volume of equilibrated beads were added to each tube. Binding reaction was performed on a carousel (12 rpm) for 1 hour at 4°C with care taken not to leave beads immobile at the bottom of the tube. Each immune reaction was then briefly centrifuged in a table top centrifuge and immobilized on a magnetic stand placed on ice. Crude extract was removed by pipetting and replaced with 0.9 ml GTEN buffer. All tubes were flipped upside down ∼5 times by hand, spun again in table top centrifuge and immobilized. This bead washing procedure was repeated three times with GTEN buffer. Each reaction was further washed in 10% glycerol, 50 mM Tris-HCl pH 7.5, 150 mM NaCl, 1 mM EDTA buffer (no detergent, no DTT, no protease inhibitor) for three more times by rotating the tubes on the magnetic rack 360°. Beads were resuspended in 200 µl of the same buffer and transferred to fresh Maxymum Recovery tubes. 20 µl (1/10^th^ of beads) was drained and denatured in 10 µl 2X Laemmli for quality checking each reaction by western blot. The rest was drained of excess buffer and flash frozen in liquid nitrogen before on-bead Trypsin digestion.

For purification of GFP, NIT1-clover and IMPDH1-clover, 11dpi mock-inoculated and TCV/TYMV infected young systemic leaves were used as starting material (n=30 plants per combination) and treated with the same protocol with the following exceptions: no AIM peptide was added, affinity resin used was Chromotek GFP-Trap^®^ magnetic agarose (Proteintech), each immune reaction contained 6.3 mg total protein in 3.8 ml crude extract contained in 5 ml LoBind Eppendorf tubes. After binding, beads were transferred to fresh 1.7 ml Axygen Maxymum Recovery tubes.

In both cases, beads were resuspended in 50 ul of 100 mM ammonium bicarbonate (ABC), supplemented with 400 ng of lysyl endopeptidase (Lys-C, Fujifilm Wako Pure Chemical Corporation) and incubated for 4 hours on a Thermo-shaker with 1200 rpm at 37°C. The supernatant was transferred to a fresh tube and reduced with 0.5 mM Tris 2-carboxyethyl phosphine hydrochloride (TCEP, Sigma) for 30 minutes at 60°C and alkylated in 3 mM methyl methanethiosulfonate (MMTS, Fluka) for 30 min at room temp protected from light. Subsequently, the sample was digested with 400 ng trypsin (Trypsin Gold, Promega) at 37°C overnight. The digest was acidified by addition of trifluoroacetic acid (TFA, Pierce) to 1%. A similar aliquot of each sample was analysed by LC-MS/MS.

### MS/MS Data acquisition and processing

The nano HPLC system (UltiMate 3000 RSLC nano system) was coupled to an Orbitrap Exploris^TM^ 480 mass spectrometer equipped with a FAIMS pro interface and a Nanospray Flex ion source (all parts Thermo Fisher Scientific).

Peptides were loaded onto a trap column (PepMap Acclaim C18, 5 mm × 300 μm ID, 5 μm particles, 100 Å pore size, Thermo Fisher Scientific) at a flow rate of 25 μl/min using 0.1% TFA as mobile phase. After loading, the trap column was switched in line with the analytical column (PepMap Acclaim C18, 500 mm × 75 μm ID, 2 μm, 100 Å, Thermo Fisher Scientific). Peptides were eluted using a flow rate of 230 nl/min, starting with the mobile phases 98% A (0.1% formic acid in water) and 2% B (80% acetonitrile, 0.1% formic acid) and linearly increasing to 35% B over the next 120 min. This was followed by a steep gradient to 95%B in 5 min, stayed there for 5 min and ramped down in 2 min to the starting conditions of 98% A and 2% B for equilibration at 30°C.

The Orbitrap Exploris 480 mass spectrometer was operated in data-dependent mode, performing a full scan (m/z range 350-1200, resolution 60.000, normalized AGC target 100%) at 3 different compensation voltages (CV-45, -60, -75), followed each by MS/MS scans of the most abundant ions for a cycle time of 0.9 (CV -45, -60) or 0.7 (CV -75) seconds per CV. MS/MS spectra were acquired using HCD collision energy of 30%, isolation width of 1.0 m/z, resolution of 30.000, max fill time 100ms, normalized AGC target of 200% and minimum intensity threshold of 2.5E4. Precursor ions selected for fragmentation (include charge state 2-6) were excluded for 45 s. The monoisotopic precursor selection (MIPS) filter and exclude isotopes feature were enabled.

For the measurement of the mCherry-ATG8E IP the method was as described above except the following changes: only 2 CVs (-45, -60) for 1sec cycle time each, a full scan AGC target of 300% and a minimum intensity of 1E4 were used. HCD energy was set to 28, resolution for MS2 was set to 45.000 and max inject time was 87ms.

For peptide identification, the RAW-files were loaded into Proteome Discoverer (version 2.5.0.400, Thermo Scientific). All MS/MS spectra were searched using MSAmanda v2.0.0.19924 ^103^. The peptide mass tolerance was set to ±10 ppm and fragment mass tolerance to ±10 ppm, the maximum number of missed cleavages was set to 2, using tryptic enzymatic specificity without proline restriction. Peptide and protein identification was performed in two steps. For an initial search the RAW-files were searched against the Arabidopsis database TAIR10 (32.785 sequences; 14.482.855 residues), supplemented with common contaminants and sequences of tagged proteins of interest using beta-methylthiolation on cysteine as a fixed modification. The result was filtered to 1 % FDR on protein level using the Percolator algorithm ^104^ integrated in Proteome Discoverer. A sub-database of proteins identified in this search was generated for further processing. For the second search, the RAW-files were searched against the created sub-database using the same settings as above and considering the following additional variable modifications: oxidation on methionine, deamidation on asparagine and glutamine, phosphorylation on serine, threonine and tyrosine, glutamine to pyro-glutamate conversion of peptide N-terminal glutamine and acetylation on protein N-terminus. The localization of the post-translational modification sites within the peptides was performed with the tool ptmRS ^105^, based on the tool phosphoRS. Identifications were filtered again to 1 % FDR on protein and PSM level, additionally an Amanda score cut-off of at least 150 was applied. Proteins were filtered to be identified by a minimum of 2 PSMs in at least 1 sample (or 3 peptides in at least 1 sample for the mCherry-ATG8E set). Protein areas have been computed in IMP-apQuant ^106^ by summing up unique and razor peptides. Resulting protein areas were normalized using iBAQ ^107^ and sum normalization was applied for normalization between samples (see Tables S20 to S23).

### Differential protein enrichment analysis

The total number of MS/MS fragmentation spectra was used to quantify each protein (Tables S21 to S27). The data matrix of Peptide-Spectrum Match (PSM) was analysed using the R package IPinquiry4 (https://github.com/hzuber67/IPinquiry4) that calculates log2 fold change and P values using the quasi-likelihood negative binomial generalized loglinear model implemented in the edgeR package ^108^. Only proteins identified with at least 3 PSM were considered. Each genotype∼treatment was triplicated per experiment, with the exception of the control mCherry and GFP IPs which were acquired as duplicates. For the first level of filtering (control *vs.* bait) all control reactions were pooled for the pairwise comparison irrespective of the treatment applied. pvalue and log2 fold change cutoffs applied to each pairwise comparison are described in Figure S5A for the mCherry-ATG8 IPs and in Figure S12A for the NIT1-clover and IMPDH1-clover IPs. Annotations were retrieved for each protein detected in a given experiment using TAIR bulk data retrieval gene description tool (https://www.arabidopsis.org/tools/bulk/genes/index.jsp). Venn diagrams were built using the Venny 2.1.0 online tool ^109^ and then redrawn manually. For candidate proteins abundance across all replicates was plotted in a heatmap as log2(PSM+1) and normalized to the mean value per protein. Rows were clustered using euclidean distance and resulting dendrograms are omitted from the figures.

### Sample preparation for parallel reaction monitoring and quantitative proteomics

For parallel reaction monitoring (PRM), samples were obtained from infected young systemic leaves (n=4-8 plants) at the indicated time after inoculation. For quantitative tandem mass tagging, samples were obtained from mock-inoculated and infected young systemic leaves (n=10 plants) at 12dpi. For each sample, tissue was coarsely ground with a spatula in liquid nitrogen and a fixed amount of material was transferred into a 1.5 ml Eppendorf Safe-Lock tube with glass beads and further ground using a Silamat S6 for 7 seconds. Samples were resuspended in 2 volumes of SDT buffer (4% SDS, 100 mM DTT, 100 mM Tris-HCl [pH7.5]). Proteins were denatured for 5 minutes at 95°C and centrifuged at 14.000 g for 5 minutes. Protein concentration was determined using either the amidoblack method or Pierce 660nm Protein assay kit following the manufacturer’s recommendation. Following steps were all done at room temperature unless otherwise indicated. Microcon-30 ultracel PL-30 (Millipore) centrifugal units were equilibrated in 100 µl of UA buffer (8 M Urea in 100 mM Tris-HCl [pH 8.0]) and centrifuged at 14.000 g for 10 minutes. 200 µl of UA buffer was then loaded onto the filter and 30 µl of sample was mixed into the UA buffer by gentle pipetting. Filter units were centrifuged for 20 minutes at 14.000 g until most liquid passed through the filter. Proteins retained on filter were equilibrated in 200 µl using the same method. 100 µl of fresh alkylation buffer (100 mM Iodoacetamide, 8 M Urea in 100 mM Tris-HCl [pH 8.0]) was added on top of the filter and samples were incubated for 20 minutes in the dark. Filters were centrifuged for 15 minutes at 14.000 g and washed in 100 µl UA buffer with the same method for a total of three times. Filters were equilibrated in 100 mM HEPES pH7.6 for three consecutive rounds using the same method. 3 µl of Trypsin gold (Promega, resuspended at 1 µg/µl in 50 mM Acetic acid) was added to 47 µl of 100 mM HEPES pH7.6 directly to the filter and mixed by pipetting and gentle flicking of the collection tube. Filters were sealed with Parafilm and incubated overnight at 37°C. Filter units were transferred to a fresh collection tube and peptides were collected by centrifugation, 10 minutes at 14.000 g. Filters were washed again in 50 µl of 100 mM HEPES pH 7.6 using the same method and both eluates were collected in the same tube. Final peptide amount was determined by separating an aliquot of the sample on a capillary-flow LC-UV system using a PepSwift^TM^ Monolithic column ^110^ based on the peak area of 100 ng of Pierce HeLa protein digest standard (PN 88329; ThermoFisher Scientific). Peptide solution was frozen at -80°C before further processing.

### Targeted mass spectrometry via parallel reaction monitoring (PRM)

The nano HPLC system and settings were the same as described above, coupled to a Q-Extractive HF-X spectrometer. The spectrometer was operated by a mixed MS method which consisted of one full scan (*m/z* range 350-1.500; 15.000 resolution; target value 1E6) followed by the scheduled PRM of targeted peptides from an inclusion list (isolation window 0.8 *m/z*; normalized collision energy (NCE) 32; 30.000 resolution, AGC target 2E5, loopN 8). The maximum injection time was set to 550 ms. A scheduled PRM method (sPRM) development, data processing and manual evaluation of results were performed in Skyline (64-bit, v22.2.0.351) ^111^. Candidate peptides for viral ORFs and for reference proteins were chosen based on shotgun analysis of two timepoints for all viruses used. Seven internal reference genes were chosen with the following criteria: >45KDa, detected with at least 5 unique peptides with normalized intensity in the 1E09 range. Reference proteins are AT5G17920, AT4G13940, AT1G56070, AT1G57720, AT1G79440, AT5G08680, AT3G45140. For each viral ORF, the average area was calculated, normalized to the average area of all reference proteins and expressed relative to the value in first time point of the Col-0 sample. Obtained values were exported to R for plotting and are represented as stacked histograms. Mass spectrometry relevant details about all proteins and their specific peptides included in the sPRM method are stated in Table S1.

### TMT-labelling, separation and LC-MS of labelled peptides

100 µg of peptides (in 100 µl 100 mM HEPES pH 7.6) were labelled with one separate channel of the TMTpro™ 16plex Label Reagent Set (Thermo Fischer Scientific), according to the manufacturer’s description. The labelling efficiency was determined by LC-MS. Samples were mixed in equimolar amounts and equimolarity was again evaluated by LC-MS/MS. The mixed sample was acidified to a pH below 2 with 10% TFA and was desalted using C18 cartridges (Sep-Pak Vac 1cc [50 mg], Waters). Peptides were eluted with 3 x 150 µl 80% Acetonitrile (ACN) and 0.1% Formic Acid (FA), followed by freeze-drying.

Separation of TMT-labelled peptides was achieved by strong cation exchange chromatography (SCX). The dried sample was dissolved in 70 µl of SCX buffer A (5 mM NaH_2_PO_4_, pH 2.7, 15% ACN) and 200 µg of peptides were loaded at a flow rate of 35 µl/min on a TSKgel SP-2SW column (1 mm ID x 300 mm, particle size 5 µm, TOSOH) on an UltiMate 3000 RSLC nano system (Thermo Fisher Scientific) at a flow rate of 35 µl/min. For the separation, a ternary gradient was used. Starting with 100% buffer A for 10 min, followed by a linear increase to 10% buffer B (5 mM NaH_2_PO_4_, pH 2.7, 1M NaCl, 15% ACN) and 50% buffer C (5 mM Na_2_HPO_4_, pH 6, 15% ACN) in 10 min, to 25% buffer B and 50% buffer C in 10 min, 50% buffer B and 50% buffer C in 5 min and an isocratic elution for further 15 min. The flow-through was collected as a single fraction, then 60 fractions were collected every minute along the gradient. Fractions with low peptide content were pooled. ACN was removed by vacuum centrifugation and the samples were acidified with 0.1% TFA and analysed by LC-MS/MS.

The nano HPLC system and settings were the same as described in AP-MS/MS section, except that the gradient was increased linearly to 40% buffer B. The Orbitrap Exploris^TM^ 480 mass spectrometer was operated in data-dependent mode, performing a full scan (m/z range 350-1200, resolution 60.000, target value 100%) at 3 different compensation voltages (CV -40, -55, -70), followed each by MS/MS scans of the most abundant ions for a cycle time of 1 second per CV. MS/MS spectra were acquired using an HCD collision energy of 34%, isolation width of 0.7 m/z, resolution of 45.000, first fixed mass 110, target value at 200%, minimum intensity of 2.5E4 and maximum injection time of 120 ms. Precursor ions selected for fragmentation (include charge state 2-6) were excluded for 45 s. The monoisotopic precursor selection (MIPS) mode was set to Peptide, the exclude isotopes and single charge state per precursor feature were enabled.

Peptide and protein identification was performed as described in the AP-MS/MS section but using only one search step and considering the following modifications: Iodoacetamide derivative on cysteine and TMTpro-16plex tandem mass tag on peptide N-Terminus were set as fixed modification, deamidation on asparagine and glutamine, oxidation on methionine, carbamylation on lysine, TMTpro-16plex tandem mass tag on lysine as well as carbamylation on peptide N-Terminus, glutamine to pyro-glutamate conversion at peptide N-Terminus and acetylation on protein N-Terminus were set as variable modifications. Results were filtered as described for IPs including an additional minimum PSM-count of 2 across all fractions. Peptides were quantified based on Reporter Ion intensities extracted by the “Reporter Ions Quantifier”-node implemented in Proteome Discoverer. Proteins were quantified by summing unique and razor peptides and filtered for quantification by at least 3 peptides. Protein-abundance-normalization was done using sum normalization. Statistical significance of differentially expressed proteins was determined using limma ^112^.

For clustering analysis, protein quantification was imported in R (version 4.3.0). Total reporter abundances per protein were normalized to 100% for each protein to eliminate effects from proteins of different abundance. Next, proteins regulated less than 2-fold were removed before clustering. Protein abundance patterns across conditions of the remaining proteins were subjected to k-means clustering ^113^ and clustered to an arbitrary number of 20 clusters, and resulted in the clustering of 2833 proteins (out of 9586 initially considered). Normalized protein abundances are presented in Table S28, and clustering results in Table S8.

To account for transcriptional upregulation affecting the proteomic readout and assignment as a virus-induced autophagy cargo, the log2 ratio of protein abundance to transcript abundance (log2 PROT/RNA, where PROT is the mean normalized protein abundance and RNA is the mean TPM of replicates for a genotype∼treatment category) was used as proxy for assessing reliance on transcription for protein abundance (X axis). Increased reliance on autophagy at the protein level was assessed by plotting log2 (TCV/mock) in *atg2* – log2 (TCV/mock) in Col-0 in which ratios are calculated from mean normalized protein abundance for replicates for a genotype∼treatment category (Y axis). Data points were then separated in quadrants by calculating the centroid for both axes. Proteins of interest were then annotated and autophagy cargos assigned on the basis of clustering are annotated in blue. Pair plots showing the distribution of data points along both axes were generated using GGally and are presented alongside the relevant axis.

### GO terms enrichment analysis

The 735 autophagy cargos identified by k-means clustering of the TMT dataset were separated into two groups based on their presence or absence amongst the DEGs. GO terms enrichment was then performed on those two separate lists using DavidGO ^114^. Gene lists were used for the functional annotation chart tool, with count set to 5. -Log10(pvalue) and fold enrichment outputted by the tool were plotted for the top 12 BP (biological processes) using ggplot2.

### Co-Immunoprecipitation

For GFP-ATG8E and GFP-ATG8H enrichment, samples were obtained from young systemic infected leaves at the indicated timepoint (n=3 to 4 plants). Samples were processed as described in the affinity purification-MS/MS section with the following exceptions: total amount of frozen sample per reaction was fixed between 100-250 mg in 1.5 ml Eppendorf Safe-Lock tubes and were resuspended in 4 volumes of GTEN, total protein concentration was not measured but identical volume of crude extract was used per immune reaction, 50 µl of in-house GFP agarose affinity matrix was used per reaction and after binding beads were washed twice in GTEN and purified complexes were eluted and denatured in 2X Laemmli, 10 minutes at 70°C.

For GFP, NIT1-clover and IMPDH1-clover coIPs, samples were obtained from infected young systemic leaves (n=30 to 50 plants for mCherry-ATG8 experiments and n=12-15 plants for NIT1-clover and IMPDH1-clover experiments) at 11dpi. Samples were processed as described in the affinity purification-MS/MS section with the following exceptions: total amount of frozen sample per reaction was fixed between 250-400 mg in 1.5 ml Eppendorf Safe-Lock tubes, total protein concentration was not measured but identical volume of crude extract was used per immune reaction, 25 µl of bead slurry was used per reaction and after binding beads were washed 3 times in GTEN and purified complexes were eluted and denatured in 2X Laemmli, 10 minutes at 70°C.

### Libraries preparation and RNAseq analysis

The same samples as the ones analysed by TMT were used to generate 3’ end sequencing libraries, as was done previously ^115^, with minor modifications. Total RNA was extracted with a high-performance RNA bead isolation kit (Molecular Biology Service, VBC Core Facilities, Vienna), using a KingFisher Duo Prime System (Thermo Fisher Scientific). RNA quantity was determined with a Fluorometer Qubit Flex (Invitrogen) and Qubit RNA BR Kit (Invitrogen). DNAse treated total RNA was diluted in nuclease-free water to 20 ng/μl in 4 μl. Next, 1 μl of barcoded RT primer (GACGTGTGCTCTTCCGATCT<7 bp barcode>NNNNNNNNTTTTTTTTTTTTTTTTTTTTN, 0.2μM) was added, heated to 72°C, 3 min and immediately moved to -20°C. RNA-primer was then added to the Reverse Transcriptase (RT) mix consisting of SmartScribe RT (1 μl), 25 mM dNTP mix (0.7 μl), 20 mM DTT (1 μl), 100 mM MnCl2 (0.3 μl), and 5x SmartScribe buffer added to the RNA-primer mix. The RT reaction was conducted at 42°C for 1 hour, followed by 70°C for 15 minutes. Samples with different barcodes were pooled and purified using SPRI beads with addition of SPRI buffer (20% PEG, 2.5 M NaCl, adjusted to pH 5.5-6.0) to dilute the beads. Next, Tn5 was used to cut the cDNA: the DNA-RNA substrate was mixed with Tn5 activation buffer (20 mM Tris, 10 mM MgCl2, 20% dimethylformamide, pH 7.6) and loaded Tn5 enzyme (TCGTCGGCAGCGTCAGATGTGTATAAGAGACAG + [phos]CTGTCTCTTATACACATCT). The reaction was incubated at 55°C for 8 minutes, followed by deactivation and removal of Tn5 with 0.2% SDS at room temperature. PCR amplification was performed using KAPA HotStart ReadyMix (with primers: AATGATACGGCGACCACCGAGATCTACAC<8 bp barcode>TCGTCGGCAGCGTCAGATGTG and CAAGCAGAAGACGGCATACGAGAT<8 bp barcode>GTGACTGGAGTTCAGACGTGTGCTCTTCCGATCT). The amplification protocol included an initial denaturation at 95°C for 90 seconds, followed by 12 cycles of denaturation, annealing, and extension, with a final extension at 72°C for 5 minutes. The PCR products were purified using a 0.8x SPRI cleanup protocol and eluted in Elution Buffer. Libraries were verified using Fragment Analyzer analysis and quantified using qPCR with P5/P7 primers. Libraries were sequenced on a NovaSeq 6000 by the Next Generation Sequencing Facility at Vienna BioCenter Core Facilities (VBCF), member of the Vienna BioCenter (VBC), Austria.

Raw FASTQ files were quality filtered and adapter trimmed using Trim Galore with default parameters. Subsequently, trimmed reads were aligned to a genome index generated from the TAIR10 genome fasta file and all transcripts in the GTF of Ensembl build TAIR10 annotation set (release version 56) using STAR ^116^. The transcriptome bam files generated were then utilized for read count quantification per gene using Salmon^117^ in the alignment-based mode. Differential gene expression analysis was then carried out using DESeq2 ^118^, for genes with a minimum of five aligned reads. Genes with ≥ 2 log2-fold differences and an adjusted p-value of ≤ 0.01 were subsequently identified as differentially expressed genes (DEGs) for the following pairwise comparisons: Col-0 TCV vs. Col-0 mock, Col-0 TYMV vs, Col-0 mock, *atg2-2* TCV *vs. atg2-2* mock and *atg2-2* TYMV *vs. atg2-2* mock (Tables S9 to S12). Genes present in at least one of the above comparisons were merged and used as query to build a heatmap. Expression was plotted as log2(TPM+1) and normalized to the mean value per gene. Rows and columns were clustered using euclidean distance and resulting dendrograms are omitted from the figure. Expression values are clipped between -5 and 5. For assigning expression clusters for the included genes, the complete method of hclust was used and membership was distributed within 7 clusters using the cutree function and displayed as annotation. Genes defined as autophagy cargos are also annotated.

For analysis of significant DEGs isolated in previous studies, a gene list of interest was compiled from the supplemental material provided by the authors. Conditions are the following: flg22 treatment in *Arabidopsis thaliana*, *Capsella rubella*, *Eutrema salsugineum* and *Cardamine hirsute.* 868 genes that are counted as DEGs in all species ^44^ were used as starting query list to build a heatmap of expression in the dataset generated in this study (717 genes present in the dataset). *Pseudomonas syringae* pv. Tomato DC3000 AvrRpm1 inoculation of Arabidopsis and sampling within the inoculation zone (IN) or adjacent leaf tissue (OUT). 745 DEGs were recovered from the gene list provided by the authors ^43^ that fulfill the following criteria: Genes induced at both 4hpi and 6hpi in the IN section and excluded of the OUT sections at the same timepoints are defined as IN-specific genes. Genes induced at 4hpi, 6hpi and overlapping in the OUT section and excluded of the IN section at the same timepoints are defined as OUT-specific genes. Both lists were used as starting query to build heatmaps of expression in the dataset generated in this study (465 genes present for IN gene set, 218 genes present for OUT gene set). β-estradiol treatment in Arabidopsis *eds1-2/pad4-2* lines expressing 35S:EDS1-HA and inducible HA-PAD4. Upregulated genes list is provided by the authors ^45^ and fulfill the following criteria: induction at 12 h, 24 h and overlapping after start of the treatment (471 genes). These were used as starting query list to build a heatmap of expression in the dataset generated in this study (425 genes present in the dataset).

### Protein expression and purification for biochemical assays

For recombinant expression and purification of NIT1 in *E. coli*, NIT1 and IMPDH1 (and their derivatives) (Table S17) were cloned with an N-terminal 6XHis-MBP solubility tag in a modified pET vector ^39^ via GreenGate cloning ^89^. Recombinant proteins were produced using *E. coli* strain Rosetta2(DE3) pLysS grown in 2×TY medium or Terrific Broth (TB) at 37°C to an OD_600_ of 0.6-0.8 followed by induction with 500 µM IPTG and overnight incubation at 18°C.

For 6XHis-MBP-NIT1 WT and AIM mutants, protein purification strategy was adapted from established protocols ^119^. in brief, cells were harvested by centrifugation and re-suspended in 20 mM HEPES pH 7.6, 500 mM NaCl, 5% (v/v) glycerol, 1mM DTT and 20 mM imidazole supplemented with 1 tablet of cOmplete™ EDTA-free protease inhibitor (Roche) per every 50 ml. Cells were disrupted by sonication (VibraCell VCX750, 5 min, 50% amplitude) and, following centrifugation at 48.000 g for at least 30 min, the clarified lysate was applied to a HisTrap^TM^ Ni2+-NTA column connected to an ӒKTA Pure system (Cytiva Life Sciences).

Proteins were step-eluted with elution buffer (20 mM HEPES pH 7.6, 500 mM NaCl, 5% (v/v) glycerol, 500 mM imidazole) and directly injected onto a HiLoad^®^ 16/600 Superdex^®^ 200 pg gel filtration column (Cytiva Life Sciences) pre-equilibrated with 20 mM HEPES pH 7.6, 250 mM NaCl and 5% (v/v) glycerol. Elution fractions were collected and evaluated by SDS-PAGE.

Fractions containing 6XHis-MBP-NIT1 were then combined and incubated overnight at 4 °C with 6XHis tagged PreScission protease (Molecular Biology Service, VBC Core Facilities, Vienna) at 1:50 ratio with 2 mM DTT. MBP solubility tag was then separated by passing the protein mixture solution through a HisTrap^TM^ Ni2+-NTA column equilibrated with 20 mM HEPES pH 7.6, 500 mM NaCl, 5% (v/v) glycerol and 20 mM imidazole. Protein of interests were collected in the flow-through and washes from the column while the 6XHis-MBP tag and the 6XHis-PreScission protease was retained until the final elution with elution buffer. The NIT1 proteins were pooled together and concentrated for further purification and buffer exchange by gel filtration onto a HiLoad^®^ 16/600 Superdex^®^ 200 pg gel filtration column (Cytiva Life Sciences) pre-equilibrated with 20 mM HEPES pH 7.6, 250 mM NaCl. Fractions containing purified NIT1 proteins were combined and concentrated using Vivaspin^®^ Centrifugal Concentrators (Sartorius) as appropriate for further biophysical studies. Protein concentration was calculated from the UV absorption at 280 nm by DS-11 FX+ Spectrophotometer (DeNovix). NIT1 WT proteins used in GST-pulldowns presented in Figure S8I were purified following the same strategy but solutions were buffered with 50 mM Tris-HCl pH 7.5 instead of 20 mM HEPES.

For 6XHis-GST-ATG8A (used in pulldowns) and 6XHis-ATG8A (used in MST), we used previously established protocols ^39,120^. Eluted fractions were buffer exchanged to 10 mM HEPES pH 7.5, 100 mM NaCl and loaded either on a RESOURCE S cation exchange chromatography column (Cytiva Life Sciences). 6XHis-GST-ATG8A and 6XHis-ATG8A were gradient eluted in 5 to 55% of Ion exchange buffer (10 mM HEPES pH 7.5, 1 M NaCl by) in 20 column volumes. Finally, the proteins were separated by size-exclusion chromatography with HiLoad 16/600 Superdex 75 pg equilibrated in 25 mM HEPES pH 7.5, 250 mM NaCl.

### GST Pulldown

For GST-pulldown in cell lysates, 6XHis-GST-ATG8A to His-MBP-IMPDH1 and derivatives, *E. coli* pellets were resuspended in 50 ml of 100 mM sodium phosphate pH 7.5, 5% glycerol, 300 mM NaCl and 1 cOmplete™ EDTA-free protease inhibitor tablet (Roche). 7ml of resuspended cells were sonicated twice for 1 min (30 seconds ON, 30 second OFF) 20% amplitude (VibraCell VCX750) and lysates were then centrifuged for 20 minutes at 20.000 g 4°C in 5 ml Eppendorf tubes. Clarified lysates of GST and MBP constructs were mixed in a resulting 1 ml and IGEPAL^®^ CA-630 and DTT were added to a final concentration of 0.1% and 1 mM, respectively. 5 µl of equilibrated GST bead slurry equivalent (Pierce^TM^ Glutathione magnetic agarose beads or Cytiva glutathione sepharose 4B) and 200 µM AIM *wt* or mutant peptide were added to each tube. Mixtures were incubated for 1 hour at 4°C on a carousel and beads were washed five times with 1 ml 100 mM sodium phosphate pH7.5, 5% glycerol, 300 mM NaCl, 0.1% IGEPAL^®^ CA-630 and 1 mM DTT. Bound proteins were eluted and denatured in 50 µl 2X Laemmli + 100 mM DTT for 5 minutes at 95°C. Samples were analysed by western blot or Coomassie staining. Blotting was performed on nitrocellulose membrane using a semi-dry Trans-Blot^®^ Turbo™ Transfer System (Bio-Rad). Membranes were then incubated with appropriated antisera and revealed as described above.

For in vitro GST pulldowns using purified 6XHis-GST-ATG8A and NIT1 WT or mutants, protein dilutions, incubations and washing steps were performed in 50mM HEPES pH 7.6, 150 mM NaCl, 0.01 % (v/v) IGEPAL^®^ CA-630, 1 mM DTT (Figure 4Q) or 50 mM Tris-HCl pH 7.5, 150 mM NaCl, 5% (v/v) glycerol and 0.01 % (v/v) IGEPAL^®^ CA-630, 1mM DTT (Figure S8I). In brief, 10 µl of bead slurry (Pierce Glutathione magnetic agarose beads) were incubated with 5 µM of purified GST or 6XHis-GST-AtATG8A for 1 hour at 4 °C. After incubation, the beads were washed with buffer twice and mixed with 5 µM of NIT1 WT or mutants (Figure 4Q) or at the indicated concentrations (Figure S8I). If indicated, AIM *wt* or AIM mutant peptides were added to the reaction to a concentration of 200 µM. The beads were washed again 5 times, following an incubation of 1 hour at 4 °C. Finally, the bound proteins were eluted with 2x Laemmli at room temperature, loaded on an SDS-PAGE and analysed by Coomassie staining.

### MST – Fluorescence

To measure affinity between NIT1 and 6XHis-ATG8A, NIT1 WT and AIM2 mutant were labeled with RED-NHS 2^nd^ Generation dye (NanoTemper Technologies, München, Germany), following the manufacturer’s protocol. A serial dilution of ATG8A (100 µM – 3 nM) in MST Buffer (20 mM HEPES pH 7.5, 250 mM NaCl, 0.005% Tween-20) was mixed 1:1 with RED-NHS labeled NIT1 WT (200 nM or 50 nM) or NIT1 AIM2 (200 nM) and loaded into standard (WT) or premium (AIM2) glass capillaries. Analysis was performed on a Monolith NT.115 (NanoTemper) instrument over 20 s, 40 % MST power, 50 % (NIT1 WT) or 75 % (NIT1 AIM2) LED power at 25 °C. The initial fluorescence was analysed by MO.Affinity Analysis v2.3 and Palmist ^121^ to obtain Kd values for binding and corresponding confidence intervals.

### SEC-MALS

SEC-MALS was performed using a Superose^TM^ 6 increase 10/300 GL (Cytiva Life Sciences) operated at RT on a Malvern OmniSEC System. The OmniSEC system consists of a pump, an autosampler, a column oven and the following detectors: refractive index (RI), right angle light scattering (RALS), low angle light scattering (LALS), a photodiode array (UV detection) and an intrinsic viscosity (viscometer) detector. The sample was stored at 10°C in the autosampler prior to injection. Each run (flow rate of 0.5 mL/min) takes ∼55 min. The sample was measured in triplicate for NIT1 WT (0.5 mg/mL) and only one injection is shown in the figure. To calibrate and measure the variability of the system, BSA was injected in duplicate at the start of the measurement and prior to the replicates.

### Mass Photometry

Measurements were performed as outlined ^122^. Briefly, OneMP mass photometer (Refeyn) was calibrated using NativeMark^TM^ unstained protein standard (Invitrogen) as a protein standard using a regular field of view. Mass Photometer signals were recorded over a time of 60 s to detect a minimum of 1500 individual molecules. The measurement was performed at a final protein concentration of 100 nM. Subsequently, the raw data was processed using the DiscoverMP software.

### Model building for NIT1

The structure prediction of NIT1 monomer was generated by AlphaFold, and the structure prediction for NIT1 dimer was generated by AlphaFold2 Multimer ^123,124^ as implemented by ColabFold ^125^. Structural analysis and figure preparation were performed using UCSF ChimeraX ^126,127^.

### NIT and IMPDH protein sequence alignment and AIM prediction

For building sequence alignment of NIT and IMPDH protein orthologues and paralogues, AtNIT1 (AT3G44310) and AtIMPDH1 (AT1G79470) were used as bait, respectively. For extracting plant orthologous sequences, both Phytozome13 (https://phytozome-next.jgi.doe.gov/) and UniProt (https://www.uniprot.org/) were queried using Blast and the search was restricted to targeted species. For extracting animal and bacterial sequences, UniProt was used. After introduction of a species, protein alignments were built using Clustal Omega (1.2.4) ^128^ from multifasta files and multiple sequence alignments were viewed using Jalview with custom color settings. Eventual duplicated and truncated sequences were manually removed, and the same principle was reiterated several times to search additional species. AIM in NIT1 and IMPDH1 were searched using the iLIR (v1.0) prediction server ^129^. Trimmed alignments of the AIM-containing regions were extracted as images and AIMs were annotated by hand.

## Data availability

All raw data, tables and R scripts associated with the figures and for analysis have been uploaded to Zenodo. The mass spectrometry proteomics data have been deposited to the ProteomeXchange Consortium via the PRIDE ^130^ partner repository. Deep sequencing files have been deposited to the NCBI Gene expression Omnibus.

## Acknowledgements

We thank Vienna Biocenter Core Facilities (VBCF), particularly Proteomics, BioOptics and Plant Sciences. We thank the CLIP cluster (https://clip.science) for access for image analysis. We acknowledge funding from Austrian Academy of Sciences, Austrian Science Fund (FWF, P32355, P34944, I 6760, SFB F79, DOC 111), Vienna Science and Technology Fund (WWTF, LS17-047, LS21-009), and European Research Council Grant (Project number: 101043370). Marion Clavel, Juan Carlos de la Concepcion and Peng Gao are supported by the Vienna International Postdoctoral Program (VIP2) and Marie Curie Fellowship (Project number: 847548). We thank Alberto Moreno Cencerrado and Pawel Pasierbek for excellent assistance with confocal microscopy. We thank Christophe Ritzenthaler, Rick Vierstra, Paul Schulze-Lefert, Martinj Van Zanten, Andrea Weber, Masanori Izumi, Johannes Stuttman, Farid El Kasmi for providing *Arabidopsis thaliana* and *Nicotiana benthamiana* lines. We thank Fred Berger and Pierre Bourguet for providing the modified YFP seed coat cassette. We are grateful to Marco Incarbone for providing the TuMV-6k2-Scarlet infectious clone, and to Damien Garcia for the TCV infectious clone. We thank all members of the Dagdas lab for fruitful exchanges, critical reading and editing of the manuscript. We thank Caroline Gutjahr and Claudia Köhler for critical reading of the manuscript.

The authors declare no competing financial interest.

## Author contributions

M. Clavel performed molecular cloning, plant treatments and inoculation, confocal microscopy and subsequent analysis, western based experiments and quantification, BN-PAGE, RNA-based assays, GST-pulldown, affinity purification and samples preparation for MS, generated Arabidopsis lines, analysed protein enrichement in MS/MS, PRM and TMT experiments, generated all plots with the assistance of chatGPT 3.5, made figures, wrote the draft and manuscript revisions. A. Bianchi designed and generated *atg5* crispr lines, performed western based assays and inoculations and provided excellent feedback on the manuscript. R. Kobylinska performed protein extraction, western blots and inoculations. R. Groh performed protein purifications, GST pulldowns, MST and SEC-MALS measurments. J. Ma performed root electron microscopy and dual axis tomography. R. K. Papareddy performed trimming, alignment and differential gene expression of NGS dataset. N. Grujic performed plant selection. L. Picchianti performed protein purifications, GST pulldowns and mass photometry. E. Stewart performed design and preparation of phenotyping experiments and all analysis of image-based phenotyping data. M. Schutzbier and K. Stejskal performed sample preparation for proteomics and data collection. JC. de la Concepcion performed protein purifications and contributed critically to the manuscript. V. Sanchez de Medina Hernandez generated NIT1 dimer model and analysis of dimerization interface and provided excellent feedback on manuscript. Y. Voichek prepared libraries and performed initial analysis of NGS datasets. P. Clauw and J. Gunis extracted RNA for RNAseq. G. Durnberger performed k-means clustering. JC. Muelders performed GFP released assays and EDS1 coIPs. A. Grimm performed western blot assays. A. Sedivy performed MST and SEC-MALS measurments. M. Erhardt performed EM of infected leaf tissue. V. Vyboishikov developed oligomycin treatment for Arabidopsis and performed westerns based experiments. P. Gao performed TEM experiments. E. Lechner and E. Vantard performed plant growth and inoculation. J. Jez supervised all phenotyping experiments. E. Roitinger supervised all MS-related measurments. P. Genschik and BH. Kang supervised students. Y. Dagdas supervised students and postdocs, secured funding, wrote the draft and manuscript revisions.

