## Supplemental figures 1 to 14 for "Metabolic enzymes moonlight as selective autophagy receptors to protect plants against viral-induced cellular damage"

### FIGURE S1

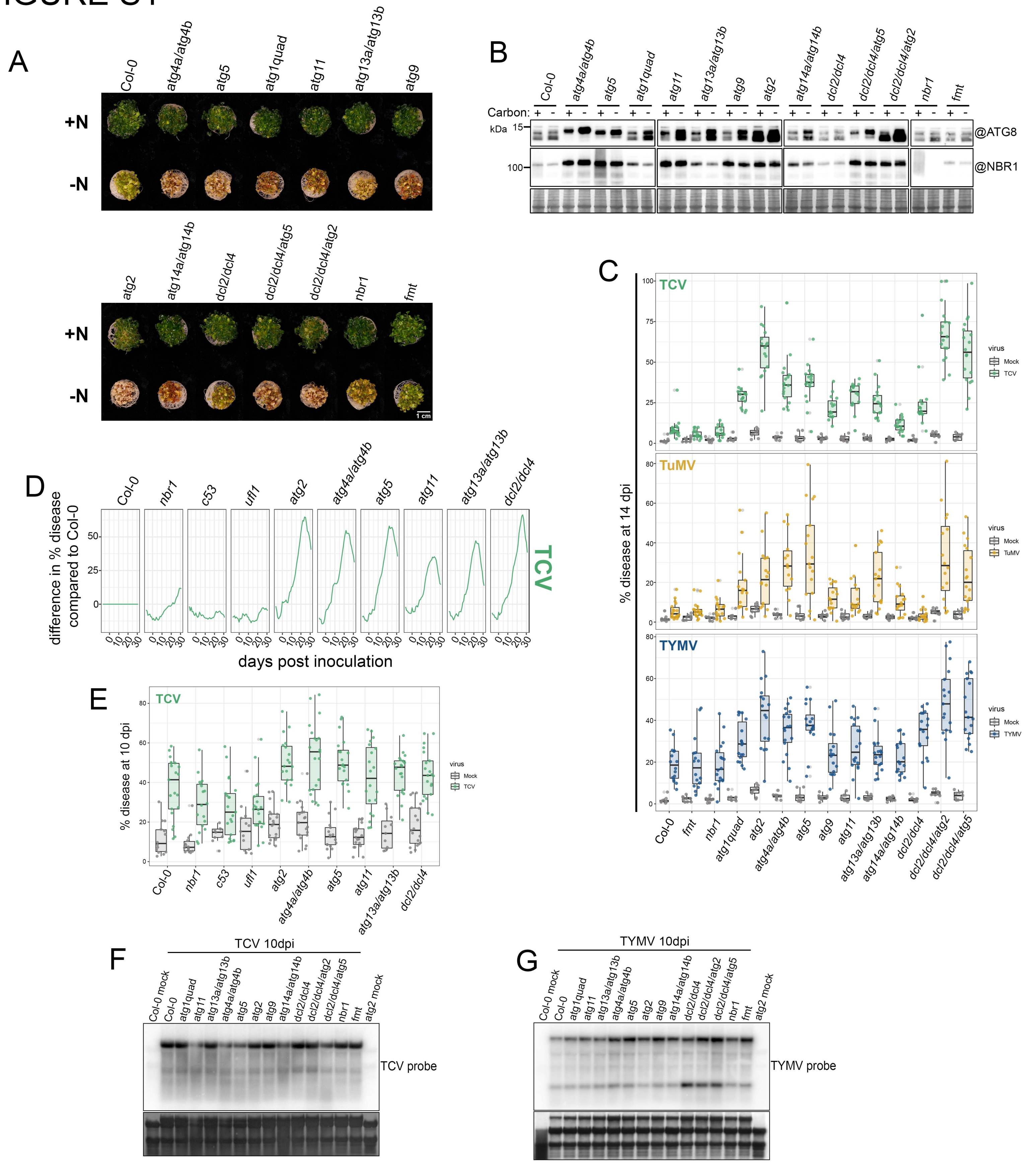
A
Col-0
atg4a/atg4b
atg5
atg1quad
atg11
atg13a/atg13b
atg9
+N
-N
atg2
atg14a/atg14b
dcl2/dcl4
dcl2/dcl4/atg5
dcl2/dcl4/atg2
nbr1
fmt
1 cm
B
Carbon:
kDa
15
100
@ATG8
@NBR1
C
TCV
TuMV
TYMV
% disease at 14 dpi
virus
Mock
D
difference in % disease compared to Col-0
days post inoculation
c53
ufl1
E
% disease at 10 dpi
F
TCV 10dpi
Col-0 mock
atg2 mock
TCV probe
G
TYMV 10dpi
TYMV probe

**FIGURE S1: Autophagy safeguards host fitness during viral infection but does not degrade viral components (supports Figure 1). (A)** Phenotypic characterization of Col-0, *atg4* double mutant, *atg5*, *atg1* quadruple mutant, *atg11, atg13* double mutant*, atg9, atg2, atg14* double mutant, *dcl2/dcl4* double mutant, *dcl2/dcl4/atg2* triple mutant*, dcl2/dcl4/atg5* triple mutant, *nbr1* and *fmt* upon nitrogen starvation. Seeds were sown in liquid ½ MS and grown for nine days. Media was either replenished (+N) or exchanged for nitrogen-deficient ½ MS media (-N) and grown for another six days before pictures were taken. **(B)** ATG8 lipidation assay and NBR1 abundance in the same mutants as in (C) in carbon replete (+) or during carbon starvation (-). **(C)** Quantification of % of disease in TCV, TuMV-6k2-sc and TYMV-infected systemic leaves for the indicated genotypes at 14 dpi. % disease is defined as plant area harboring symptomatic colors, and each data point is a single individual. **(D)** Phenotypic characterization of *Arabidopsis* Col-0 (WT), *nbr1, c53, ufl1, atg2, atg4* double mutant*, atg5, atg11, atg13* double mutant*, dcl2/dcl4* double mutant using automated camera phenotyping. % of disease is defined as plant area harboring symptomatic colors. The obtained values for Col-0 is subtracted to all mutants to obtain the difference in % disease compared to Col-0 from -10 days (pre inoculation) to + 30 days (post inoculation). **(E)** Quantification of % of disease in the TCV-infected systemic leaves for the indicated genotypes at 10 dpi. % disease is defined as in (C) and each data point is a single individual. **(F)** Northern blot of viral (+) RNA accumulation in TCV-infected systemic leaves of the indicated genotypes. 3-week-old rosettes were inoculated with buffer (mock) or with TCV and young systemic leaves were collected at 10 dpi. n=3 to 4 plants per sample-genotype combination. 5µg of total RNA was loaded per well and membrane was hybridized with a radiolabeled probe specific for the 3' sequence of TCV. Membrane was stained with methylene blue to verify loading. **(G)** Northern blot of viral (+) RNA accumulation in TYMV-infected systemic leaves of the indicated genotypes. 3-week-old rosettes were inoculated with buffer (mock) or with TYMV and young systemic leaves were collected at 10 dpi. n=3 to 4 plants per sample-genotype combination. 5µg of total RNA was loaded per well and membrane was hybridized with a radiolabeled probe specific for the 3' sequence of TYMV. Membrane was stained with methylene blue to verify loading.

### FIGURE S2

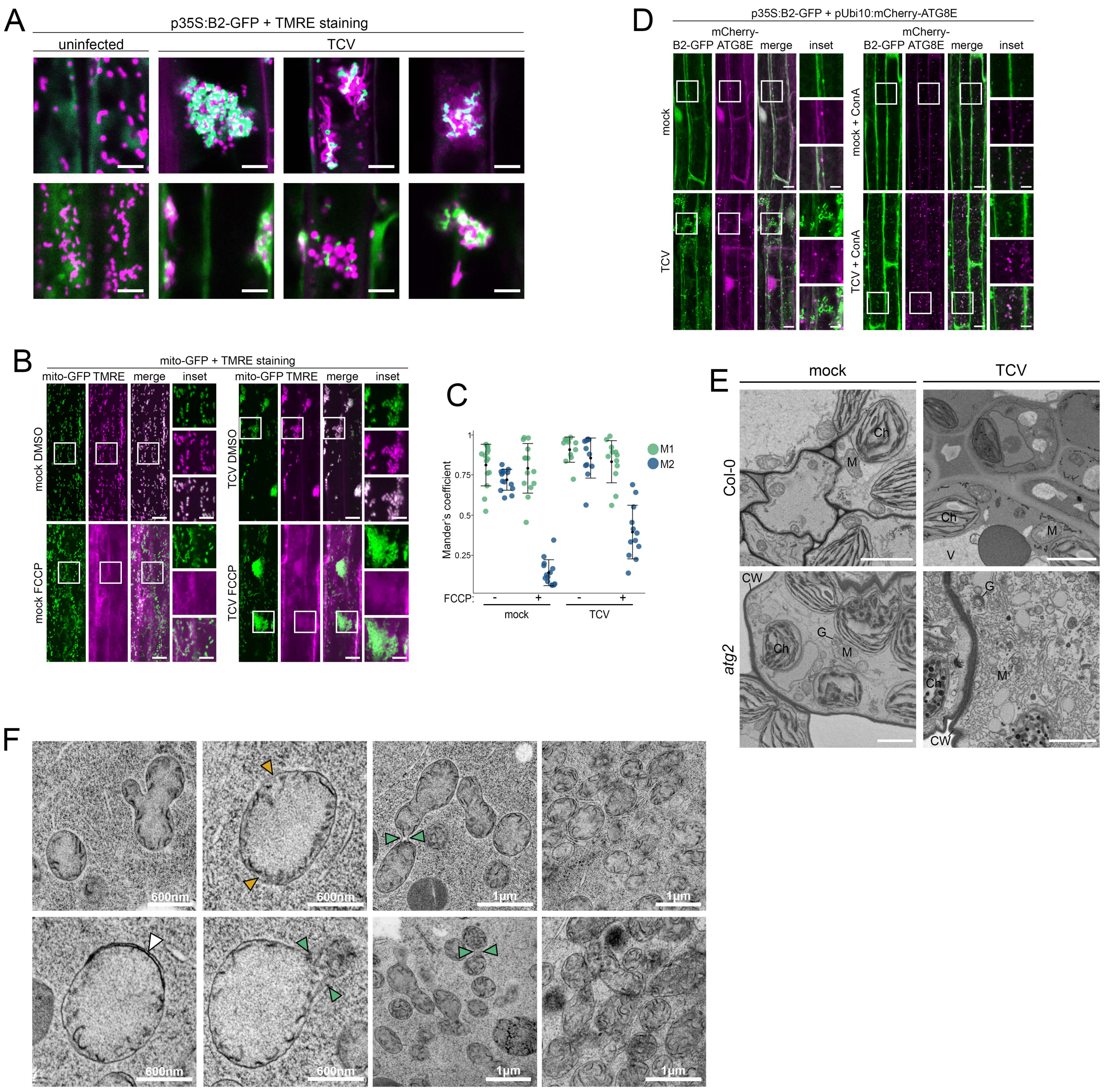
A
p35S:B2-GFP + TMRE staining
uninfected
TCV
D
p35S:B2-GFP + pUbi10:mCherry-ATG8E
B2-GFP
mCherry-ATG8E
merge
inset
mock
TCV
mock + ConA
TCV + ConA
B
mito-GFP + TMRE staining
mito-GFP
TMRE
merge
inset
mock DMSO
mock FCCP
TCV DMSO
TCV FCCP
C
Mander's coefficient
M1
M2
FCCP:
-
+
mock
TCV
E
mock
TCV
Col-0
atg2
Ch
M
V
CW
G
F
600nm
1µm

**FIGURE S2: Viral replication organelle remodeling induces autophagy (supports Figure 2). (A)** Gallery of Viral replication complexes in roots of *Arabidopsis* stable lines expressing B2-GFP (green) under the 35S promoter and stained with TMRE (magenta) in mock-inoculated or TCV-infected conditions at 5dpi. Seedlings were immerged in ½ MS with 500nM of TMRE for a few minutes then mounted in ½ MS for image acquisition. Each image is of a single slice obtained by CLSM. Scale bar 5µm. **(B)** Spinning disk microscopy images of *Arabidopsis* roots expressing mito-GFP (green) and stained with TMRE (magenta). 9-day-old seedlings were inoculated with TCV and systemic root images were acquired at 6dpi. Whole seedlings were additionally treated with 150µM of FCCP or an equivalent volume of DMSO in ½ MS medium for 10 to 60 minutes before being submerged in ½ MS with 500nM of TMRE for a few minutes then mounted in ½ MS for image acquisition. Each image is a maximum intensity projection of 10 to 20 slices. Scale bar 10µm and 5µm in inset. **(C)** Quantification of dataset shown in (B). Mander's colocalization coefficient between mitoGFP and TMRE signal, obtained from two single slices per captured z-stack. M1, fraction of mitoGFP overlapping with TMRE. M2, fraction of TMRE overlapping with mitoGFP. Bars indicate the mean ± SD of 12-14 slices. **(D)** Confocal microscopy images of *Arabidopsis* root expressing B2-GFP (green) under the 35S promoter and mCherry-ATG8E (magenta) under a ubiquitin 10 promoter in mock-inoculated or TCV-infected conditions at 5 and 6 dpi. Left panel: Seedlings were directly mounted in ½ MS for image acquisition. Each image is a maximum intensity projection of a full z-stack. Right panel: Seedlings were immerged in ½ MS with 1µM ConcanamycinA (ConA) for 3.5h to 5h then directly mounted in ½ MS for image acquisition. Each image is a single snap in the focal plane of the vacuole. Scale bar 10µm and 5µm in inset. See Figure 2 (H) and (I) for quantification. **(E)** Representative electron micrographs of mock and TCV infected Col-0 and *atg2* systemic leaf cells at 7 dpi. Electron dense tubular structures accumulates in *atg2* infected cells. M: mitochondria, Ch: chloroplast, G: Golgi stack, CW: cell wall, V: vacuole. Scale bar 5µm. **(F)** Representative root cells electron micrographs of 4-day-old seedlings expressing P29-mTurquoise2-myc. Upon expression of the viral protein P29, mitochondria show enlarged and abnormal morphology with marked constrictions (green arrows) membrane proliferation (white arrows) and loss of membrane integrity (yellow arrows) as well as clumping. Scale is as indicated in the image.

### FIGURE S3

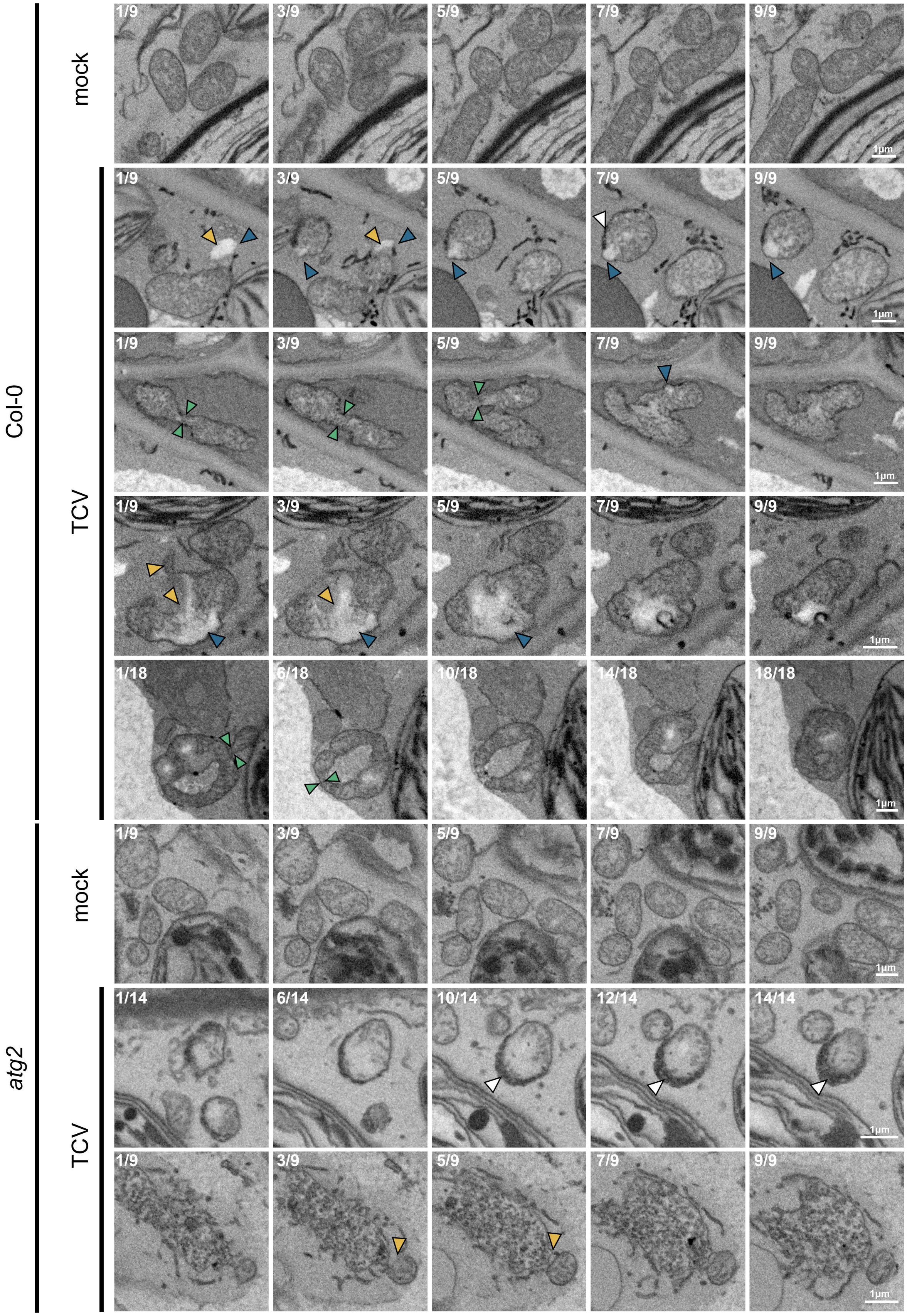

Col-0
mock
TCV
atg2
mock
TCV
1/9
3/9
5/9
7/9
9/9
1/18
6/18
10/18
14/18
18/18
1/14
6/14
10/14
12/14
14/14
1μm

**FIGURE S3: TCV replication-associated damage to mitochondria (supports Figure 2).** Serial sections of transmission electron microscopy micrographs showing diverse mitochondria shapes in mock and TCV infected Col-0 and *atg2* systemic leaf cells at 7 dpi. Mitochondria in infected cells display a variety of defects such as mitochondria herniation (blue arrows), complete loss of membrane integrity (yellow arrows), constrictions (green arrows), and apparent membrane thickening (white arrows). Note that one panel of Figure 2J is extracted from the last serial in *atg2*. Scale bar 1μm.

### FIGURE S4

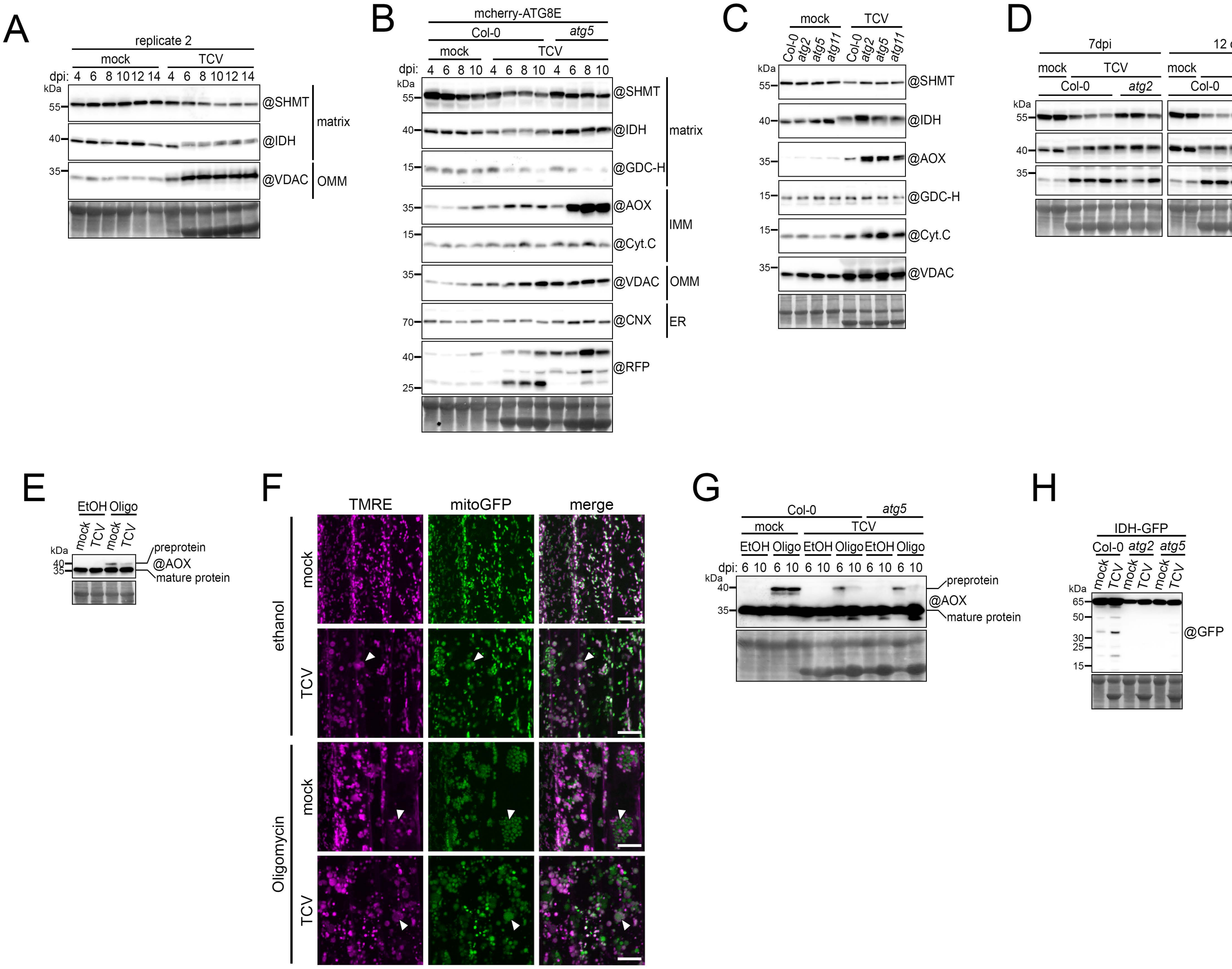
A
replicate 2
mock
TCV
dpi: 4 6 8 10 12 14 4 6 8 10 12 14
kDa
55
40
35
@SHMT
@IDH
@VDAC
matrix
OMM
B
mcherry-ATG8E
Col-0
atg5
mock
TCV
dpi: 4 6 8 10 4 6 8 10 4 6 8 10
@SHMT
@IDH
@GDC-H
@AOX
@Cyt.C
@VDAC
@CNX
@RFP
matrix
IMM
OMM
ER
C
mock
TCV
Col-0 atg2 atg5 atg11 Col-0 atg2 atg5 atg11
@SHMT
@IDH
@AOX
@GDC-H
@Cyt.C
@VDAC
D
7dpi
12 dpi
mock
TCV
Col-0
atg2
@SHMT
@IDH
@VDAC
E
EtOH
Oligo
mock TCV mock TCV
preprotein
@AOX
mature protein
F
TMRE
mitoGFP
merge
ethanol
Oligomycin
mock
TCV
G
Col-0
atg5
mock
TCV
EtOH Oligo EtOH Oligo EtOH Oligo
dpi: 6 10 6 10 6 10 6 10 6 10 6 10
preprotein
@AOX
mature protein
H
IDH-GFP
Col-0 atg2 atg5
mock TCV mock TCV mock TCV
@GFP
HA-GFP-TOM5
Col-0 atg5
mock TCV mock TCV
@HA

**FIGURE S4: Autophagy degrades single mitochondrial proteins that leak into the cytosol (supports Figure 3).** **(A)** Immunoblot of mitochondrial proteins in mock inoculated and TCV infected systemic leaves. 3-week-old rosettes were inoculated with buffer (mock) or with TCV and young systemic leaves were collected at the indicated time points. n=3 to 4 plants per sample-timepoint combination. Total soluble proteins were extracted at a fixed fresh weight to buffer volume ratio and equal volume was loaded per well. Membranes were hybridized with the indicated antibodies (@). Membranes were then stained with amidoblack to verify loading. Also see Figure 3A. **(B)** Immunoblot of mitochondrial proteins in mock inoculated and TCV infected systemic leaves expressing mCherry-ATG8E in WT or *atg5* background. 3-week-old rosettes were inoculated with buffer (mock) or with TCV and young systemic leaves were collected at the indicated time points. n=3 to 4 plants per sample-timepoint combination. Total soluble proteins were extracted at a fixed fresh weight to buffer volume ratio and equal volume was loaded per well. Membranes were hybridized with the indicated antibodies (@). Membranes were then stained with amidoblack to verify loading. **(C)** Immunoblot of mitochondrial proteins in mock and TCV inoculated systemic leaves of WT, *atg2*, *atg5* and *atg11* plants. 3-week-old rosettes were inoculated with buffer (mock) or with TCV and young systemic leaves were collected at 12 dpi. n=10 plants. Total soluble proteins were extracted at a fixed fresh weight to buffer volume ratio and equal volume was loaded per well. Membranes were hybridized with the indicated antibodies (@). Membranes were then stained with amidoblack to verify loading. Also see Figure 3D. (**D)** Immunoblot of mitochondrial proteins in mock and TCV inoculated systemic leaves of WT and *atg2* plants. 3-week-old rosettes were inoculated with buffer (mock) or with TCV and young systemic leaves were collected at the indicated time points. n=10 plants per sample-timepoint combination, each lane is a biological replicate. Total soluble proteins were extracted at a fixed fresh weight to buffer volume ratio and equal volume was loaded per well. Membranes were hybridized with the indicated antibodies (@). Membranes were then stained with amidoblack to verify loading. See Figure 3E for quantification. **(E)** Western blots showing mature and pre-protein of endogenous AOX1/2 protein from *Arabidopsis* whole seedling. 9-day-old seedlings were mock-inoculated or TCV-infected and grown for an additional 5 days before being transferred to liquid ½ MS + 1% sucrose with either ethanol (EtOH, vehicle) or 20μM Oligomycin A (Oligo) for 8.5 hours. 10μg of total protein was loaded per well and membrane was hybridized with an antibody against AOX1/2 protein. Membrane was stained with amidoblack to verify loading. Also see Figure 3G. **(F)** Confocal microscopy images of *Arabidopsis* roots expressing mito-GFP (green) and stained with TMRE (magenta). 9-day-old seedlings were inoculated with TCV and systemic root images were acquired at 5dpi. Whole seedlings were additionally treated with 20μM of Oligomycin A or an equivalent volume of ethanol in ½ MS medium for 5.5 to 7.5 hours before being submerged in ½ MS with 500nM of TMRE for a few minutes then mounted in ½ MS for image acquisition. White arrows show swollen and/or depolarized mitochondria that can occur in some TCV infected cells, are caused by Oligomycin A treatment and exacerbated by the double treatment. Each image is a maximum intensity projection of 6 to 8 slices 2μm apart. Scale bar 10μm. **(G)** Western blots showing mature and pre-protein of endogenous AOX1/2 protein from Col-0 and *atg5* whole seedling. 9-day-old seedlings were mock-inoculated or TCV-infected and grown for an additional 6 and 10 days before being transferred to liquid ½ MS + 1% sucrose with either ethanol (EtOH, vehicle) or 20 μM Oligomycin A (Oligo) for 10 hours. 25 μg of total protein was loaded per well and membrane was hybridized with an antibody against AOX1/2 protein. Membrane was stained with amidoblack to verify loading. Also see Figure 3G. **(H)** Western blots showing IDH-GFP and HA-GFP-TOM5 cleavage level in mock and TCV inoculated systemic leaves of Col-0, *atg5* and/or *atg2*. 3-week-old rosettes were inoculated with buffer (mock) or with TCV and young systemic leaves were collected at the indicated time points. n=10 plants per condition. Total proteins were extracted and 25μg of total protein was loaded per well and membranes were hybridized with the indicated antibodies (@). Membranes were then stained with amidoblack to verify loading. Also see Figure 3L.

FIGURE S5

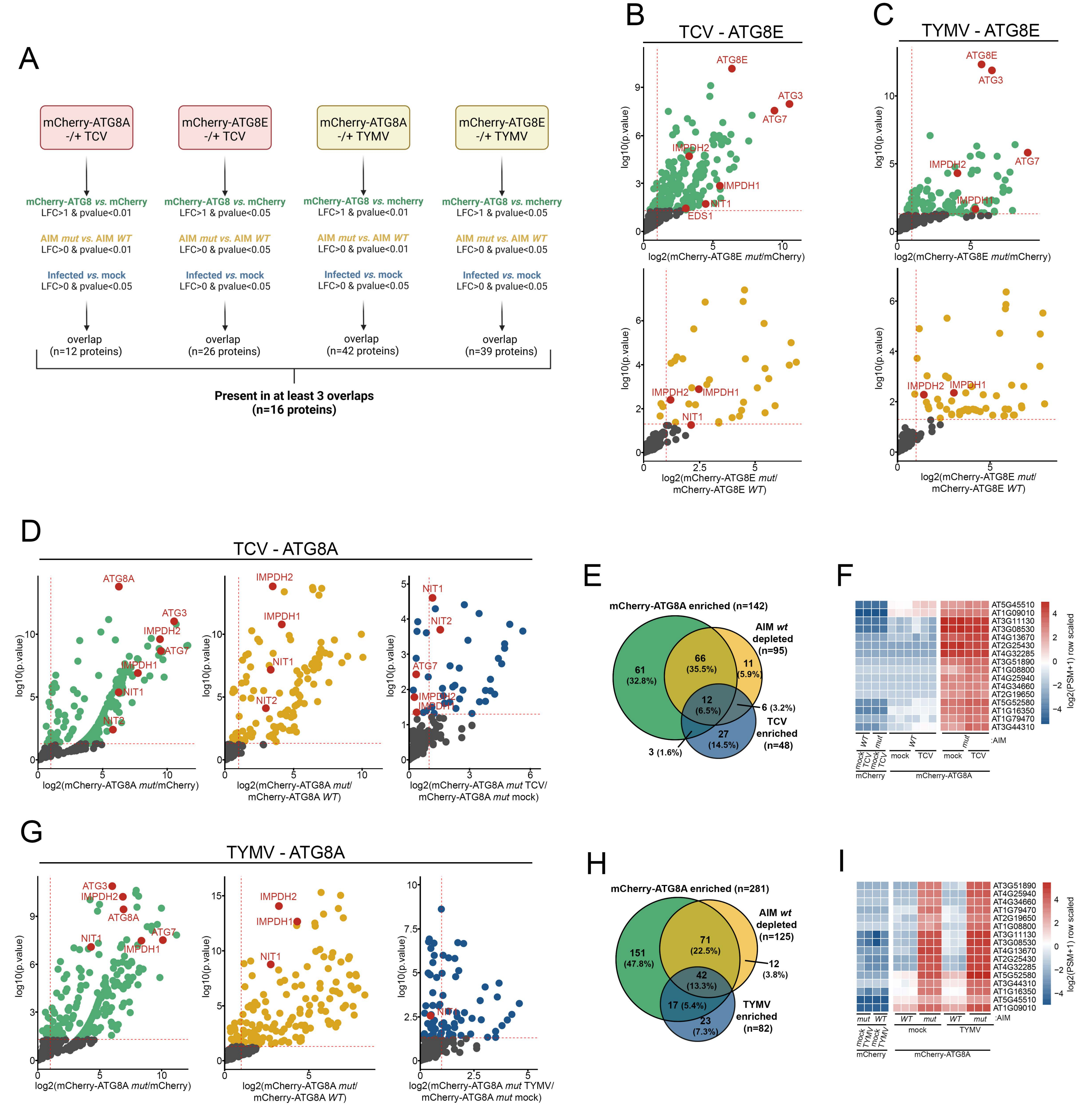
A
mCherry-ATG8A -/+ TCV
mCherry-ATG8E -/+ TCV
mCherry-ATG8A -/+ TYMV
mCherry-ATG8E -/+ TYMV
mCherry-ATG8 vs. mCherry
LFC>1 & pvalue<0.01
mCherry-ATG8 vs. mCherry
LFC>1 & pvalue<0.05
mCherry-ATG8 vs. mcherry
LFC>1 & pvalue<0.01
mCherry-ATG8 vs. mcherry
LFC>1 & pvalue<0.05
AIM mut vs. AIM WT
LFC>0 & pvalue<0.01
AIM mut vs. AIM WT
LFC>0 & pvalue<0.05
AIM mut vs. AIM WT
LFC>0 & pvalue<0.01
AIM mut vs. AIM WT
LFC>0 & pvalue<0.05
Infected vs. mock
LFC>0 & pvalue<0.05
overlap
(n=12 proteins)
overlap
(n=26 proteins)
overlap
(n=42 proteins)
overlap
(n=39 proteins)
Present in at least 3 overlaps
(n=16 proteins)
B
TCV - ATG8E
C
TYMV - ATG8E
log10(p.value)
log2(mCherry-ATG8E mut/mCherry)
log2(mCherry-ATG8E mut/
mCherry-ATG8E WT)
ATG8E
ATG3
ATG7
IMPDH2
IMPDH1
NIT1
EDS1
D
TCV - ATG8A
log2(mCherry-ATG8A mut/mCherry)
log2(mCherry-ATG8A mut/
mCherry-ATG8A WT)
log2(mCherry-ATG8A mut TCV/
mCherry-ATG8A mut mock)
ATG8A
NIT2
E
mCherry-ATG8A enriched (n=142)
AIM wt
depleted
(n=95)
TCV
enriched
(n=48)
61
(32.8%)
66
(35.5%)
11
(5.9%)
12
(6.5%)
6 (3.2%)
27
(14.5%)
3 (1.6%)
F
AT5G45510
AT1G09010
AT3G11130
AT3G08530
AT4G13670
AT2G25430
AT4G32285
AT3G51890
AT1G08800
AT4G25940
AT4G34660
AT2G19650
AT5G52580
AT1G16350
AT1G79470
AT3G44310
log2(PSM+1) row scaled
:AIM
mock
TCV
mCherry
mCherry-ATG8A
G
TYMV - ATG8A
log2(mCherry-ATG8A mut TYMV/
mCherry-ATG8A mut mock)
H
mCherry-ATG8A enriched (n=281)
AIM wt
depleted
(n=125)
TYMV
enriched
(n=82)
151
(47.8%)
71
(22.5%)
12
(3.8%)
42
(13.3%)
17 (5.4%)
23
(7.3%)
I
AT3G51890
AT4G25940
AT4G34660
AT1G79470
AT2G19650
AT1G08800
AT3G11130
AT3G08530
AT4G13670
AT2G25430
AT4G32285
AT5G52580
AT3G44310
AT1G16350
AT5G45510
AT1G09010

**FIGURE S5: Identification of NIT1 and IMPDH1 as infection specific SARs (supports Figure 4). (A)** Selection scheme employed for selecting the final list of candidate Virus-specific Selective Autophagy Receptors (VSAR). Proteins were considered as final candidates of they were present at the overlap between the three pairwise comparisons for at least three out of four datasets. **(B)** Top panel: enrichment of proteins co-purified with mCherry-ATG8E in TCV-infected and mock systemic leaves *vs.* mCherry control line in the identical condition, represented by a volcano plot (green data points). Bottom panel: enrichment of proteins co-purified with mCherry-ATG8E in AIM mutant-treated samples (mock and TCV combined) *vs.* mCherry-ATG8E in AIM *wt*-treated samples (mock and TCV combined), represented by a volcano plot (yellow data points). The horizontal dashed line indicates the threshold above which proteins are significantly enriched ($p$ value < 0.05, quasi-likelihood negative binomial generalized log-linear model) and the vertical dashed line the threshold for which proteins log2 fold change is above 1. **(C)** Top panel: enrichment of proteins co-purified with mCherry-ATG8E in TYMV-infected and mock systemic leaves *vs.* mCherry control line in the identical condition, represented by a volcano plot (green data points). Bottom panel: enrichment of proteins co-purified with mCherry-ATG8E in AIM mutant-treated samples (mock and TYMV combined) *vs.* mCherry-ATG8E in AIM *wt*-treated samples (mock and TYMV combined), represented by a volcano plot (yellow data points). The horizontal dashed line indicates the threshold above which proteins are significantly enriched ($p$ value < 0.05, quasi-likelihood negative binomial generalized log-linear model) and the vertical dashed line the threshold for which proteins log2 fold change is above 1. **(D)** Enrichment of proteins co-purified with mCherry-ATG8A in TCV-infected systemic leaves in the indicated pairwise comparisons. Colors and comparisons are the same as in panel A and B and Figure 4 A and D. **(E)** Venn diagram of three overlapping pairwise comparisons for AP-MS conducted in mCherry-ATG8A plants infected with TCV: mCherry-ATG8A (only mutant AIM-treated replicates) *vs.* mCherry control; green circle. mCherry-ATG8A AIM mutant-treated samples (mock and TCV treated combined) *vs.* mCherry-ATG8A AIM WT-treated samples; yellow circle. mCherry-ATG8A TCV-treated samples (mutant AIM-treated only) *vs.* mCherry-ATG8A mock-treated samples (mutant AIM-treated only); blue circle. **(F)** Protein abundance pattern represented by a heatmap (Log2(PSM+1) – meanPSM per protein) for the sixteen proteins identified as uniquely enriched in AIM-dependent ATG8 interactome upon infection (at least three of four datasets), in the mCherry-ATG8A TCV dataset. **(G)** Enrichment of proteins co-purified with mCherry-ATG8A in TYMV-infected systemic leaves in the indicated pairwise comparisons. Colors and comparisons are the same as in panel A and B and Figure 4 A and D. **(H)** Venn diagram of three overlapping pairwise comparisons for AP-MS conducted in mcherry-ATG8A plants infected with TYMV: mCherry-ATG8A (only mutant AIM-treated replicates) *vs.* mCherry control; green circle. mCherry-ATG8A AIM mutant-treated samples (mock and TYMV treated combined) *vs.* mCherry-ATG8A AIM *wt*-treated samples; yellow circle. mCherry-ATG8A TYMV-treated samples (mutant AIM-treated only) *vs.* mCherry-ATG8A mock-treated samples (mutant AIM-treated only); blue circle. **(I)** Protein abundance pattern represented by a heatmap (Log2(PSM+1) – meanPSM per protein) for the sixteen proteins identified as uniquely enriched in AIM-dependent ATG8 interactome upon infection (at least three of four datasets), in the mCherry-ATG8A TYMV dataset.

### FIGURE S6

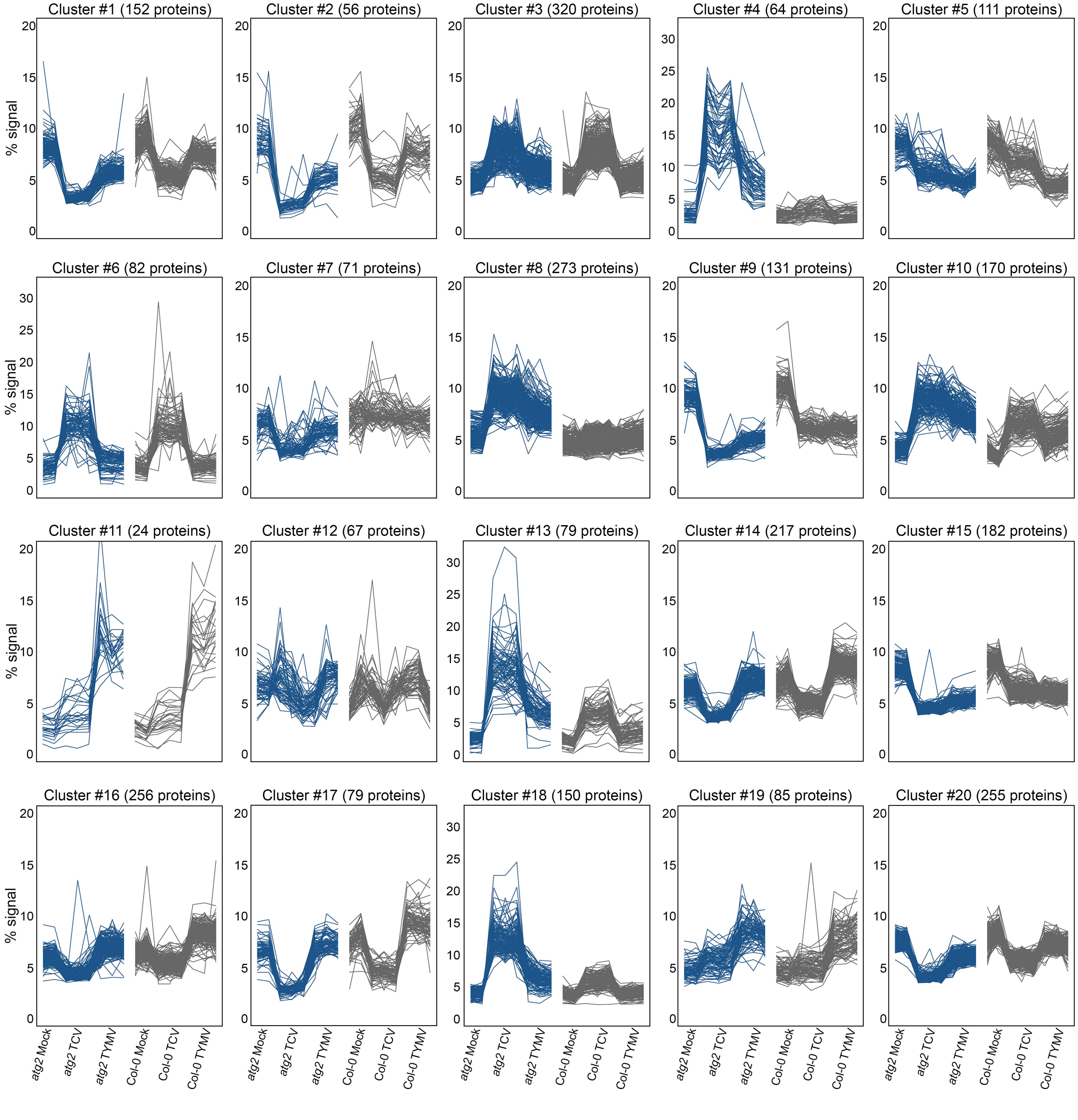
Cluster #1 (152 proteins)
Cluster #2 (56 proteins)
Cluster #3 (320 proteins)
Cluster #4 (64 proteins)
Cluster #5 (111 proteins)
Cluster #6 (82 proteins)
Cluster #7 (71 proteins)
Cluster #8 (273 proteins)
Cluster #9 (131 proteins)
Cluster #10 (170 proteins)
Cluster #11 (24 proteins)
Cluster #12 (67 proteins)
Cluster #13 (79 proteins)
Cluster #14 (217 proteins)
Cluster #15 (182 proteins)
Cluster #16 (256 proteins)
Cluster #17 (79 proteins)
Cluster #18 (150 proteins)
Cluster #19 (85 proteins)
Cluster #20 (255 proteins)
% signal
atg2 Mock
atg2 TCV
atg2 TYMV
Col-0 Mock
Col-0 TCV
Col-0 TYMV

**FIGURE S6: Quantitative proteomics of infected systemic tissue reveals autophagy-dependent degradome (supports Figure 4).** Protein abundance was measured in mock, TCV and TYMV infected systemic leaves of Col-0 and *atg2* plants (duplicates for mock and triplicates for infected samples) using 16-plex TMT labelling. Normalized abundance of proteins regulated > 2-fold was then used for k-means clustering and sorted in 20 separate clusters.

FIGURE S7

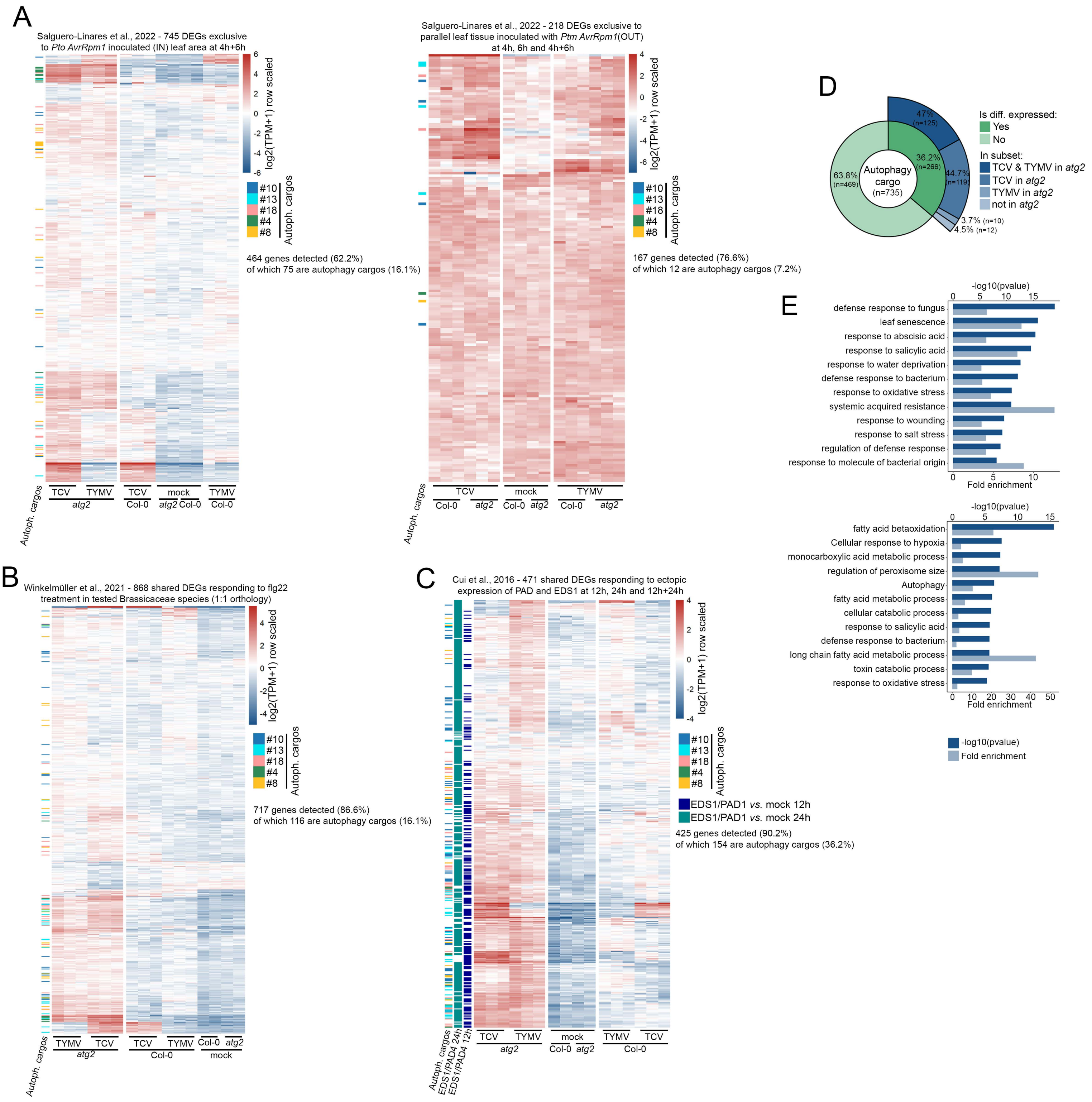

A
Salguero-Linares et al., 2022 - 745 DEGs exclusive to Pto AvrRpm1 inoculated (IN) leaf area at 4h+6h
log2(TPM+1) row scaled
#10
#13
#18
#4
#8
Autoph. cargos
464 genes detected (62.2%)
of which 75 are autophagy cargos (16.1%)
TCV
TYMV
atg2
TCV
Col-0
mock
atg2 Col-0
TYMV
Col-0
Salguero-Linares et al., 2022 - 218 DEGs exclusive to parallel leaf tissue inoculated with Ptm AvrRpm1(OUT) at 4h, 6h and 4h+6h
167 genes detected (76.6%)
of which 12 are autophagy cargos (7.2%)
TCV
Col-0
atg2
mock
Col-0
atg2
TYMV
Col-0
atg2
B
Winkelmüller et al., 2021 - 868 shared DEGs responding to flg22 treatment in tested Brassicaceae species (1:1 orthology)
717 genes detected (86.6%)
of which 116 are autophagy cargos (16.1%)
TYMV
TCV
atg2
TCV
TYMV
Col-0
Col-0
atg2
mock
C
Cui et al., 2016 - 471 shared DEGs responding to ectopic expression of PAD and EDS1 at 12h, 24h and 12h+24h
EDS1/PAD1 vs. mock 12h
EDS1/PAD1 vs. mock 24h
425 genes detected (90.2%)
of which 154 are autophagy cargos (36.2%)
Autoph. cargos
EDS1/PAD4 24h
EDS1/PAD4 12h
TCV
TYMV
atg2
mock
Col-0
atg2
TYMV
TCV
Col-0
D
Is diff. expressed:
Yes
No
In subset:
TCV & TYMV in atg2
TCV in atg2
TYMV in atg2
not in atg2
Autophagy cargo (n=735)
63.8% (n=469)
36.2% (n=266)
47% (n=125)
44.7% (n=119)
3.7% (n=10)
4.5% (n=12)
E
-log10(pvalue)
defense response to fungus
leaf senescence
response to abscisic acid
response to salicylic acid
response to water deprivation
defense response to bacterium
response to oxidative stress
systemic acquired resistance
response to wounding
response to salt stress
regulation of defense response
response to molecule of bacterial origin
Fold enrichment
fatty acid betaoxidation
Cellular response to hypoxia
monocarboxylic acid metabolic process
regulation of peroxisome size
Autophagy
fatty acid metabolic process
cellular catabolic process
response to salicylic acid
defense response to bacterium
long chain fatty acid metabolic process
toxin catabolic process
response to oxidative stress
-log10(pvalue)
Fold enrichment

**FIGURE S7: Transcriptional reprogramming in *atg2* infected plants bears hallmarks of immune response (supports Figure 4). (A)** Left panel: Expression of genes defined as induced by *Pto AvrRpm1* in the inoculated portion of Arabidopsis leaf (IN), at both 4h and 6h post inoculation. DEGs list was recovered from supplemental table 3 of Salguero-Linares et al., 2022, and their expression is plotted as Log2(TPM+1) and normalized to the mean value per gene, in the libraries generated in the present study. Right panel: Expression of upregulated genes defined as induced by *Pto AvrRpm1* in the region adjacent to the inoculated portion of Arabidopsis leaf (OUT), at 4h, 6h and both, post inoculation. Gene list retrieved from the same study and plotted using the same method. Rows and columns were clustered and resulting dendrograms are omitted from the figure. Genes defined as autophagy cargos in Figure 4G are annotated to the left side of the heatmap. **(B)** Expression of differentially expressed genes defined as induced by flg22 treatment in *Arabidopsis thaliana*, *Capsella rubella*, *Eutrema salsugineum* and *Cardamine hirsute* with a 1:1 orthology relationship to genes annotated in Col-0. DEGs list was recovered from supplemental dataset 2 of Winkelmüller et al., 2021, and their expression is plotted as Log2(TPM+1) and normalized to the mean value per gene, in the libraries generated in the present study. Rows and columns were clustered and resulting dendrograms are omitted from the figure. Genes defined as autophagy cargos in Figure 4G are annotated to the left side of the heatmap. **(C)** Expression of differentially expressed genes defined as induced by ectopic expression of HA-PAD4 (beta estradiol inducible) and EDS1-HA (35S) in the *eds1/pad4* background at 12h, 24h and both. DEGs list was recovered from table S2 of Cui et al., 2016, and their expression is plotted as Log2(TPM+1) and normalized to the mean value per gene, in the libraries generated in the present study. Rows and columns were clustered and resulting dendrograms are omitted from the figure. Genes defined as autophagy cargos in Figure 4G and expression at 12h or 24h post beta estradiol treatment *vs.* DMSO in the original study are annotated to the left side of the heatmap. **(D)** Nested donut chart showing the percentage of virus-specific cargos assigned as DEGs (green inner donut). The cargos also being DEGS are further subdivided on the basis of genotype˜treatment responsible for their deregulation (blue outer donut). This analysis reveals that the majority of the cargos also being DEGs are explained by an *atg2*-specific transcriptional effect during infection. **(E)** Top GO terms for molecular function enriched in virus-specific autophagy cargos. Top panel: Top twelve enriched GO terms for molecular function for the cargos that are also DEGs (36.2%, 266 proteins). Bottom panel: Top twelve enriched GO terms for molecular function for the cargos that are not found amongst the DEGs (63.8%, 469 proteins). Terms are ranked according to their significance as calculated by DavidGO and -log10(pvalue) is represented on the top axis (dark blue) while fold enrichment as calculated by DavidGO is represented on the bottom axis (light blue).

### FIGURE S8

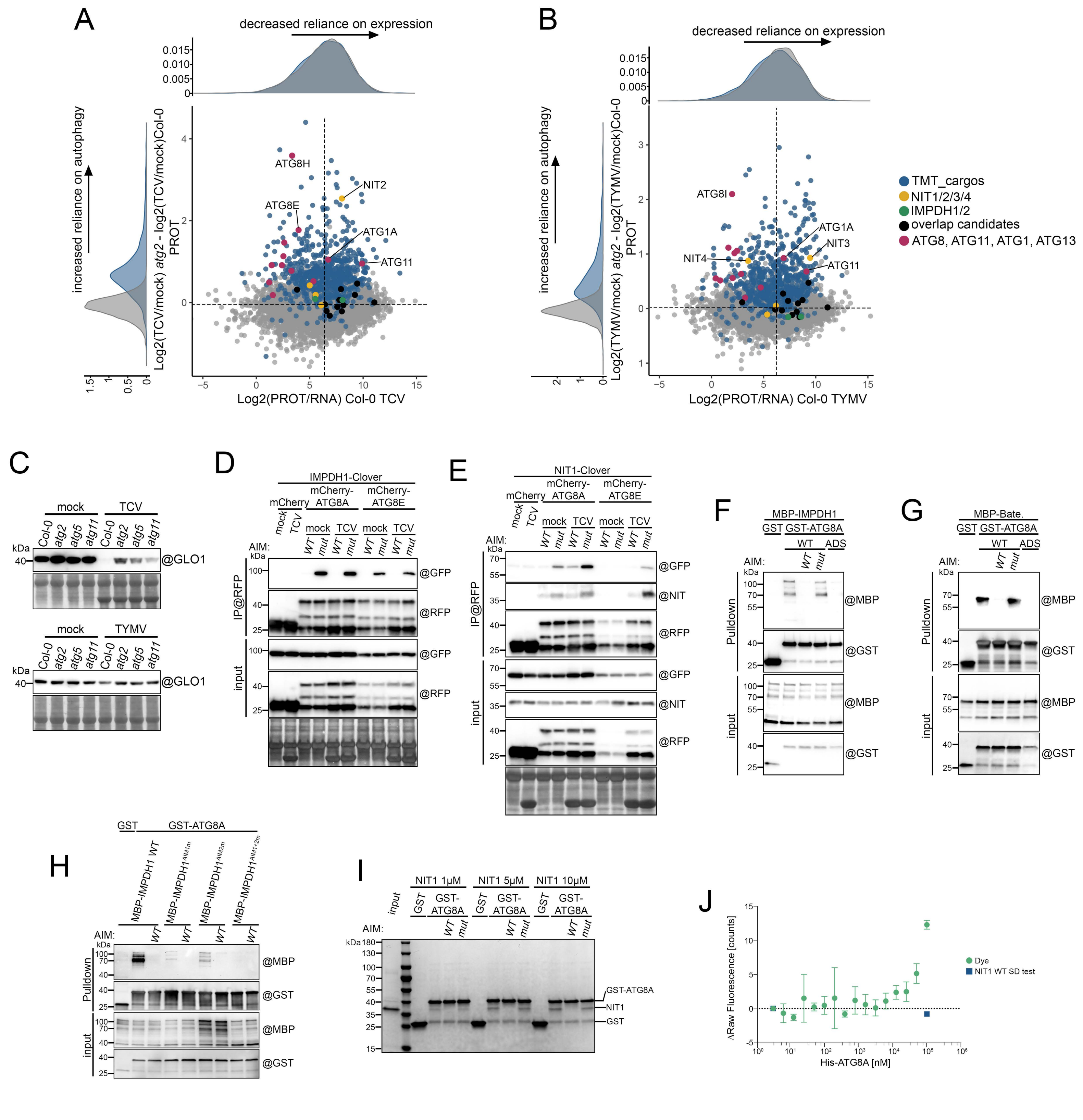

A
decreased reliance on expression
increased reliance on autophagy
Log2(TCV/mock) atg2 - log2(TCV/mock)Col-0 PROT
Log2(PROT/RNA) Col-0 TCV
ATG8H
NIT2
ATG8E
ATG1A
ATG11
B
decreased reliance on expression
increased reliance on autophagy
Log2(TYMV/mock) atg2 - log2(TYMV/mock)Col-0 PROT
Log2(PROT/RNA) Col-0 TYMV
ATG8I
ATG1A
NIT3
NIT4
ATG11
TMT_cargos
NIT1/2/3/4
IMPDH1/2
overlap candidates
ATG8, ATG11, ATG1, ATG13
C
mock
TCV
Col-0
atg2
atg5
atg11
@GLO1
TYMV
D
IMPDH1-Clover
mCherry
mCherry-ATG8A
mCherry-ATG8E
mock
TCV
AIM:
WT
mut
IP@RFP
input
@GFP
@RFP
E
NIT1-Clover
@NIT
F
MBP-IMPDH1
GST
GST-ATG8A
ADS
Pulldown
@MBP
@GST
G
MBP-Bate.
H
MBP-IMPDH1 WT
MBP-IMPDH1AIM1m
MBP-IMPDH1AIM2m
MBP-IMPDH1AIM1+2m
I
NIT1 1μM
NIT1 5μM
NIT1 10μM
GST-ATG8A
NIT1
GST
J
ΔRaw Fluorescence [counts]
His-ATG8A [nM]
Dye
NIT1 WT SD test

**FIGURE S8: NIT1 and IMPDH1 AIM-dependent interaction to ATG8 (supports Figure 4). (A)** Graphical representation of the ratio of protein abundance to RNA abundance in Col-0 TCV samples (log2(protein abundance/transcript abundance) as captured by quantitative proteomics and transcriptomic) on the x axis versus the ratio of protein abundance in atg2 TCV-infected samples with the removed contribution of the WT Col-0 background (log2(TCV/mock) in *atg2* – log2(TCV/mock) in Col-0) as captured by quantitative proteomics on the y axis. Reliance on transcription for overall abundance of a protein decreases as x axis values increase, while increased reliance on the autophagic machinery at the protein level increases with the y axis. Quadrants are determined using centroids for all the plotted datapoints. Data distribution for both axis is plotted next to the relevant axis. Proteins mapped as virus-specific autophagy cargos are colored in blue, known autophagy machinery components that undergo degradation are plotted in magenta and the detectable candidate SARs identified by AP-MS are plotted in black. NIT paralogues are plotted in yellow and IMPDH-paralogues in green (the later are not present in the cargo list). **(B)** Graphical representation of the ratio of protein abundance to RNA abundance in Col-0 TYMV samples (log2(protein abundance/transcript abundance) as captured by quantitative proteomics and transcriptomic) on the x axis versus the ratio of protein abundance in atg2 TYMV-infected samples with the removed contribution of the WT Col-0 background (log2(TYMV/mock) in *atg2* – log2(TYMV/mock) in Col-0) as captured by quantitative proteomics on the y axis. Reliance on transcription for overall abundance of a protein decreases as x axis values increase, while increased reliance on the autophagic machinery at the protein level increases with the y axis. Quadrants are determined using centroids for all the plotted datapoints. Data distribution for both axis is plotted next to the relevant axis. Proteins mapped as virus-specific autophagy cargos are colored in blue, known autophagy machinery components that undergo degradation are plotted in magenta and the detectable candidate SARs identified by AP-MS are plotted in black. NIT paralogues are plotted in yellow and IMPDH-paralogues in green (the later are not present in the cargo list). **(C)** Immunoblot of GLO1 protein in mock, TCV (upper panel) and TYMV (lower panel) inoculated systemic leaves of WT, *atg2*, *atg5* and *atg11* plants. 3-week-old rosettes were inoculated with buffer (mock) or with TCV and young systemic leaves were collected at 12 dpi. n=10 plants. Total soluble proteins were extracted at a fixed fresh weight to buffer volume ratio and equal volume was loaded per well. Membranes were hybridized with a GLO1 (GOX1) antibody (@). Membranes were then stained with amidoblack to verify loading. **(D)** *In vivo* affinity purification of mCherry-ATG8A and mCherry-ATG8E coupled to AIM peptide competition confirms AIM-dependent interaction with IMPDH1-Clover. Systemic leaves expressing mCherry control, mCherry-ATG8A or mCherry-ATG8E under ubiquitin 10 promoter and IMPDH1-Clover under HTR5 promoter in mock-inoculated or TCV-infected conditions at 11 dpi (n=40 plants). 150 µM of AIM peptide *wt* or mutant was added to the total cell lysate and affinity purification was performed using @RFP beads. Immunoblot against the indicated antibodies (@) was used to detect bait and prey in the input and RFP purified fraction. Membrane corresponding to the input fraction was then stained with amidoblack to verify loading. See Figure 4L for additional experimental evidence. **(E)** *In vivo* affinity purification of mCherry-ATG8A and mCherry-ATG8E coupled to AIM peptide competition confirms AIM-dependent interaction with NIT1-Clover and endogenous NIT protein. Systemic leaves expressing mCherry control, mCherry-ATG8A or mCherry-ATG8E under ubiquitin 10 promoter and IMPDH1-Clover under HTR5 promoter in mock-inoculated or TCV-infected conditions at 11 dpi (n=40 plants). 150 µM of AIM peptide *wt* or mutant was added to the total cell lysate and affinity purification was performed using @RFP beads. Immunoblot against the indicated antibodies (@) was used to detect bait and preys in the input and RFP purified fraction. Membrane corresponding to the input fraction was then stained with amidoblack to verify loading. See Figure 4M for additional experimental evidence. **(F)** IMPDH1 interacts with ATG8A in an AIM-dependent manner. Bacterial lysates containing recombinant MBP-IMPDH1, GST, GST-ATG8A WT or GST-ATG8A deficient in the AIM docking site (ADS) were mixed, 200 μM AIM *wt* or mutant peptide was added to the indicated reaction and proteins were pulled down using GST beads. Immunoblot against the indicated antibodies (@) was used to detect bait and prey in the input and pull-down fraction. See Figure 4N for additional experimental evidence. **(G)** The Bateman regulatory domain of IMPDH1 interacts with ATG8A in an AIM-dependent manner. Bacterial lysates containing recombinant MBP-Bateman, GST, GST-ATG8A WT or GST-ATG8A deficient in the AIM docking site (ADS) were mixed, 200 μM AIM *wt* or mutant peptide was added to the indicated reaction and proteins were pulled down using GST beads. Immunoblot against the indicated antibodies (@) was used to detect bait and prey in the input and pull-down fraction. See Figure 4O for additional experimental evidence. **(H)** The AIM1 in the Bateman domain of IMPDH1 mediates interaction with ATG8A. Bacterial lysates containing MBP-IMPDH1 WT, derivative AIM mutants, GST and GST-ATG8A were mixed with, 200 μM AIM peptide was added to the indicated reaction and proteins were pulled down using GST beads. Immunoblot against the indicated antibodies (@) was used to detect bait and prey in the input and pull-down fraction. See Figure 4P for additional experimental evidence. **(I)** NIT1 interacts with ATG8A in an AIM-dependent manner. Increasing concentrations of purified recombinant NIT1 were mixed with purified recombinant GST or GST-ATG8A, 200 μM AIM *wt* or mutant peptide was added to the indicated reaction and proteins were pulled down using GST beads. In-gel Coomassie staining was used to detect bait and prey proteins in the input and pull-down fraction. See Figure 4Q for additional experimental evidence. **(J)** Fluorescence binding control of ATG8A to dye shows no interaction. The ligand-induced Fluorescence change specificity was tested by the SDS denaturation test (SD-Test).

### FIGURE S9

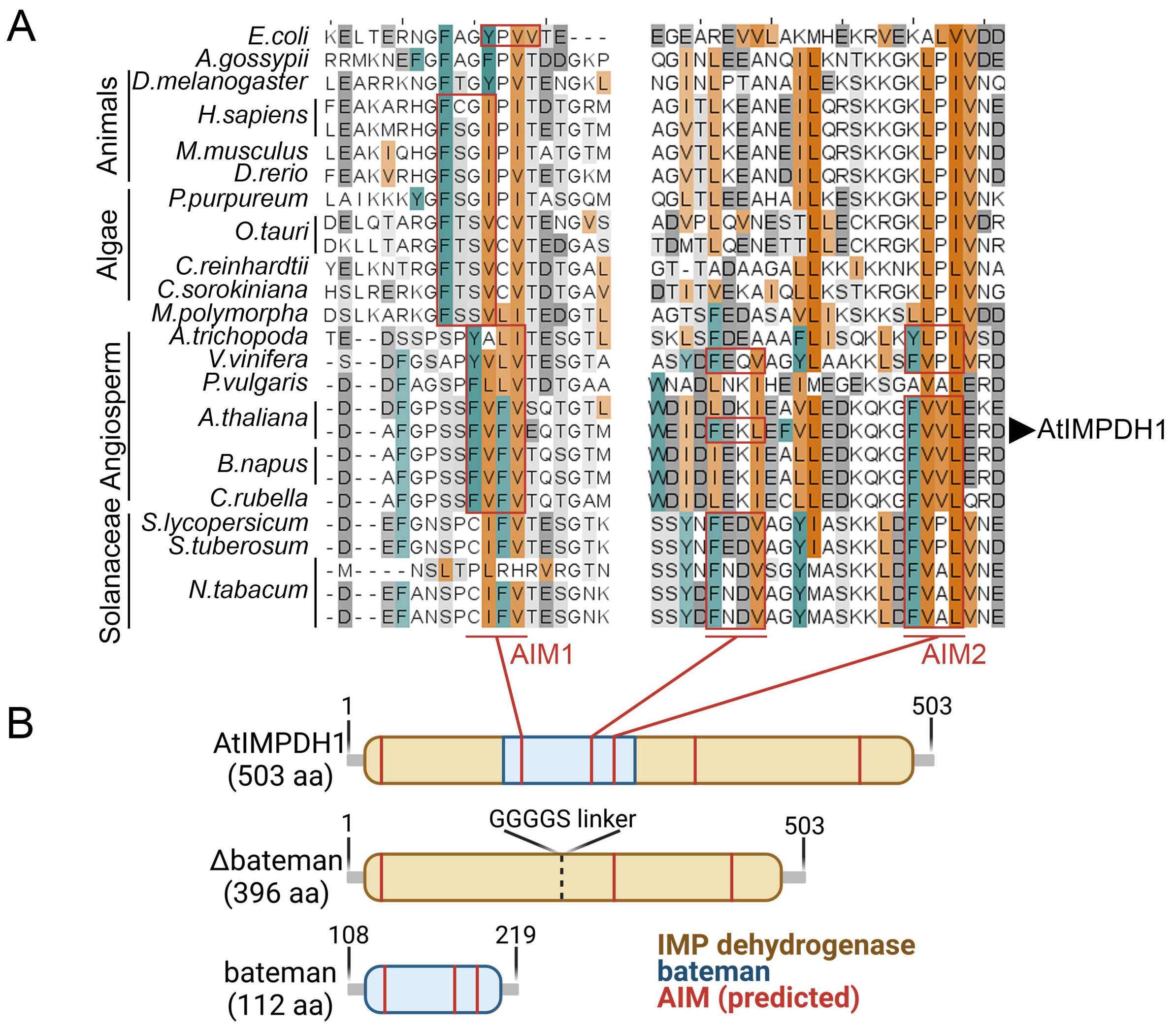
A
E.coli
A.gossypii
D.melanogaster
H.sapiens
M.musculus
D.rerio
P.purpureum
O.tauri
C.reinhardtii
C.sorokiniana
M.polymorpha
A.trichopoda
V.vinifera
P.vulgaris
A.thaliana
B.napus
C.rubella
S.lycopersicum
S.tuberosum
N.tabacum
Animals
Algae
Angiosperm
Solanaceae
AtIMPDH1
AIM1
AIM2
B
1
503
AtIMPDH1
(503 aa)
GGGGS linker
Δbateman
(396 aa)
108
219
bateman
(112 aa)
IMP dehydrogenase
bateman
AIM (predicted)

**FIGURE S9: AIM positions and conservation in IMPDH-type proteins (supports Figure 4). (A)** Trimmed multiple sequence alignment of IMPDH proteins depicting the conservation of the predicted canonical AIMs (iLIR on Arabidopsis IMPDH1 sequence used as bait) and their position in the aligned proteins. AIMs are boxed in red, W/F/Y (position Θ) residues are underlined in teal, L/I/V residues (position Γ) in orange and S/T (phosphorylatable) and E/D (negatively charged) are underlined in light and dark grey respectively. Color intensity is proportional to conservation (cutoff threshold 10%). Species of origin are indicated on the left. **(B)** Schematic representation of linear protein domain organization and putative AIM positions within the sequence of the indicated protein. Only AIMs that were experimentally tested are numbered. Schematic representation of the truncation constructs used for pulldown assays in Figure 4.

### FIGURE S10

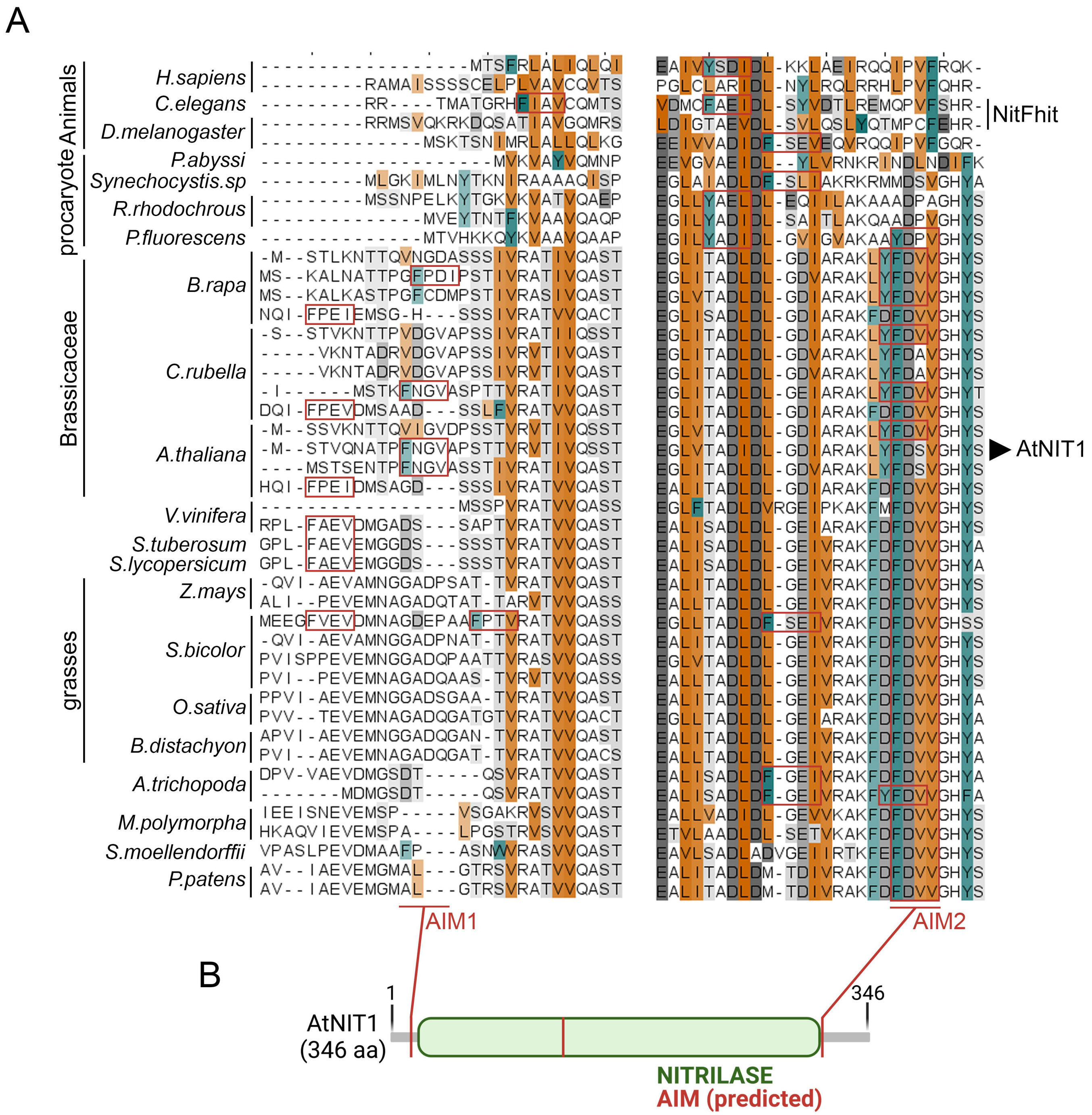
A
B
H.sapiens
C.elegans
D.melanogaster
P.abyssi
Synechocystis.sp
R.rhodochrous
P.fluorescens
B.rapa
C.rubella
A.thaliana
V.vinifera
S.tuberosum
S.lycopersicum
Z.mays
S.bicolor
O.sativa
B.distachyon
A.trichopoda
M.polymorpha
S.moellendorffii
P.patens
Animals
procaryote
Brassicaceae
grasses
NitFhit
AtNIT1
AIM1
AIM2
AtNIT1
(346 aa)
1
346
NITRILASE
AIM (predicted)

**FIGURE S10: AIM positions and conservation in NIT-type proteins (supports Figure 4). (A)** Trimmed multiple sequence alignment of NIT proteins depicting the conservation of the predicted canonical AIMs (iLIR on Arabidopsis NIT1 sequence used as bait) and their position in the aligned proteins. AIMs are boxed in red, W/F/Y (position Θ) residues are underlined in teal, L/I/V residues (position Γ) in orange and S/T (phosphorylatable) and E/D (negatively charged) are underlined in light and dark grey respectively. Color intensity is proportional to conservation (cutoff threshold 10%). Species of origin are indicated on the left. NitFhit: Nit Fhit protein fusion. **(B)** Schematic representation of linear protein domain organization and putative AIM positions within the sequence of the indicated protein. Only AIMs that were experimentally tested are numbered.

FIGURE S11

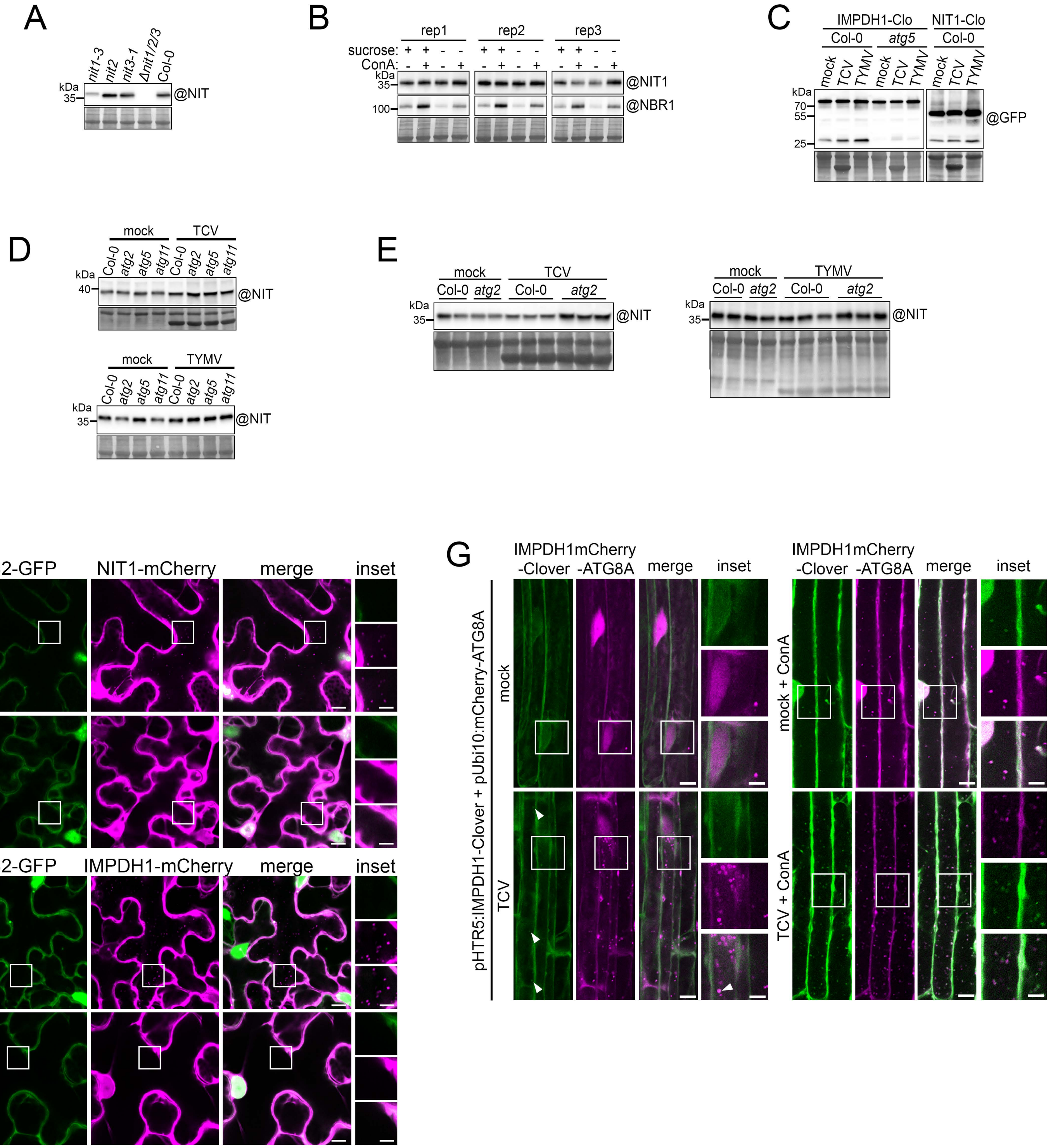
A
kDa
35
nit1-3
nit2
nit3-1
Δnit1/2/3
Col-0
@NIT
B
rep1
rep2
rep3
sucrose:
ConA:
@NIT1
@NBR1
100
C
IMPDH1-Clo
NIT1-Clo
Col-0
atg5
mock
TCV
TYMV
70
55
25
@GFP
D
mock
TCV
TYMV
Col-0
atg2
atg5
atg11
40
@NIT
E
F
B2-GFP
NIT1-mCherry
IMPDH1-mCherry
merge
inset
EV
G
IMPDH1-Clover
mCherry-ATG8A
pHTR5:IMPDH1-Clover + pUbi10:mCherry-ATG8A
mock + ConA
TCV + ConA

**FIGURE S11: NIT and IMPDH are virus-specific selective autophagy receptor *in vivo* (supports Figure 5). (A)** Endogenous NIT antibody detection specificity. Detection of a signal at the expected size for the NITRILASE proteins in Col-0, *nit1-3*, *nit2*, *nit3-1* and *Δnit1/2/3* seedlings. 10 µg of total protein was loaded per well and membranes were hybridized with a commercial NIT1 antibody (@). Membranes were then stained with amidoblack to verify loading. **(B)** Autophagic flux in 8-day-old seedlings treated with or without carbon starvation with or without 1µM ConcanamycinA (ConA) for 24 hours. 15 µg of total protein was loaded per well and membranes were hybridized with the indicated antibodies (@). Membranes were then stained with amidoblack to verify loading. Quantification is provided in Figure 5C. **(C)** Autophagic flux in systemic leaves of plants expressing NITRILASE1-Clover (NIT1-clo, left panel) and IMPDH1-Clover (IMPDH1-clo, right panel) under the HTR5 promoter. Plants were either treated with buffer (mock), TCV or TYMV and samples were collected at 10 dpi. n=10 plants per genotype-treatment combination. 25 µg of total protein was loaded per well and membranes were hybridized with a GFP antibody (@). Membranes were then stained with amidoblack to verify loading. **(D)** Immunoblot of endogenous NIT protein content in mock and TCV (top panel), in mock and TYMV (bottom panel) inoculated systemic leaves of WT, *atg2*, *atg5* and *atg11* plants. 3-week-old rosettes were inoculated with buffer (mock), TCV or TYMV and young systemic leaves were collected at 12 dpi. n=10 plants. Total soluble proteins were extracted at a fixed fresh weight to buffer volume ratio and equal volume was loaded per well. Membranes were hybridized with the NIT antibody (@). Membranes were then stained with amidoblack to verify loading. See Figure 5E and 5F for a biological replicate. **(E)** Immunoblot of endogenous NIT protein content in mock and TCV inoculated systemic leaves of WT and *atg2* plants (top panel) and the same genotypes inoculated with TYMV (bottom panel). 3-week-old rosettes were inoculated with buffer (mock) with TCV or TYMV and young systemic leaves were collected at12 dpi. n=10 plants per sample, each lane is a biological replicate. Total soluble proteins were extracted at a fixed fresh weight to buffer volume ratio and equal volume was loaded per well. Membranes were hybridized with the NIT antibody (@). Membranes were then stained with amidoblack to verify loading. See Figure 5G and H for quantification. **(F)** Confocal microscopy images of *N. Benthamiana* local leaves stably expressing B2-GFP (green) infiltrated with NIT1-mCherry or IMPDH1-mCherry under the 35S promoter and co-infiltrated with pBin TCV (TCV) or the empty vector control (EV) at 3 dpi. Infection in the leaves was verified by clustering of B2-GFP signal in the cytosol and panels are from two consecutive focal planes. Scale bar 10µm and 5µm in inset. **(G)** Confocal microscopy images of *Arabidopsis* root expressing IMPDH1-Clover (green) under the HTR5 promoter and mCherry-ATG8A (magenta) under a ubiquitin 10 promoter in mock-inoculated or TCV-infected conditions at 7 dpi. Left panels: Seedlings were directly mounted in ½ MS for image acquisition. Each image is a maximum intensity projection of a full z-stack. Right panels: Seedlings were immerged in ½ MS with 1µM ConcanamycinA (ConA) for 3h to 4h then directly mounted in ½ MS for image acquisition. Each image is a single snap in the focal plane of the vacuole. Scale bar 10µm and 5µm in inset.

### FIGURE S12

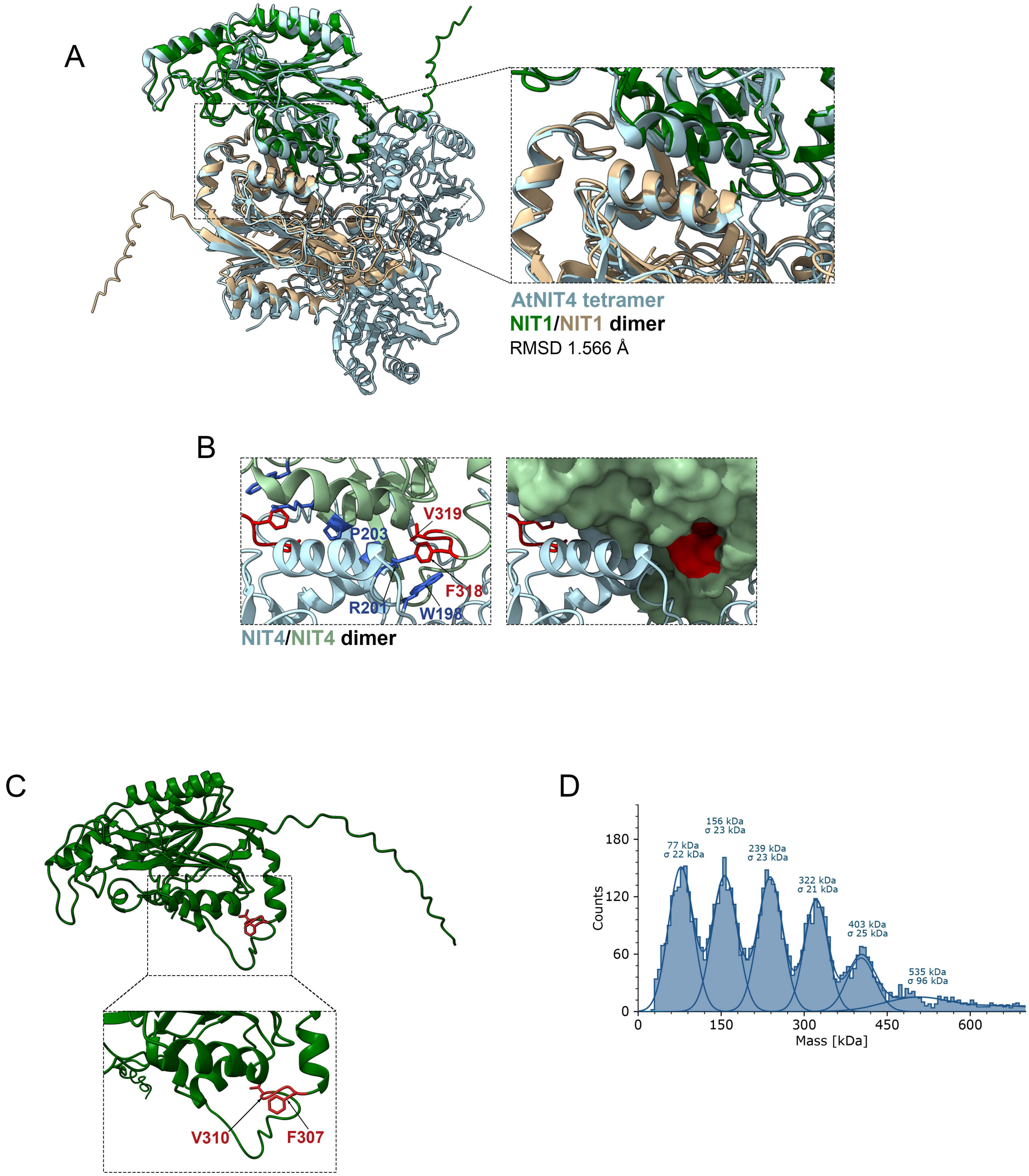
A
AtNIT4 tetramer
NIT1/NIT1 dimer
RMSD 1.566 Å
B
V319
P203
F318
R201
W198
NIT4/NIT4 dimer
C
V310
F307
D
156 kDa
σ 23 kDa
77 kDa
σ 22 kDa
239 kDa
σ 23 kDa
322 kDa
σ 21 kDa
403 kDa
σ 25 kDa
535 kDa
σ 96 kDa
Counts
180
120
60
0
0
150
300
450
600
Mass [kDa]

**FIGURE S12: NIT1 structural prediction of dimerization interface is well supported by NIT4 cryoEM structure (supports Figure 5). (A)** Overlay of NIT1 dimer modeled with AlphaFold2 (in green and tan) and cryoEM structure of Arabidopsis NIT4 filament (6i00, light blue). Only four successive NIT4 protomers are shown. **(B)** NIT4 dimerization interface. NIT4 monomers are shown as ribbon diagrams (left panel) or as a surface representation (right panel). NIT4 monomers are coloured in light green and light blue. AIM residues are highlighted in red and residues corresponding to NIT1 dimerization interface, as shown in Figure 5J are highlighted in blue. **(C)** NIT1 monomer model with AlphaFold. NIT1 is coloured in green and the AIM residues in position Θ (F307) and Γ (V310) are highlighted in red. C-terminal part of the protein is unfolded. **(D)** Mass distribution of NIT1 oligomeric states at 100nM measured with OneMP. NIT1 wt shows several species corresponding to dimer (76 kDa), tetramer (152 kDa), hexamer (228 kDa), octamer (304 kDa) and decamer (380 kDa).

### FIGURE S13

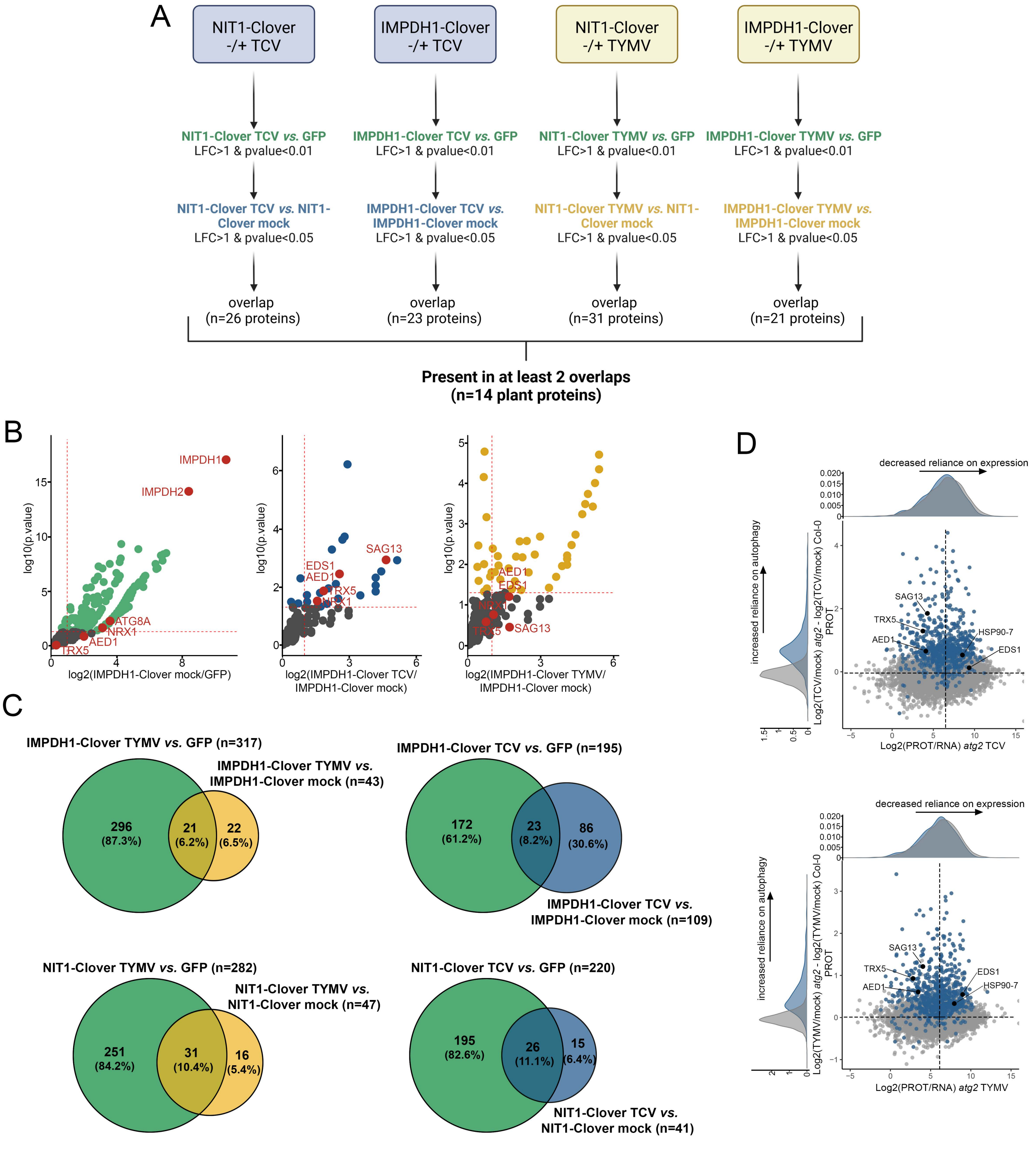

A
NIT1-Clover -/+ TCV
IMPDH1-Clover -/+ TCV
NIT1-Clover -/+ TYMV
IMPDH1-Clover -/+ TYMV
NIT1-Clover TCV vs. GFP
LFC>1 & pvalue<0.01
IMPDH1-Clover TCV vs. GFP
LFC>1 & pvalue<0.01
NIT1-Clover TYMV vs. GFP
LFC>1 & pvalue<0.01
IMPDH1-Clover TYMV vs. GFP
LFC>1 & pvalue<0.01
NIT1-Clover TCV vs. NIT1-Clover mock
LFC>1 & pvalue<0.05
IMPDH1-Clover TCV vs. IMPDH1-Clover mock
LFC>1 & pvalue<0.05
NIT1-Clover TYMV vs. NIT1-Clover mock
LFC>1 & pvalue<0.05
IMPDH1-Clover TYMV vs. IMPDH1-Clover mock
LFC>1 & pvalue<0.05
overlap (n=26 proteins)
overlap (n=23 proteins)
overlap (n=31 proteins)
overlap (n=21 proteins)
Present in at least 2 overlaps (n=14 plant proteins)
B
log10(p.value)
log2(IMPDH1-Clover mock/GFP)
IMPDH1
IMPDH2
ATG8A
NRX1
AED1
TRX5
log2(IMPDH1-Clover TCV/ IMPDH1-Clover mock)
SAG13
EDS1
AED1
TRX5
NRX1
log2(IMPDH1-Clover TYMV/ IMPDH1-Clover mock)
AED1
EDS1
NRX1
TRX5
SAG13
C
IMPDH1-Clover TYMV vs. GFP (n=317)
IMPDH1-Clover TYMV vs. IMPDH1-Clover mock (n=43)
296 (87.3%)
21 (6.2%)
22 (6.5%)
IMPDH1-Clover TCV vs. GFP (n=195)
IMPDH1-Clover TCV vs. IMPDH1-Clover mock (n=109)
172 (61.2%)
23 (8.2%)
86 (30.6%)
NIT1-Clover TYMV vs. GFP (n=282)
NIT1-Clover TYMV vs. NIT1-Clover mock (n=47)
251 (84.2%)
31 (10.4%)
16 (5.4%)
NIT1-Clover TCV vs. GFP (n=220)
NIT1-Clover TCV vs. NIT1-Clover mock (n=41)
195 (82.6%)
26 (11.1%)
15 (6.4%)
D
decreased reliance on expression
increased reliance on autophagy
Log2(TCV/mock) atg2 - log2(TCV/mock) Col-0 PROT
Log2(PROT/RNA) atg2 TCV
SAG13
TRX5
AED1
HSP90-7
EDS1
decreased reliance on expression
increased reliance on autophagy
Log2(TYMV/mock) atg2 - log2(TYMV/mock) Col-0 PROT
Log2(PROT/RNA) atg2 TYMV
SAG13
TRX5
AED1
EDS1
HSP90-7

**FIGURE S13: Identification of NIT1 and IMPDH1 virus-specific cargo proteins (supports Figure 5). (A)** Selection scheme employed for selecting the final list of candidate Virus-specific NIT1-Clover and/or IMPDH1-Clover interacting partners. Proteins were considered as final candidates if they were present at the overlap between two pairwise comparisons for at least two out of four datasets. **(B)** Enrichment of proteins co-purified with IMPDH1-Clover represented by a volcano plot. Left panel: pairwise comparison of IMPDH1-Clover mock libraries to GFP control (all treatments) to reveal constitutive IMPDH1-Clover interaction partners. Middle panel: pairwise comparison of IMPDH1-Clover treated with TCV to IMPDH1-Clover mock treated to reveal TCV-inducible interacting partners of IMPDH1-Clover. Right panel: pairwise comparison of IMPDH1-Clover treated with TYMV to IMPDH1-Clover mock treated to reveal TYMV-inducible interacting partners of IMPDH1-Clover. The horizontal dashed line indicates the threshold above which proteins are significantly enriched ($p$ value < 0.05, quasi-likelihood negative binomial generalized log-linear model) and the vertical dashed line the threshold for which proteins log2 fold change is above 1. **(C)** Venn diagram of two overlapping pairwise comparisons for AP-MS conducted in NIT1-Clover and IMPDH1-Clover plants infected with TCV or TYMV: Bait *vs.* GFP control; green circles for the respective bait and treatment. TYMV-infected bait *vs.* mock-inoculated bait; yellow circles. TCV-infected bait *vs.* mock-inoculated bait; blue circles. **(D)** Graphical representation of the ratio of protein abundance to RNA abundance in *atg2* infected samples (log2(protein abundance/transcript abundance) as captured by quantitative proteomics and transcriptomic) on the x axis versus the ratio of protein abundance in *atg2*-infected samples with the removed contribution of the WT Col-0 background (log2 infected/mock) in *atg2* – log2(infected/mock) in Col-0) as captured by quantitative proteomics on the y axis. Top panel: TCV, bottom panel: TYMV. Quadrants are determined using centroids for all the plotted datapoints. Data distribution for both axis is plotted next to the relevant axis. Proteins mapped as virus-specific autophagy cargos are colored in blue, while the five candidate proteins are highlighted in black.

### FIGURE S14

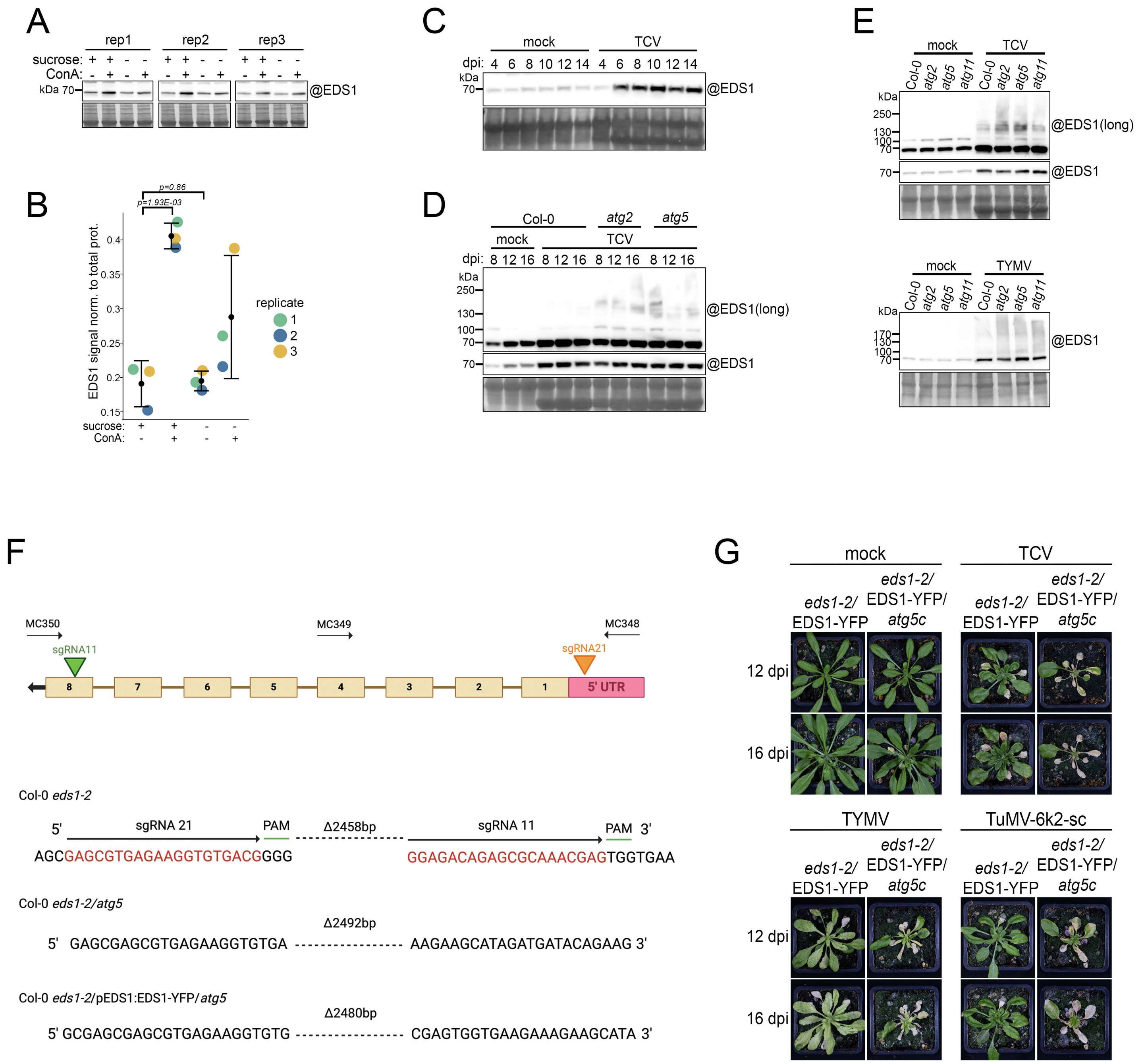

A
rep1
rep2
rep3
sucrose: + + - - + + - - + + - -
ConA: - + - + - + - + - + - +
kDa 70
@EDS1
B
p=0.86
p=1.93E-03
EDS1 signal norm. to total prot.
0.4
0.35
0.3
0.25
0.2
0.15
sucrose: + + - -
ConA: - + - +
replicate
1
2
3
C
mock
TCV
dpi: 4 6 8 10 12 14 4 6 8 10 12 14
kDa
70
@EDS1
D
Col-0
atg2
atg5
mock
TCV
dpi: 8 12 16 8 12 16 8 12 16 8 12 16
kDa
250
130
100
70
@EDS1(long)
70
@EDS1
E
mock
TCV
Col-0 atg2 atg5 atg11 Col-0 atg2 atg5 atg11
kDa
250
130
100
70
@EDS1(long)
70
@EDS1
mock
TYMV
Col-0 atg2 atg5 atg11 Col-0 atg2 atg5 atg11
kDa
170
130
100
70
@EDS1
F
MC350
MC349
MC348
sgRNA11
sgRNA21
8 7 6 5 4 3 2 1
5' UTR
Col-0 eds1-2
5'
sgRNA 21
PAM
Δ2458bp
sgRNA 11
PAM
3'
AGCGAGCGTGAGAAGGTGTGACGGGG
GGAGACAGAGCGCAAACGAGTGGTGAA
Col-0 eds1-2/atg5
Δ2492bp
5' GAGCGAGCGTGAGAAGGTGTGA ------------------ AAGAAGCATAGATGATACAGAAG 3'
Col-0 eds1-2/pEDS1:EDS1-YFP/atg5
Δ2480bp
5' GCGAGCGAGCGTGAGAAGGTGTG ------------------ CGAGTGGTGAAGAAAGAAGCATA 3'
G
mock
TCV
eds1-2/ EDS1-YFP
eds1-2/ EDS1-YFP/ atg5c
12 dpi
16 dpi
TYMV
TuMV-6k2-sc
eds1-2/ EDS1-YFP
eds1-2/ EDS1-YFP/ atg5c
12 dpi
16 dpi

**FIGURE S14: EDS1 is a virus-specific cargo of NIT1 and IMPDH1 (supports Figure 5). (A)** Autophagic flux in 8-day-old seedlings treated with or without carbon starvation with or without 1µM ConcanamycinA (ConA) for 24 hours. 15 µg of total protein was loaded per well and membranes were hybridized with an anti EDS1 antibody (@). Membranes were then stained with amidoblack to verify loading. **(B)** Quantification of endogenous EDS1 as presented in (A). Signal intensity is normalized to total protein signal. **(C)** Immunoblot of EDS1 endogenous protein in mock inoculated and TCV infected systemic leaves. 3-week-old rosettes were inoculated with buffer (mock) or with TCV and young systemic leaves were collected at the indicated time points. n=3 to 4 plants per sample-timepoint combination. Total soluble proteins were extracted at a fixed fresh weight to buffer volume ratio and equal volume was loaded per well. Membranes were hybridized with an anti EDS1 antibody (@). Membranes were then stained with amidoblack to verify loading. **(D)** Immunoblot of EDS1 endogenous protein and high molecular weight signal in mock and TCV infected systemic leaves of WT, *atg2* and *atg5* plants. 3-week-old rosettes were inoculated with buffer (mock) or with TCV and young systemic leaves were collected at the indicated time points. n=10 plants per sample-timepoint combination. Total soluble proteins were extracted at a fixed fresh weight to buffer volume ratio and equal volume was loaded per well. Membranes were hybridized with an anti EDS1 antibody (@). Membranes were then stained with amidoblack to verify loading. **(E)** Immunoblot of endogenous EDS1 protein content in mock, TCV (top panel) and TYMV (bottom panel) inoculated systemic leaves of WT, *atg2*, *atg5* and *atg11* plants. 3-week-old rosettes were inoculated with buffer (mock), with TCV or TYMV and young systemic leaves were collected at 12 dpi. n=10 plants. Total soluble proteins were extracted at a fixed fresh weight to buffer volume ratio and equal volume was loaded per well. Membranes were hybridized with the EDS1 antibody (@). Membranes were then stained with amidoblack to verify loading. See Figure 5P for a biological replicate. **(F)** Top panel: Schematic representation of the AT5G17290 locus with sequencing oligonucleotides and sgRNA targets indicated. Colored arrows indicate designated sgRNA target sites. Each block represents one of the eight exons of the AT5G17290 gene, while the pink box represents the 5' UTR region. Bottom panel: Col-0 *eds1* sequence reference with sgRNAs used annotated. CRISPR/Cas9 deletions obtained in AT5G17290 locus for Col-0 *eds1/atg5c* and Col-0 *eds1*/pEDS1:EDS1-YFP/*atg5c* mutants according to Sanger DNA sequencing results. **(G)** Phenotypic characterization of *Arabidopsis eds1*/EDS1-YFP and *eds1*/EDS1-YFP/*atg5* crispr (*atg5c*) mutant plants after inoculation with buffer (mock), with TCV, TYMV and TuMV-6k2-sc, 12 and 16 days-post-inoculation (dpi). See Figure 5Q for additional genotypes.

#### Supplemental videos

**Video S1**: Distribution of pixel hue values extracted form plants inoculated with buffer (mock) from -10 dpi to +26 dpi. Scale bar represents hue values for the full colour spectrum, values around 100 (green) indicate healthy tissue whereas a shift towards 0 (yellow-orange) indicates progression of viral infection symptoms. Values were obtained from the same dataset as Figure 1B.

**Video S2**: Distribution of pixel hue values extracted form plants inoculated with TCV from -10 dpi to +26 dpi. Scale bar represents hue values for the full colour spectrum, values around 100 (green) indicate healthy tissue whereas a shift towards 0 (yellow-orange) indicates progression of viral infection symptoms. Values were obtained from the same dataset as Figure 1B.

**Video S3**: Distribution of pixel hue values extracted form plants inoculated with TuMV-6k2-Scarlet from -10 dpi to +26 dpi. Scale bar represents hue values for the full colour spectrum, values around 100 (green) indicate healthy tissue whereas a shift towards 0 (yellow-orange) indicates progression of viral infection symptoms. Values were obtained from the same dataset as Figure 1B.

**Video S4**: Distribution of pixel hue values extracted form plants inoculated with TYMV from -10 dpi to +26 dpi. Scale bar represents hue values for the full colour spectrum, values around 100 (green) indicate healthy tissue whereas a shift towards 0 (yellow-orange) indicates progression of viral infection symptoms. Values were obtained from the same dataset as Figure 1B.

**Video S5**: Confocal microscopy timelapse of a TCV replication complex in *Arabidopsis* root expressing B2-GFP (green) under the 35S promoter and stained with TMRE (magenta) at 5 dpi. Seedlings were immerged in ½ MS with 500nM of TMRE for a few minutes then mounted in ½ MS for image acquisition. 10 seconds frame interval.

**Video S6:** Transmission electron tomogram series of a mitochondrion in root cells expressing P29 and three-dimensional model of the mitochondrion. P29-induced small vesicles are in red and their associated cristae in green. Vesicle-free cristae are rendered in blue and outer mitochondrial membrane in white.

**Video S7:** Confocal microscopy timelapse an autophagosome (mCherry-ATG8A) moving along with IDH-GFP puncta in the cytosol in TCV-infected roots.

#### Supplemental tables

**Table S1:** **(A)** PRM peptides detected in TCV infected samples. **(B)** PRM peptides detected in TYMV infected samples. **(C)** PRM peptides detected in TuMV-6k2-Scarlet infected samples.

**Table S2:** Normalized peptide areas for PRM experiment in TCV infected samples.

**Table S3:** Normalized peptide areas for PRM experiment in TYMV infected samples.

**Table S4:** Normalized peptide areas for PRM experiment in TuMV-6k2-Scarlet infected samples.

**Table S5:** **(A)** Differential protein enrichment in mCherry-ATG8E (AIM *mut*) compared to mCherry control line. **(B)** Differential protein enrichment in mCherry-ATG8E (AIM *mut*) compared to mCherry-ATG8E (AIM *wt*). **(C)** Differential protein enrichment in mCherry-ATG8E (AIM *mut*) TCV infected compared to mCherry-ATG8E (AIM *mut*) mock-inoculated.

**Table S6:** **(A)** Differential protein enrichment in mCherry-ATG8A (AIM *mut*) compared to mCherry control line. **(B)** Differential protein enrichment in mCherry-ATG8A (AIM *mut*) compared to mCherry-ATG8A (AIM *wt*). **(C)** Differential protein enrichment in mCherry-ATG8A (AIM *mut*) TYMV infected compared to mCherry-ATG8A (AIM *mut*) mock-inoculated. **(D)** Differential protein enrichment in mCherry-ATG8E (AIM *mut*) compared to mCherry control line. **(E)** Differential protein enrichment in mCherry-ATG8E (AIM *mut*) compared to mCherry-ATG8E (AIM *wt*). **(F)** Differential protein enrichment in mCherry-ATG8E (AIM *mut*) TYMV infected compared to mCherry-ATG8E (AIM *mut*) mock-inoculated.

**Table S7:** **(A)** Differential protein enrichment in mCherry-ATG8A (AIM *mut*) compared to mCherry control line. **(B)** Differential protein enrichment in mCherry-ATG8A (AIM *mut*) compared to mCherry-ATG8A (AIM *wt*). **(C)** Differential protein enrichment in mCherry-ATG8A (AIM *mut*) TCV infected compared to mCherry-ATG8A (AIM *mut*) mock-inoculated.

**Table S8:** Quantitative proteomic cluster results. Detected proteins are either not clustered (NA), or part of cluster 1 to 20.

**Table S9:** DESeq2 Differential gene expression analysis Col-0 TCV *vs.* Col-0 mock

**Table S10:** DESeq2 Differential gene expression analysis Col-0 TYMV *vs.* Col-0 mock

**Table S11:** DESeq2 Differential gene expression analysis *atg2* TCV *vs. atg2* mock

**Table S12:** DESeq2 Differential gene expression analysis *atg2* TYMV *vs. atg2* mock

**Table S13:** **(A)** Differential protein enrichment in NIT1-Clover mock inoculated compared to GFP control. **(B)** Differential protein enrichment in NIT1-Clover TCV inoculated compared to GFP control. **(C)** Differential protein enrichment in NIT1-Clover TYMV inoculated compared to GFP control. **(D)** Differential protein enrichment in NIT1-Clover TCV infected compared to NIT1-Clover mock inoculated. **(E)** Differential protein enrichment in NIT1-Clover TYMV infected compared to NIT1-Clover mock inoculated.

**Table S14:** **(A)** Differential protein enrichment in IMPDH1-Clover mock inoculated compared to GFP control. **(B)** Differential protein enrichment in IMPDH1-Clover TCV infected compared to GFP control. **(C)** Differential protein enrichment in IMPDH1-Clover TYMV infected compared to GFP control. **(D)** Differential protein enrichment in IMPDH1-Clover TCV infected compared to IMPDH1-Clover mock inoculated. **(E)** Differential protein enrichment in IMPDH1-Clover TYMV infected compared to IMPDH1-Clover mock inoculated.

**Table S15:** Plant lines used in this study.

**Table S16:** Oligonucleotides used in this study.

**Table S17:** DNA constructs used in this study.

**Table S18:** Viral strains and infectious clones used in this study.

**Table S19:** Key reagents used in this study.

**Table S20:** Normalized protein areas, coverage details, peptide isoforms, PSM details and quality control for proteins detected in mCherry-ATG8E, TCV infected samples.

**Table S21:** Normalized protein areas, coverage details, peptide isoforms, PSM details and quality control for proteins detected in mCherry-ATG8E and mCherry-ATG8A TYMV infected samples.

**Table S22:** Normalized protein areas, coverage details, peptide isoforms, PSM details and quality control for proteins detected in mCherry-ATG8A, TCV infected samples.

**Table S23:** Normalized protein areas, coverage details, peptide isoforms, PSM details and quality control for proteins detected in NIT-Clover and IMPDH1-Clover TCV and TYMV-infected samples.

**Table S24:** **(A)** Data matrix of Peptide-Spectrum Match (PSM) for proteins identified in mCherry-ATG8E, TCV infected samples. **(B)** Sample table and treatment table for IPinquiry4 analysis. **(C)** TAIR10 annotations for each detected protein in mCherry-ATG8E TCV infected samples.

**Table S25:** **(A)** Data matrix of Peptide-Spectrum Match (PSM) for proteins identified in mCherry-ATG8E and mCherry-ATG8A TYMV infected sample. **(B)** Sample table and treatment table for IPinquiry4 analysis. **(C)** TAIR10 annotations for each detected protein in mCherry-ATG8E and mCherry-ATG8A TYMV infected samples.

**Table S26:** **(A)** Data matrix of Peptide-Spectrum Match (PSM) for proteins identified in mCherry-ATG8A, TCV infected samples. **(B)** Sample table and treatment table for IPinquiry4 analysis.

**Table S27:** **(A)** Data matrix of Peptide-Spectrum Match (PSM) for proteins identified in NIT-Clover and IMPDH1-Clover, TCV and TYMV infected samples. **(B)** Sample table and treatment table for IPinquiry4 analysis. **(C)** TAIR10 annotations for each detected protein in NIT-Clover and IMPDH1-Clover TCV and TYMV-infected samples

**Table S28:** All detected proteins in quantitative proteomic approach. Normalized protein abundance was extracted for k-means clustering
